# MX2 restricts HIV-1 and herpes simplex virus type 1 by forming cytoplasmic biomolecular condensates that mimic nuclear pore complexes

**DOI:** 10.1101/2023.06.22.545931

**Authors:** George D. Moschonas, Louis Delhaye, Robin Cooreman, Franziska Hüsers, Anayat Bhat, Delphine De Sutter, Eef Parthoens, Anne-Sophie Desmet, Aleksandra Maciejczuk, Hanna Grzesik, Saskia Lippens, Zeger Debyser, Beate Sodeik, Sven Eyckerman, Xavier Saelens

## Abstract

Human myxovirus resistance 2 (MX2) can potently restrict HIV-1 and herpesviruses at a post-entry step by a process that requires MX2 interaction with the capsids of these viruses. The involvement of other host cell factors in this process, however, remains poorly understood. Here, we mapped the proximity interactome of MX2 revealing strong enrichment of phenylalanine-glycine (FG)-rich proteins related to the nuclear pore complex as well as proteins that are part of cytoplasmic ribonucleoprotein granules. MX2 interacted with these proteins to form multiprotein cytoplasmic biomolecular condensates that were essential for its anti-HIV-1 and -herpes simplex virus-1 (HSV-1) activity. MX2 condensate formation required the disordered N-terminal region of MX2 and its dimerization. Incoming HIV-1 and HSV-1 capsids associated with MX2 at these dynamic cytoplasmic biomolecular condensates. Our results demonstrate that MX2 forms cytoplasmic condensates that act as nuclear pore decoys, which trap capsids and induce premature viral genome release, and thereby interfere with nuclear targeting of HIV-1 and HSV-1.

## Introduction

Vertebrate cells respond to type I interferon (IFN) by the induction of hundreds of IFN-stimulated genes (ISGs), which establishes an “antiviral state” in the cell^1,2^. Humans, like most mammals, have two paralogues of the potent *Myxovirus Resistance* (*MX)* ISGs named *MX1* and *MX2* and their gene products, named MX1 (or MxA) and MX2 (or MxB), can restrict a broad range of viruses^3^. MX2 has two isoforms: full-length MX2(1-715) and a short isoform MX2(26-715) lacking the first 25 N-terminal residues^4^.

Human MX2 can restrict HIV-1 and other primate lentiviruses^5–7^. More recently, it was reported that MX2 can also suppress the replication of herpesviruses^8–13^, Hepatitis C Virus (HCV)^14,15^, and Hepatitis B Virus (HBV)^16^. In addition, MX2 is implicated in mitochondrial integrity^17^ and in the control of long interspersed element type 1 retrotransposition^18^.

MX proteins belong to the dynamin superfamily of large GTPases^19^. The crystal structure of the MX2 dimer, lacking the first N-terminal 83 residues, revealed a three-domain architecture with an N-terminal globular GTPase domain connected to a stalk domain by a bundle signaling element^20^. The GTPase domain enables GTP-binding and -hydrolysis, which, remarkably, is required for MX2 antiviral activity against herpesviruses and HBV but not for HIV-1 restriction^5,6,9,11,13,21^. MX2 oligomerization, at least as a dimeric assembly, and localization are major determinants of MX2 activity against HIV-1, herpesviruses, and HBV^9,13,16,20,22,23^.

The N-terminal domain (NTD) of MX2 is also a key determinant for MX2 antiviral activity^24^. The first 25 residues contain a nuclear localization signal (NLS) that targets MX2 to the nuclear membrane and a triple arginine motif that facilitates HIV-1 capsid binding^21,25,26^. The NTD also harbors a disordered region of residues 26-90 that is crucial for MX2 localization and HIV-1 and HBV antiviral activity^16,21^. In contrast to HIV-1, restricting HSV-1 depends on the NTD, not on the arginine motif, but on the methionine at position 83^8,9,13,27^.

Proteins with disordered regions are prone to undergo so-called liquid-liquid phase separation (LLPS), a compartmentalization process that is associated with the formation of biomolecular condensates and membraneless organelles (MLOs) in cells^28–30^. Biomolecular condensates are subcellular compartments of highly concentrated proteins and, often, nucleic acids, which can rapidly exchange material with their surroundings, dissipate, and reassemble^31,32^. So-called client proteins preferentially partition into condensates that have been formed by condensate-initiating drivers^29^.

Here we used an optimized proximal labeling method to identify MX2 interaction partners, revealing SAMD4A, TNPO1, and NUP35 as crucial MX2 partner proteins in HIV-1 and HSV-1 restriction. Interaction with MX2 occurs in multiprotein cytoplasmic and perinuclear biomolecular condensates, which are essential for MX2-mediated HIV-1 and HSV-1 restriction. MX2 condensates engage with incoming HIV-1 and HSV-1 capsids, acting as decoys for the nuclear pore, and prevent nuclear entry of incoming viral genomes.

## Results

### MX2 interacts with FG-rich nucleoporins and granule-associated ribonucleoproteins

To identify MX2 protein partners, we performed proximity-dependent biotin identification (BioID) using TurboID, a highly active promiscuous biotin ligase^33,34^. MX2(1-715) was genetically fused with TurboID through a T2A self-cleaving site for control conditions (MX2 and TurboID expressed as separate proteins), and through a mutant T2A (MUTT2A) linker for the experimental condition (MX2 fused to TurboID)^35^ (Figure S1A). MX2-MUTT2A-TurboID maintained capacity to dimerize and co-localize with co-expressed MX2(1-715), its ability to restrict HSV-1 and HIV-1, and its localization at the nuclear envelope (Figure S1B, S1C, S1D, S1E, S1F and S1G).

Next, we generated two doxycycline-inducible U87-MG cell lines which expressed either MX2 and TurboID separately (MX2-T2A-TurboID) or as fused (MX2-MUTT2A-TurboID) proteins, together with untagged MX2 that can support the physiological function of the fused construct (Figure S1B, S1D and S1G). This ”T2A split-link design” ensures equal biotin ligase activity in both cell lines (Figure S1H)^36^. We performed BioID experiments with these cell line pairs under three conditions: (i) doxycycline-treated (-IFN), (ii) doxycycline-treated combined with IFNa2a stimulation (+IFN), and (iii) doxycycline-treated and infected with HSV-1 (HSV-1) (Figure S1I). Our T2A split-link design resulted in comparable MX2 differential enrichment in all three conditions (Figure S1J).

We identified nearly all previously reported MX2-interacting proteins except RUNX3, PNRC1 and KLHL6^37^, and PPP1CB^38^. We also found novel, highly enriched, MX2-interacting candidates known to localize at the nuclear pore or in the cytoplasm (Figure 1A and Table S1). There were 253 proteins enriched in all three conditions (Figure 1B and Table S2). A subset of these proteins participate in nuclear transport and are part of or interact with the nuclear pore complex (NPC). This nuclear pore network includes novel MX2-interacting candidates such as NUPL1(NUP58), NUP54, NUP35, and nuclear transport receptor transportin-2 (TNPO2), which were highly enriched in the interactome screen. Unexpectedly, a large fraction (106 proteins) of the common enriched proteins participate in the regulation of translation and RNA catabolic processes, and are part of cytosolic ribonucleoprotein (RNP) granules, cellular bodies which have been proposed to form via intracellular phase transitions (Figure 1B and S2A). Other common candidate MX2-interacting proteins included mitochondrial and coated vesicle proteins (Figure S2B, S2C and Table S2).

**Figure 1:**
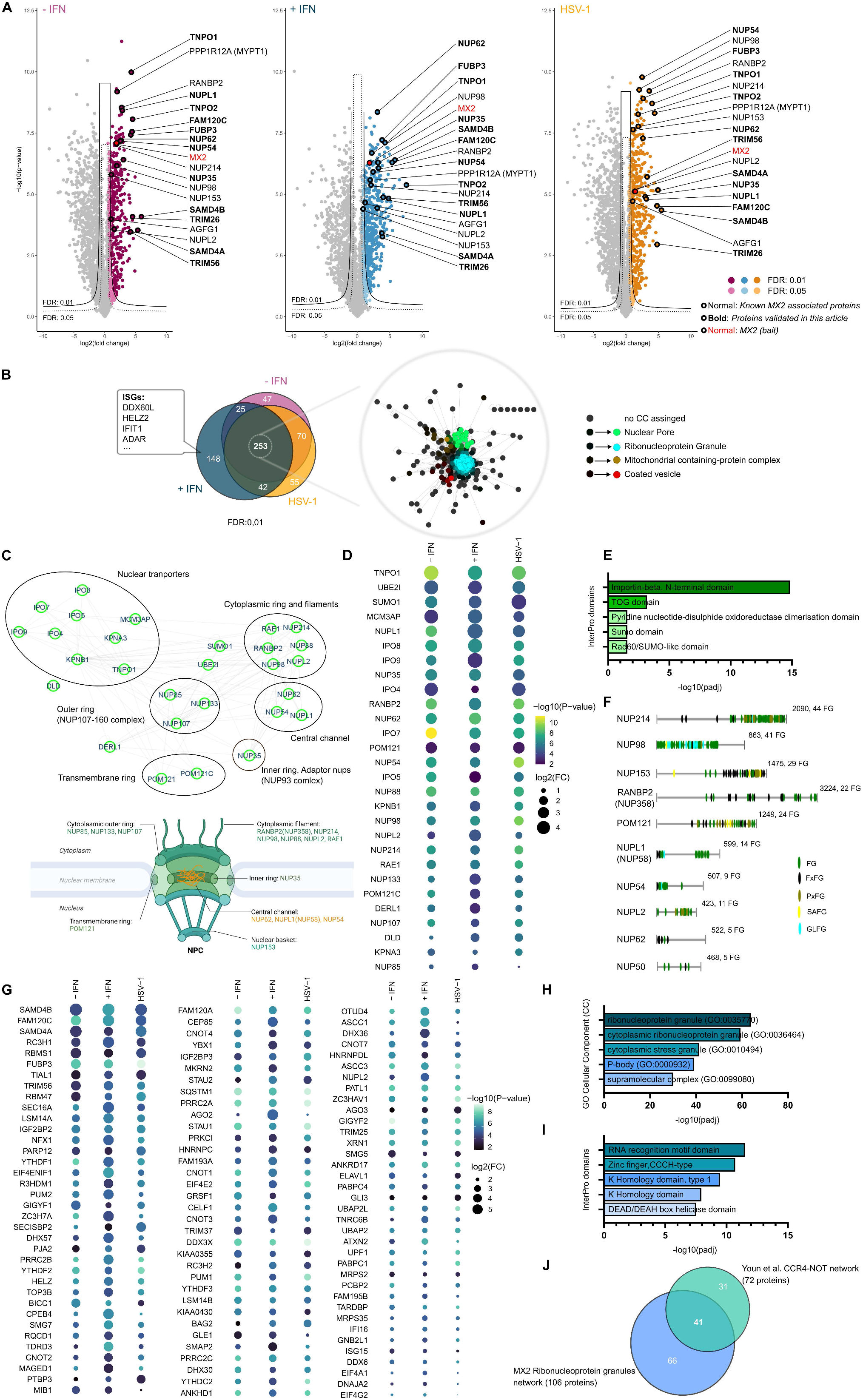
MX2 associates with nuclear transport proteins and cytoplasmic granule-associated ribonucleoproteins. **(A)** Volcano plots of the three MX2 BioID conditions (-IFN, +IFN, HSV-1). Specific MX2 interaction partners outside the FDR 0.05 and 0.01 curved lines are annotated by gene name. Reported MX2 associated partners are indicated with normal, while novel candidate protein partners researched in this study are indicated with bold font. Bait protein MX2 is indicated in red. **(B)** Left: Venn diagram of MX2 associated proteins (FDR 0.01) for the three BioID conditions, indicating ISGs only identified in the +IFN condition. Right: SAFE visualization of the MX2 network. Each node(protein) in the network is colored according to its association to a specific Gene Ontology (GO) cellular component term. **(C)** Top: SAFE enriched ‘’nuclear pore’’ domain protein network, clustered in specific protein sub-clusters and subcomplexes. Bottom: The NPC according to^94^, created with BioRender software. **(D)** Dotplot enrichment analysis of nuclear pore domain proteins. **(E)** Bar chart showing the top 5 InterPro analysis predicted domains for the nuclear pore domain proteins. **(F)** Domain architecture of disordered human FG-rich Nups according to UniProt. **(G)** Dotplot enrichment analysis of of granule ribonucleoprotein domain proteins. **(H, I)** Bar chart showing the top 5 (**H)** GO cellular components terms, (**I)** InterPro analysis predicted domains, for the proteins of the granule ribonucleoprotein network. **(J)** Venn diagram of the granule ribonucleoprotein network and the Youn *et al*.^45^ PB subnetwork.

The nuclear pore network consists of nucleoporins (Nups), which are structural components of the NPC, and nuclear transport factors (Figure 1C). The enriched nuclear transport factors were primarily members of the importin-beta family, while importins-alpha, their partners to mediate nuclear cargo translocation, were poorly enriched and not consistently identified (Figure 1C, 1D and 1E). MX2 is localized at the cytoplasmic face of the NPC^37^. The cytoplasmic outer ring components NUP133, NUP85, and NUP107, however, were the least enriched Nups, while Nups that form part of the central channel (e.g. NUPL1), cytoplasmic filaments (e.g. NUP358/RANBP2), inner ring (e.g. NUP35), and even the transmembrane ring of the NPC (e.g. POM121) were highly enriched (Figure 1C and 1D). These highly enriched Nups comprised phenylalanine-glycine (FG)-rich Nups of the human NPC^39^ (Figure 1D and 1F). This was a striking observation because FG-Nups can assemble into hydrogels and biomolecular condensates, a property shared with inner ring Nup NUP35^40–43^.

The ribonucleoprotein granule network consists of RNA-binding and disordered domain-containing proteins that are present in cytoplasmic MLOs, specifically in processing bodies (PBs) and stress granules (SGs) (Figure 1G, 1H, 1I and S2D). SAMD4B, FAM120C, SAMD4A, FUBP3, and TRIM56 were among the more enriched of these ribonucleoprotein granule network proteins (Figure 1G). For example, SAMD4A (also known as Smaug1) forms granular cytoplasmic LLPS condensates that are distinct from PBs and SGs^44^. This MX2-interaction network overlapped substantially with a reported PB subnetwork^45^ (Figure 1J and Table S3). Our proximity interactome analysis suggests that MX2 interacts with FG-rich NPC components and with cytoplasmic granule-associated ribonucleoproteins.

### HIV-1 and HSV-1 inhibition by MX2 involves SAMD4A, TNPO1, and NUP35

We performed RNA interference (RNAi) knock down (KD) experiments to evaluate functional relevance of candidate MX2-interacting proteins on MX2-mediated restriction of HIV-1 and HSV-1 using doxycycline-inducible MX2 HeLa(MX2) cells. We selected 12 genes based on enrichment in our screening and/or their reported roles for nuclear targeting of incoming HIV-1 or HSV-1 capsids^46,47^ (Figure S3A). KD of SAMD4A, TNPO1, or NUP35 impaired MX2-mediated restriction of HSV-1 and HIV-1. KD of NUP62, a central channel Nup interacting with HIV-1 but not HSV-1 capsids, only affected HIV-1 restriction^48,49^ (Figure 2A and 2B). The observations for TNPO1, SAMD4A, NUP35, and NUP62 were confirmed in HeLa(MX2) and U87-MG(MX2) by multiple round infection with an HSV-1 C12 GFP reporter strain and single round infection experiments with HSV-1 and HIV-1. We did not detect any synergistic effects by simultaneous KD of two genes (Figure 2C, 2D, 2E, S3B, S3C and S3D). In U87-MG(MX2) cells, however, KD of SAMD4A and NUP35 did not affect HIV-1 restriction (Figure 2E). HSV-1 and HIV-1 restriction by endogenous MX2 upon IFN stimulation depended on TNPO1, SAMD4A, and NUP35 in U87-MG and HeLa cells (Figure 2F, 2G, 2H, S3E and S3F). Thus, TNPO1, SAMD4A, and NUP35 play a critical role in MX2-mediated restriction of both HIV-1 and HSV-1, while NUP62 KD only affects HIV-1 restriction.

**Figure 2:**
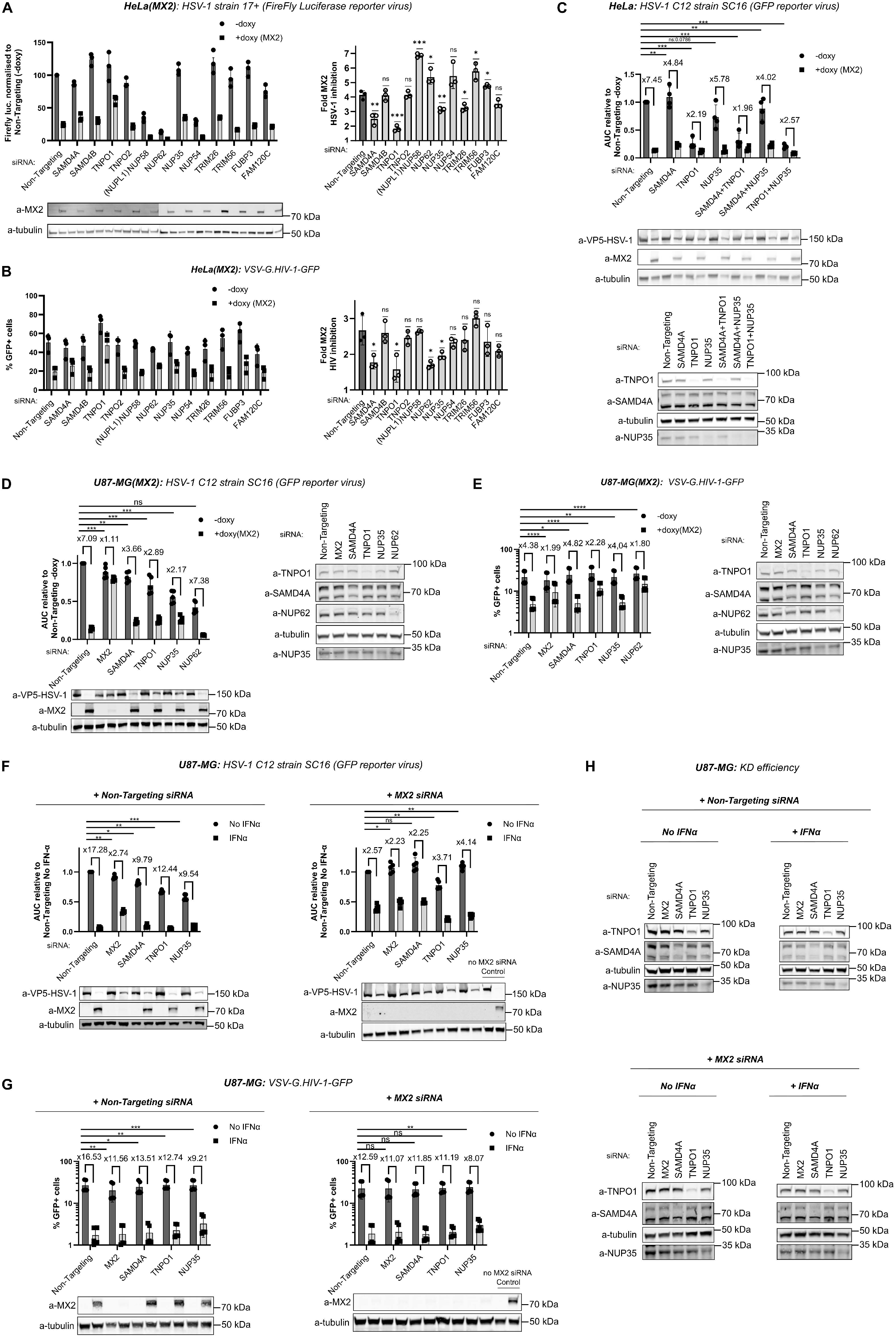
SAMD4A, TNPO1, and NUP35 are required for MX2-mediated antiviral activity against HIV-1 and HSV-1, whereas NUP62 is required for HIV-1 only. **(A, B, C)** Doxycycline-inducible HeLa(MX2) cells were transfected with siRNA and left untreated or treated with 100 ng mL^-1^ doxycycline. **(A)** Left: 48 h after transfection, cells were infected with HSV-1 firely reporter virus (MOI:1). Luminescence is shown relative to the Non-Targeting (-doxy) condition. Right: Fold MX2-mediated HSV-1 inhibition calculated by dividing luminescence of untreated by doxycycline-induced samples in each condition. **(B)** Left: 48 h after transfection, cells were inoculated with VSV-G.HIV-1-GFP (MOI:0.4). Right: Fold MX2-mediated HIV-1 inhibition calculated by dividing % GFP+ cells of untreated by doxycycline-induced samples in each condition. **(A, B)** (mean ± s.d., n = 3 independent experiments, two-tailed unpaired *t*-test to Non-Targeting siRNA; *p* values indicated with \**p* < 0.05, \*\**p* < 0.01, \*\*\**p* < 0.001). **(C)** Twenty-four hours after transfection, cells were inoculated with HSV-1 strain C12 (MOI:0.05). Viral growth area under the curve (AUC) are relative to Non-Targeting (-doxy) samples which are each set to 1. Ratio of AUC of untreated divided by the doxycycline-induced samples in each condition is shown (mean ± s.d., n = 4 independent experiments, two-tailed paired *t*-test to Non-Targeting siRNA; *p* values indicated with \**p* < 0.05, \*\**p* < 0.01, \*\*\**p* < 0.001). **(D, E)** Doxycycline-inducible U87-MG(MX2) cells were transfected twice with siRNA and left untreated or treated with 100 ng mL^-1^ doxycycline. **(D)** Twenty-four hours after transfection, cells were inoculated with HSV-1 strain C12 (MOI:0.05). Viral growth area under the curve (AUC) are relative to Non-Targeting (-doxy) samples which are each set to 1. Ratio of AUC between untreated and the doxycycline-induced samples in each condition is shown. **(E)** Forty-eight hours after transfection, cells were inoculated with VSV-G.HIV-1-GFP (MOI:0.2). Ratio of % GFP+ cells between untreated and the doxycycline-induced samples in each condition is shown. (**D, E)** (mean ± s.d., n = 5 independent experiments, two-tailed paired *t*-test to Non-Targeting siRNA; *p* values indicated with \**p* < 0.05, \*\**p* < 0.01, \*\*\**p* < 0.001). **(F, G)** U87-MG were transfected twice with siRNAs targeting the indicated genes together with either Non-Targeting or MX2 siRNAs and left untreated or treated with 1,000 IU ml^−1^ of rhIFN-a2a. **(F)** Twenty-four hours after transfection, cells were inoculated with HSV-1 strain C12 (MOI:0.05). Viral replication was determined similar to Figure 2D. **(G)** Twenty-four hours after transfection, cells were inoculated with VSV-G.HIV-1-GFP (MOI:0.2). Transduction efficiency was assessed similar to Figure 2E. **(A-H)** Immunoblots were used to analyse protein expression using antibodies specific for the indicated proteins.

### MX2 interacts with SAMD4A, TNPO1, and NUP35 at dynamic cytoplasmic biomolecular condensates

We next used high-resolution confocal fluorescent microscopy to assess the subcellular localization of the MX2 interaction partners. As reported before, endogenous and ectopically expressed MX2 localized at the nuclear membrane and in cytoplasmic puncta^5,6,21,26,50,51^ (Figure 3A). We confirmed the interaction of MX2 with SAMD4A, TNPO1, and NUP35 using a luciferase-based protein complementation assay (Figure 3B). SAMD4A was localized in both MX2-containing puncta while still forming distinct SAMD4A-containing condensates^52^. MX2 changed the diffuse cytoplasmic distribution of TNPO1 and NUP35, with both colocalizing with MX2 at the nuclear envelope and in cytoplasmic MX2-containing puncta (Figure 3C).

**Figure 3:**
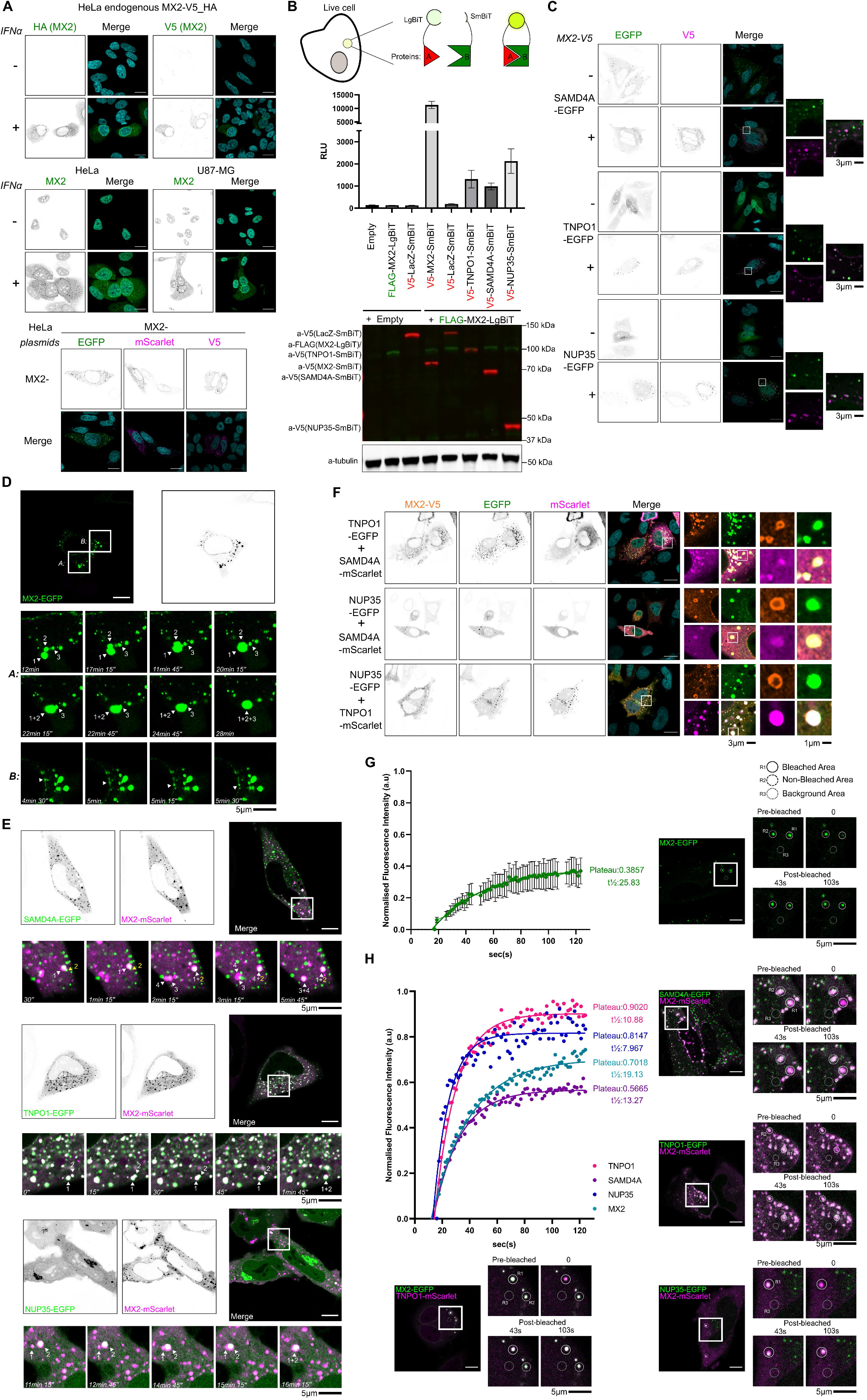
SAMD4A, TNPO1, and NUP35 interact with MX2 at dynamic cytoplasmic biomolecular condensates. **(A)** Confocal microscopy of: (Top) HeLa cells with endogenously V5- and HA-tagged MX2 alleles, (middle) HeLa and U87MG cells left untreated or treated with 1,000 IU ml^−1^ of rhIFN-a2a for 24 h. (Bottom): HeLa cells expressing EGFP-, mScarlet- or V5-tagged MX2. **(B)** HeLa cells mock- or (co)transfected with LgBiT/FLAG-tagged MX2 and SmBiT/V5-tagged MX2, LacZ, SAMD4A, TNPO1, or NUP35. Top: Schematic model of NanoBit assay. Reconstituted luminescence measurements for the (co)expressed LgBiT-, SmBiT-tagged proteins (mean ± s.d., 2 biological replicates). Bottom: Proteins expression were analysed by immunoblot using anti-FLAG, anti-V5, and anti-tubulin antibodies. Representative experiment performed at least three times. RLU: Relative light units. **(C)** Confocal microscopy of HeLa cells expressing EGFP-tagged SAMD4A, TNPO1, or NUP35 in the presence or absence of MX2-V5. **(D-E)** HeLa cells expressing **(D)** MX2-EGFP or **(E)** MX2-mScarlet together with EGFP-tagged SAMD4A, TNPO1, or NUP35 imaged under time-lapse conditions. Bottom “zoom” panels show an enlarged view of the white boxed areas. Moving and fusing MX2 condensates are indicated with white arrows and Smaug1 condensates with yellow arrows. Real imaging time is indicated at the bottom left side of each image. Scale bar: 10 µm. **(F)** Confocal microscopy of HeLa cells expressing MX2-V5 together with different combinations of EGFP- and mScarlet-tagged proteins. **(G-H)** HeLa cells expressing **(G)** MX2-EGFP or **(H)** MX2-EGFP together with TNPO1-mScarlet or EGFP-tagged SAMD4A, TNPO1 or NUP35 together with MX2-mScarlet. FRAP was performed for the complete individual condensates 24 h post-transfection. Left panel: Normalized FRAP curve(s) obtained for EGFP signal over time (n=10). Plateau and half-time are indicated on the right. Right panel: Representative images showing the biomolecular condensates during FRAP (before bleaching, during bleaching, and 43 and 103 sec post-bleaching). Bleached, non-bleached, and background area are highlighted with solid, dashed, or dotted white circles, respectively. Scale bar: 10 µm. **(A, C, F)** Nuclei were stained with DAPI (cyan). Scale bar: 20 µm. Right “zoom” panels of the white boxed area in individual fluorescence channels and channels overlay (without DAPI). For all fluorescence imaging, single channels are shown in inverted grey and the merged images with colours indicated in each panel.

Next, we performed live cell-imaging experiments using EGFP or mScarlet tagged MX2 that maintained anti-HIV-1 but had lost anti-HSV-1 activity (Figure S4A). In MX2-EGFP expressing HeLa cells, the MX2 puncta were highly dynamic and coalesced into larger condensates. These results are consistent with MX2 undergoing an intracellular phase transition and becoming enriched in cytoplasmic biomolecular condensates, some of which were also associated with the nuclear envelope (Figure 3D, Video S1, S1_ZoomA and S1_ZoomB). In live cells, SAMD4A colocalized with the MX2 condensates and also presented as SAMD4A-containing Smaug1 condensates adjacent to MX2 condensates. Similarly, TNPO1- and NUP35-positive MX2 condensates were observed that fused with each other (Figure 3E and Video S2, S2_Zoom, S3, S3_ Zoom, S4 and S4_ Zoom). We observed cytoplasmic colocalization of MX2 with SAMD4A, TNPO1, and NUP35, indicating that all four proteins are components of the same cytoplasmic condensates (Figure 3F). Biomolecular condensates can have distinct material states including liquid, hydrogel, liquid crystal, and amyloid-like fibrils that can be distinguished by their fluidity and (ir)reversible assembly^53^. Fluorescent recovery after photobleaching (FRAP) showed high exchange rate of protein constituents which marks biomolecular condensates. Variation in FRAP plateaus likely reflects the different physicochemical properties of the constituent proteins. For EGFP-tagged MX2, FRAP reached a plateau of 39% reconstitution within 100sec. Notably, in the presence of TNPO1-mScarlet, MX2-EGFP signal recovery was increased, suggesting a higher fluidity (Figure 3G and 3H). Collectively, this shows that MX2 forms dynamic cytoplasmic biomolecular condensates that comprise SAMD4A, TNPO1, and NUP35.

### MX2 assembles FG-Nups and importin-beta transporters at cytoplasmic biomolecular condensates

Our proteomics data suggest that MX2 interacts with FG-Nups and importin-beta transporters. FG-Nups create a selective nuclear pore permeability barrier and can spontaneously self-assemble into dense hydrogels with NPC-like permeability^42,54^. Importin-beta transporters, on the other hand, have multiple FG-binding pockets that facilitate the rapid transfer of their cargo into the nucleus^55,56^. We hypothesized that MX2 could promote the assembly of FG-Nups to create an NPC-mimicking compartment in the cytoplasm. To test this, we selected NUPL1 as FG-Nups and IPO4 as another importin-beta representative, respectively. NUPL1 overexpression induced cytoplasmic condensates which became MX2-positive in the presence of MX2 (Figure 4A). The diffused cytoplasmic distribution of IPO4, however, completely changed in the presence of MX2 with IPO4 relocating to the MX2 condensates (Figure 4A). A closer inspection revealed that both NUPL1 and MX2 assembled at the outer face of the same condensates even during the fusion processes as observed in the presence of IPO4 (Figure 4B-D). In live cells, EGFP-tagged NUPL1 was localized at the surface of fusion-competent mScarlet-tagged MX2 condensates, while IPO4-positive MX2 condensates were also observed fusing in the cytoplasm and interacting with MX2 at the nuclear membrane (Figure 4E and Video S5, S5_Zoom, S6 and S6_Zoom). Furthermore, the rapid signal recovery after photobleaching of IPO4 confirmed that importin-beta transporters interact with low affinity with FG-Nups and thus can easily cross the rim of the MX2 condensates in a similar way as they traverse the NPC barrier. In contrast, NUPL1 and MX2 had a lower fluidity than IPO4 suggesting that they were part of a distinct subcompartment with more gel-like phase separation material state at the cytoplasmic boundary of these condensates (Figure 4F). We confirmed that endogenous and ectopically expressed MX2 condensates could assemble endogenous FG-Nups (detected with the Mab414: mainly recognizes NUP358/RANBP2, NUP214 and NUP62)^57^ and NUP35 (Figure 4G and 4H). These results highlight MX2’s ability to assemble FG-Nups and importins-beta to form nuclear pore-mimicking biomolecular condensates.

**Figure 4:**
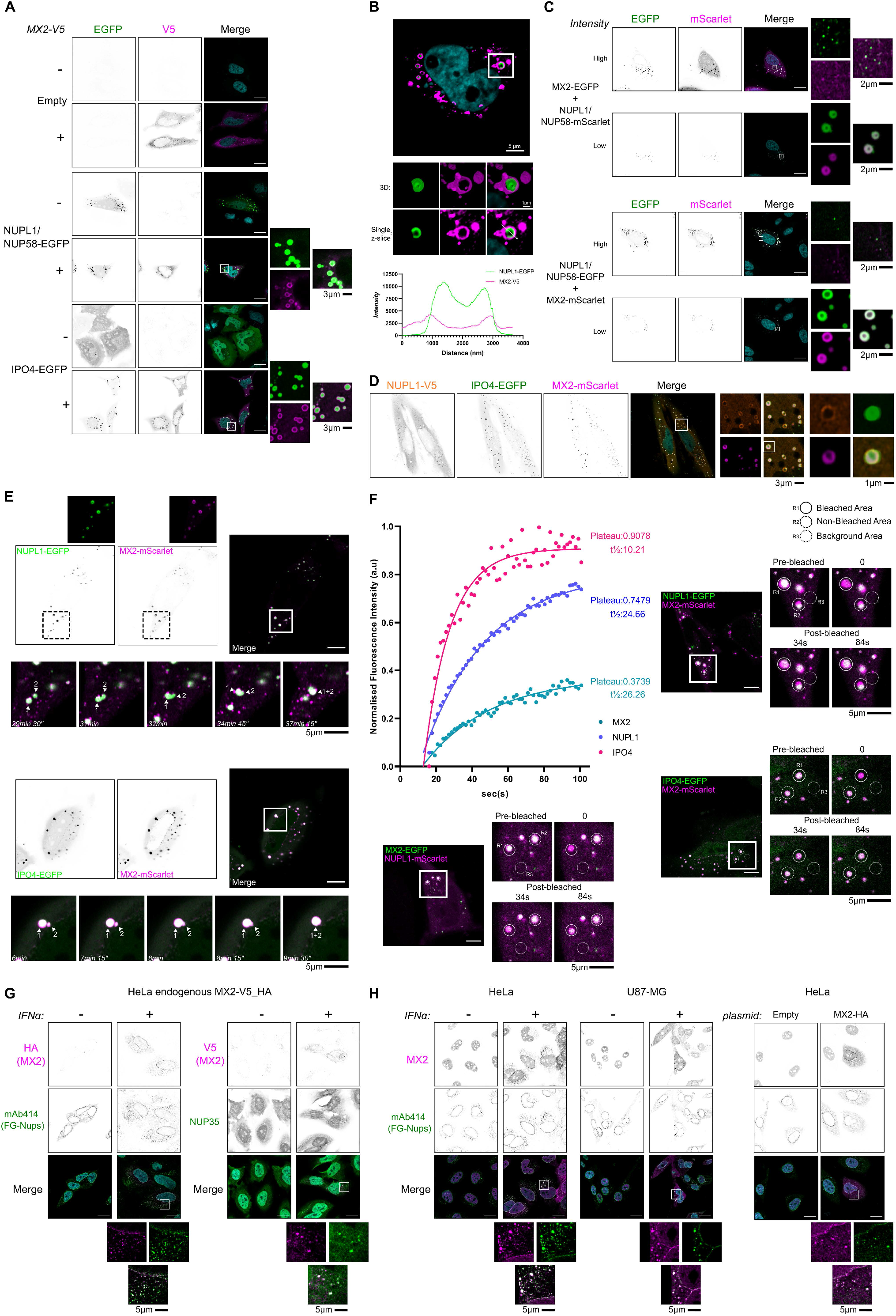
MX2 assembles FG-Nups and importins-beta at cytoplasmic biomolecular condensates. **(A)** Confocal microscopy of HeLa cells mock-transfected or expressing EGFP-tagged NUPL1 or IPO4 in the presence or absence of MX2-V5. **(B-F)** HeLa cells expressing **(B)** NUPL1-EGFP and MX2-V5 from panel **A**. Confocal microscopy with z-series image acquisition. Top: Image representing a single z-slice of the complete cell. Middle: “zoom” panels show an enlarged view of the white boxed area in individual fluorescence channels and channels overlay, represented as 3D volume reconstruction or as a single z-slice. Bottom: Intensity profile graph of NUPL1 and MX2. **(C)** NUPL1-EGFP or mScarlet and MX2-EGFP or mScarlet analysed by confocal microscopy. High and low intensity (display) setting are shown for each confocal image. **(D)** NUPL1-V5, MX2-mScarlet and IPO4-EGFP analysed by confocal microscopy. **(E)** mScarlet-tagged MX2 together with EGFP-tagged NUPL1 or IPO4 were imaged under time-lapse conditions. For MX2 and NUPL1 individual fluorescence channels, upper ‘’zoom’’ panels show an enlarged view of the highlighted area (white dashed squares). Bottom “zoom” panels show an enlarged view of the highlighted white boxed areas. Moving and fusing MX2 condensates are indicated with white arrows. Real time of imaging is indicated at the bottom left side of each image. Scale bar: 10 µm. **(F)** MX2-EGFP together with NUPL1-mScarlet or expressing EGFP-tagged NUPL1 or IPO4, together with MX2-mScarlet. FRAP was performed for the complete individual condensates 24 h post-transfection. Left panel: Normalized FRAP curves obtained for GFP signal over time (n=7). Plateau and half-time are indicated on the right of each curve. Right panel: Representative images showing the biomolecular condensates during FRAP (before bleaching, during bleaching, and 34 and 84 sec post-bleaching). Bleached, non-bleached, and background area are highlighted with solid, dashed, or dotted, white circles, respectively. Scale bar: 10 µm. **(G)** Confocal microscopy of HeLa cells with endogenously V5- and HA-tagged MX2 alleles, left untreated or were stimulated with 1,000 IU ml^−1^ rhIFN-a2a. Intracellular localization of MX2, endogenous FG-Nups and NUP35 was assessed 24 h post-IFN-stimulation by confocal microscopy. **(H)** Confocal microscopy of: Left: HeLa and U87MG cells left untreated or induced with 1,000 IU ml^−1^ rhIFN-a2a. Right: HeLa mock-transfected or expressing HA-tagged MX2. Intracellular localization of MX2 and endogenous FG-Nups was assessed 24 h post-IFN-stimulation/transfection by confocal microscopy. **(A, B, C, D, G, H)** Nuclei were stained with DAPI (cyan). Scale bar: 20 µm. Right or bottom “zoom” panels show an enlarged view of the white boxed area in individual fluorescence channels and channels overlay (without DAPI). For all fluorescence imaging, single channels are shown in inverted grey and the merged images with colours indicated at each panel.

### MX2 cytoplasmic biomolecular condensates associate with cytoplasmic MLOs and contain RNA-binding and disordered domain proteins

The second cluster of proteins in our proteomics data showed an association of MX2 with components of ribonucleoprotein granules that have LLPS properties such as PBs and SGs. We selected LSM14A (identified in the MX2 BioID) and G3BP1 (not identified in the MX2 BioID) as respective marker proteins of these MLOs^45^. In both fixed and live cells, LSM14A was detected within MX2 condensates while mainly present in separate LSM14A-containing PBs that were in close proximity to and interacting with MX2 condensates (Figure S5A, S5B and Video S7 and S7_Zoom). In contrast, the diffuse cytoplasmic pattern of G3BP1 was not altered in the presence of MX2. However, after arsenite-induced stress, G3BP1 localized, as expected, into SGs and some of the MX2 biomolecular condensates were observed close to or overlapping with these SGs (Figure S5C).

Next, we investigated SAMD4B, FUBP3, and CNOT4 as representative proteins with disordered and/or RNA-binding domains, which co-localized with MX2 and NUPL1 in dynamic cytoplasmic condensates (Figure S5D and S5E). These condensates exhibited rapid movement and fusion in the cytoplasm (Figure S5F and Video S8, S8_Zoom, S9 and S9_Zoom). SAMD4B and FUBP3 displayed rapid FRAP reconstitution, indicating their presence in a highly fluid subcompartment of MX2 condensates with liquid-like properties (Figure S5G). These findings suggest that MX2 cytoplasmic condensates are multiprotein assemblies capable of associating with cytoplasmic MLOs.

### The N-terminal disordered domain and dimerization of MX2 are required for cytoplasmic condensate formation

MX2 could act as a driver or client of condensate formation. If MX2 acts as a driver, certain regions within its sequence must facilitate the required molecular interactions. In silico analysis (dSCOPE^58^) of MX2’s primary structure predicted the N-terminal regions 23-54 and 64-87 as potential sites for phase separation (Figure 5A).

**Figure 5:**
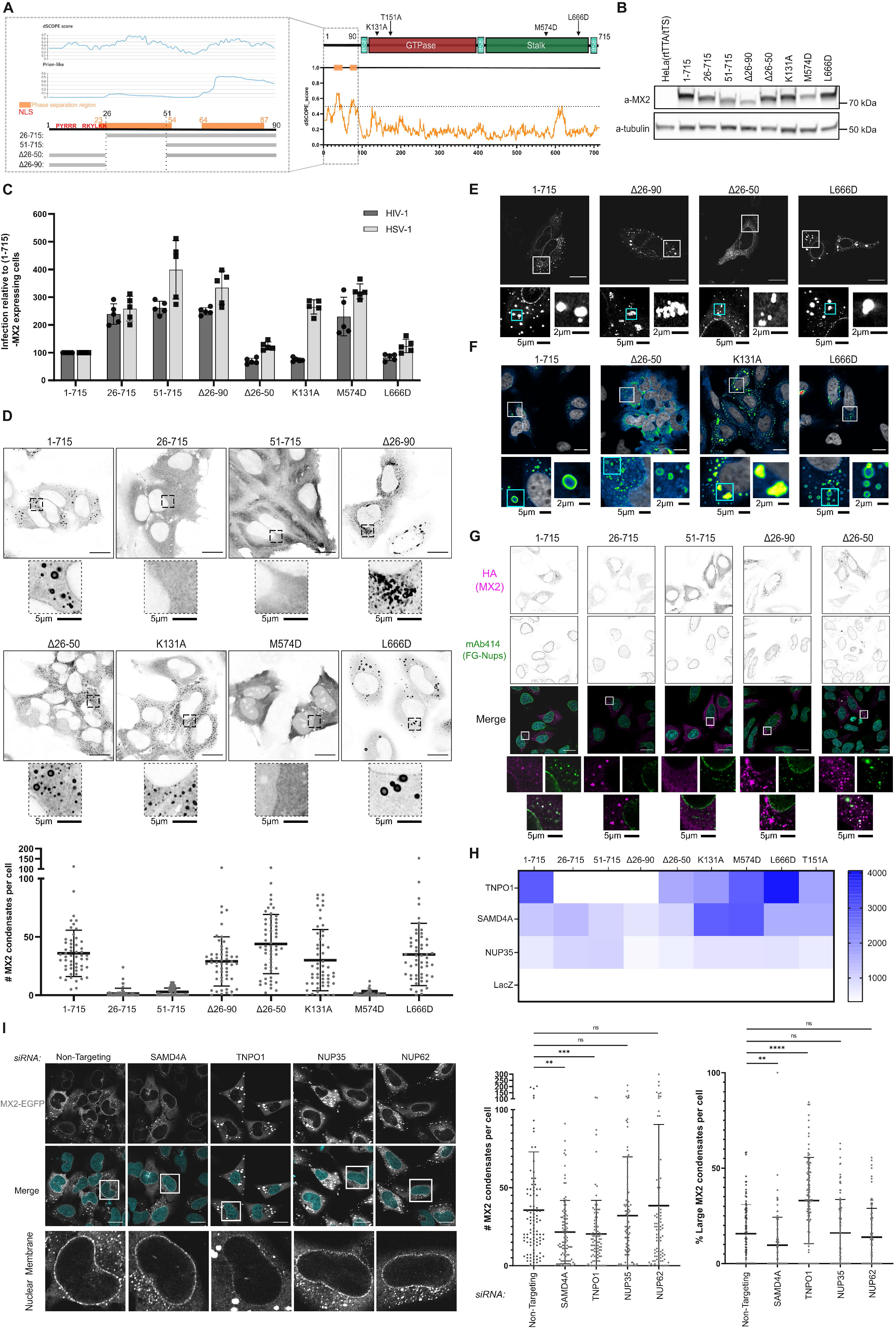
MX2 condensate formation requires the N-terminal domain and dimerization and is regulated by TNPO1 and SAMD4A. **(A)** Right: Domain structure of MX2 with single amino acid substitutions indicated. Disorder plot dSCOPE scores (y-axis) of MX2 (X-axis). Left: Disorder plots dSCOPE scores and Prion-like score (y-axis) of MX2 90 N-terminal amino acids (X-axis). Phase separation region, (NLS) according to ^4^, and N-terminal MX2 deletion mutations are indicated. **(B-D, F)** HeLa (rtTA/tTS) cells and doxycycline-inducible HeLa cells expressing MX2(1-715) or mutant variants were treated with different doxycycline amounts for comparable expression levels. **(B)** Twenty-four hours after stimulation, protein expression was analysed by immunoblot using anti-MX2 and anti-tubulin antibodies. **(C)** Antiviral activities of MX2 mutant proteins against HSV-1 strain C12 and VSV-G.HIV-1-GFP, relative to HeLa MX2(1-715) samples which are each set to 100 (mean ± s.d., n = 5 independent experiments). **(D)** Localization of V5-tagged MX2 proteins and number of MX2 condensates (>0.1 µm^3^ volume) per cell. **(E)** HeLa cells expressing C-terminally EGFP-tagged MX2 proteins. Localization of EGFP-tagged MX2 proteins visualized in white. **(F)** Localization of V5-tagged MX2 proteins visualized using pseudocoloring blue to yellow gradient based on intensity. **(G)** HeLa cells expressing C-terminally HA-tagged MX2 proteins. **(H)** HeLa cells (co)expressing LgBiT/FLAG-tagged MX2 mutants and SmBiT/V5-tagged LacZ, TNPO1, SAMD4A, and NUP35. Heat map representing reconstituted luminescence measurements for all the combinations of LgBiT-, SmBiT-tagged proteins (mean ± s.d., 2 biological replicates). Protein expression was analysed by immunoblot using anti-FLAG, anti-V5, and anti-tubulin antibodies (see **Figure S6F**). Representative experiment performed at least three times. **(I)** Doxycycline-inducible HeLa MX2(1-715) C-terminally EGFP-tagged were transfected with siRNAs and stimulated with 100 ng mL^-1^ doxycycline. Number of MX2 condensates (>0.1 µm^3^ volume) and % of large MX2 condensates (>2 µm^3^ volume) per cell (mean ± s.d., two-tailed unpaired *t*-test to Non-Targeting siRNA; *p* values indicated with \**p* < 0.05, \*\**p* < 0.01, \*\*\**p* < 0.001). Localization of EGFP-tagged MX2 proteins visualized in white. **(D, E, F, G, I)** Intracellular localization of: (**D, E, F, I)** MX2, **(G)** MX2 and endogenous FG-Nups was assessed 24 h (for **D, E, F, G)** or 48 h (for **I)** post-transfection/doxycycline-stimulation by confocal microscopy. Nuclei were stained with DAPI **(I, G)** (cyan), **(F)** (white). Scale bar: 20 µm. Bottom “zoom” panels show an enlarged view of the highlighted area (without DAPI). For **(D, I)** Confocal microscopy with z-series image acquisition. For each comparison, either **(D)** 54 cells or **(I)** 96 cells were randomly selected. For all fluorescence imaging, single channels are shown in inverted grey and the merged images with colours indicated at each panel, unless indicated otherwise.

We created 8 doxycycline-inducible HeLa cell lines expressing different V5-tagged MX2 mutants to examine the influence of MX2’s N-terminal region, GTP-binding and -hydrolysis, and oligomerization on its localization and antiviral activity (Figure 5A, 5B, S6A and S6B). MX2(26-715)^8,9,21,25^ and MX2(51-715) mutants, lacking antiviral activity, displayed diffuse cytoplasmic expression with few large condensates (Figure 5C and 5D). MX2(Δ26-90), lacking both predicted phase separation regions, failed to restrict the viruses and exhibited reduced nuclear membrane localization, forming dynamic condensate-like structures which did not coalesce into larger condensates (Figure 5C-E and Video S10). In contrast, MX2(Δ26-50), retaining one phase separation region, maintained its antiviral activity and formed condensates (Figure 5A, 5C-E and Video S14). GTP-binding-deficient MX2(K131A) inhibited HIV-1 but not HSV-1^5,6,9,21,59^, formed condensates with altered morphology with a more homogenous MX2 distribution and a less intense outer ring-like pattern (Figure 5C, 5D and 5F). Monomeric MX2(M574D) did not inhibit either virus^9,20,22,23^ and had a completely altered localization with a diffuse nuclear and cytoplasmic pattern maintaining some localization in the nuclear membrane. Dimeric interface 1 mutant MX2(L666D) retained properties similar to MX2(1-715)^20,22,23,60^ (Figure 5C-F). Transient expression of the same EGFP-tagged MX2 mutants displayed similar localization patterns. This included MX2(T151A), which can bind but not hydrolyze GTP and had strong nuclear membrane accumulation maintaining the ability to form cytoplasmic condensate (Figure S6C and Videos S10-S18). Overall, MX2 biomolecular condensation and both HIV-1 and HSV-1 restriction depended on the N-terminal disordered region and MX2 dimerization.

### TNPO1 and SAMD4A regulate phase separation properties of MX2

MX2’s interaction with nucleoporins is crucial for its anti-HIV-1 activity^37^. While MX2(1-715) and MX2(Δ26-50) associated with nuclear membranes, and their condensates assembled FG-Nups (mAb414), MX2(26-715), MX2(51-715), or Mx2(Δ26-90) did not (Figure 5G). MX2 N-terminal domain chimeric MX1 constructs displayed similar properties as the corresponding MX2 truncated mutants (Fig S6D), while chimeric EGFP constructs failed to form condensates or associate with FG-Nups in the cytoplasm or nuclear envelop, likely due to EGFP’s monomeric nature (Fig S6E).

We next characterized MX2 mutants’ ability to bind TNPO1, SAMD4A, and NUP35. MX2(26-715), MX2(51-715) and MX2(Δ26-90), failed to bind TNPO1. SAMD4A exhibited higher association for MX2(K131A) and MX2(M574D) (Figure 5H and S6F). In doxycycline-inducible HeLa MX2(1-715)-EGFP cells, TNPO1 KD reduced MX2 nuclear membrane accumulation and induced fewer, but larger condensates. SAMD4A depletion decreased cytoplasmic condensates without affecting MX2 nuclear membrane accumulation. NUP35 or NUP62 KD had no effect on MX2 localization or condensate formation (Figure 5I). Thus, both TNPO1 and SAMD4A affected MX2 biomolecular condensate formation, which for TNPO1, was dependent on MX2 N-terminal motifs.

### MX2 condensates mimic the nuclear pore LLPS environment to interact with incoming HIV-1 and HSV-1 capsids

MX2 restricts HIV-1 and HSV-1 by preventing nuclear entry of their genome^11^. To gain insights about the potential impact of the MX2 condensates on this mechanism, we investigated the fate of incoming HIV-1 and HSV-1 capsids in the presence or absence of MX2(1-715) and mutant MX2. For HIV-1, doxycycline-inducible EGFP-tagged MX2(1-715), and -(T151A) but not MX2(26-715) HeLa cells generated cytoplasmic and nuclear envelope-associated MX2 condensates that contained incoming HIV-1 capsids (Figure S7A-D Figure 6A). Similar observations were made with endogenous MX2 and wt envelope HIV-1 infection (Figure S7E). Additionally, we infected HeLaP4 cells expressing MX2-mScarlet with HIV IN-EGFP and followed single functionally active pre-integration complexes (PICs) during infection by visualizing the fluorescently labelled HIV-1 integrase (IN). Nuclear entry of HIV-1 PICs was prevented, and we observed EGFP signal colocalizing with cytoplasmic MX2 condensates (Figure 6B).

**Figure 6:**
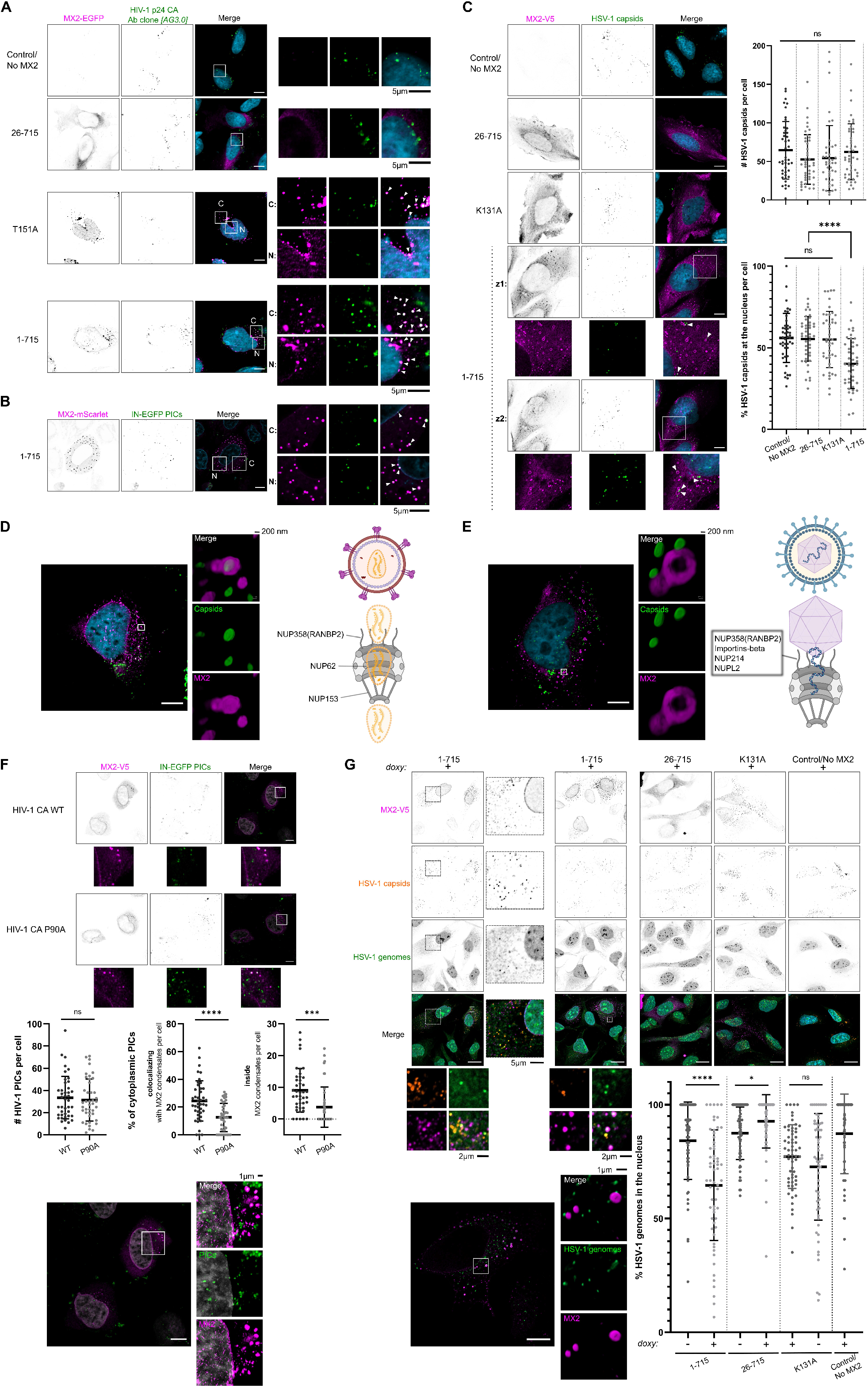
MX2 condensates mimic the NPC LLPS environment to interact with incoming HIV-1 and HSV-1 capsids. **(A)** Confocal microscopy of VSV-G.HIV-1-GFP virus capsids intracellular localization in HeLa (rtTA/tTS-Control/NoMX2) and doxycycline-inducible HeLa cells expressing similar amounts of EGFP-tagged MX2(1-715) or mutant MX2 variants. C: cytoplasm, N: nuclear envelope. **(B)** Confocal microscopy of HIV IN-EGFP particles intracellular localization in_HeLaP4 cells expressing MX2(1-715)-mScarlet. C: cytoplasm, N: nuclear envelope. **(C)** Confocal microscopy analysis of HIV-1 capsids intracellular localization in HeLa (rtTA/tTS-Control/NoMX2) and doxycycline-inducible HeLa cells expressing similar amounts of V5-tagged MX2(1-715) or mutant MX2 variants. z1, z2: z-slice 1, 2. Right: total intracellular HSV-1 capsid number and percentage of HSV-1 capsids coinciding with the nuclear staining per cell. For each comparison, 44 cells were randomly selected (mean ± s.d., two-tailed unpaired *t*-test to cells Control/NoMX2; *p* values indicated with \**p* < 0.05, \*\**p* < 0.01, \*\*\**p* < 0.001). **(D)** HeLa cells expressing MX2(1-715)-EGFP infected with HIV-1 based lentiviral vector from **Figure S7D**. Single z-slice view of the complete cell. Middle: enlarged view of the highlighted area in individual fluorescence channels and channel overlay, represented as 3D volume reconstruction. Right: graphs showing the HIV-1^49^ capsids NPC association. Figure created with BioRender software. **(E)** HeLa MX2(1-715)-V5 cells infected with HSV-1 from **(C).** Single z-slice view of the complete cell. Middle panel: enlarged view of the highlighted area in individual fluorescence channels and channel overlay, represented as 3D volume reconstruction. Right: graphs showing the HSV-1^48^ capsids NPC association. Figure created with BioRender software. **(F)** Confocal microscopy of HIV IN-EGFP (wild-type capsid and P90A capsid) intracellular localisation in doxycycline-inducible V5-tagged MX2(1-715) HeLa cells. Middle: total intracellular HIV-1 particles number and percentage of cytoplasmic HIV-1 particles colocalizing or coinciding completely with MX2 condensates per cell. For each comparison, 42 cells were randomly selected (mean ± s.d., two-tailed unpaired *t*-test to cells between CA WT and P90A infected cells; *p* values indicated with \**p* < 0.05, \*\**p* < 0.01, \*\*\**p* < 0.001). Bottom: HeLa MX2(1-715)-V5 cells infected with HIV IN-EGFP WT CA. Single z-slice view of the complete cell. Enlarged view of the highlighted area in individual fluorescence channels and channel overlay, represented as 3D volume reconstruction. **(G)** CLICK experiments for the analysis of intracellular EdC/A labelled HSV-1 genomes in HeLa (rtTA/tTS-Control/NoMX2) and doxycycline-inducible HeLa cells expressing similar amounts of V5-tagged MX2(1-715) or mutant MX2 variants. Top: Intracellular localization of the MX2 proteins, incoming viral capsids and viral genomes was assessed by confocal microscopy. Bottom left: High-resolution image of HeLa MX2(1-715)-V5 cells infected with EdC/A pre-labelled HSV-1. Single z-slice view of the complete cell with enlarged view of the highlighted area in individual fluorescence channels and channel overlay, represented as 3D volume reconstruction. Right bottom: percentage of HSV-1 genomes coinciding with the nuclear DAPI area per cell. For each condition, 61 randomly selected cells were recorded/documented (mean ± SD, two-tailed unpaired t-test cells between not induced (-) or induced (+) with doxycycline for MX2 expression; p values indicated with *p < 0.05, **p < 0.01, ***p < 0.001). (**B-F** and **G bottom left)** Confocal microscopy with z-series image acquisition. Single z-slice view of the complete cell is shown. Nuclei were stained with (**A, C, D, E, G)** DAPI (cyan) or **(B, F)** NucSpot 650/665 **(B)** (cyan) **(F)** white. Scale bar: (**A-G)** 10 µm; Bottom or right “zoom” panels show an enlarged view of the highlighted area (without DAPI). For all fluorescence imaging, single channels are shown in inverted grey and the merged images with colours indicated at each panel, unless indicated otherwise.

In doxycycline-inducible V5-tagged MX2(1-715), (26-715), and (K131A) HeLa cells, only MX2(1-715) colocalized with incoming HSV-1 capsids and prevented their targeting to the nuclei. In contrast to the HIV-1 capsids, HSV-1 capsids colocalized with MX2(1-715) condensates at the boundary between the condensates and the cytoplasm, and thus seemed to be unable to penetrate the surface of these condensates. HSV-1 capsids that did not colocalize with MX2 were also observed in the cytoplasm of the HeLa MX2(1-715) cells (Figure 6C).

The distinct association of HIV-1 or HSV-1 capsids with MX2 condensates resembles their distinct mode of nuclear targeting. HIV-1 capsids can dock at the NPC cytoplasmic filaments (NUP358/RANBP2) and then, by interacting with Nups of the central channel (NUP62) and nuclear basket (NUP153), can reach the nucleoplasm^47,49,61,62^ (Figure 6D). In contrast, HSV-1 capsids dock via importins-beta to the NPCs, likely by interactions with NUP358(RANBP2), NUP214 and NUPL2, to release their genome through the NPCs, and leave empty, cytosolic capsids behind^48,63^ (Figure 6E).

To bolster our understanding of MX2’s role in restricting HIV-1 PICs from nuclear entry, we compared its ability to associate with incoming HIV IN-EGFP (PICs) between wild-type CA and the P90A (CypA binding deficient) MX2 escape mutant^5,64^. We observed a reduced presence of P90A CA IN-EGFP PICs colocalizing and within MX2 condensates compared to wild-type CA (Figure 6F).

To further investigate the effect of MX2 on the intracellular fate of incoming genomes, we inoculated HeLa cells with HSV-1 virions harboring genomes to which we could cross-link fluorescent dyes. The nuclei of HeLa cells expressing V5-tagged MX2(1-715) contained fewer viral genomes than the same cells without doxycycline induction or the HeLa(rtTA/tTS-Control/NoMX2) cells. Instead, the incoming HSV-1 genomes often co-localized with the cytoplasmic MX2 condensates (Figure 6G). Interestingly, like the cytoplasmic HSV-1 genomes, the capsid signals in the MX2(1-715) expressing cells appeared more spread than in the respective control cells (Figure 6C and 6G). In cells expressing the GTPase deficient mutant MX2(K131A), there was also a non-significant reduction, and in cells with the short isoform MX2(26-715) a slight increase in nuclear HSV-1 genomes. As observed for the HSV-1 capsids, the genomes were located on the surface of MX2(1-715) condensates (Figure 6G and S7F). Collectively, these results show that the MX2 biomolecular condensates associate with incoming HIV-1 and HSV-1 capsids and genomes, and thereby prevent their nuclear targeting.

### MX2 condensates interact with incoming HIV-1 and HSV-1 capsids at different steps of their journey towards the nucleus

Both HIV-1 and HSV-1 capsids use microtubules (MTs) for transport towards the nucleus^48,65^. MX2 condensates appeared to colocalize with MTs, suggesting their capability to move along these structures (Figure 7A, S8A). To further explore the possible role of MTs in facilitating the encounter of MX2 condensates with viral capsids travelling along the MTs, cells were treated with nocodazole, a MT depolymerizing agent. This treatment did not affect MX2 nuclear membrane localization, condensate formation, or distribution (Figure S8B). The levels of HIV-1 infection remained unaffected in nocodazole treated cells^66^, however, the MX2-mediated HIV-1 restriction decreased with increasing disruption of the MT network (Figure 7B and S8C). In contrast, depolymerization of MTs significantly reduced HSV-1 infectivity (Figure 7C)^67,68^. Surprisingly, MX2’s ability to hinder HSV-1 infection increased with increasing nocodazole concentrations (Figure 7C).

**Figure 7:**
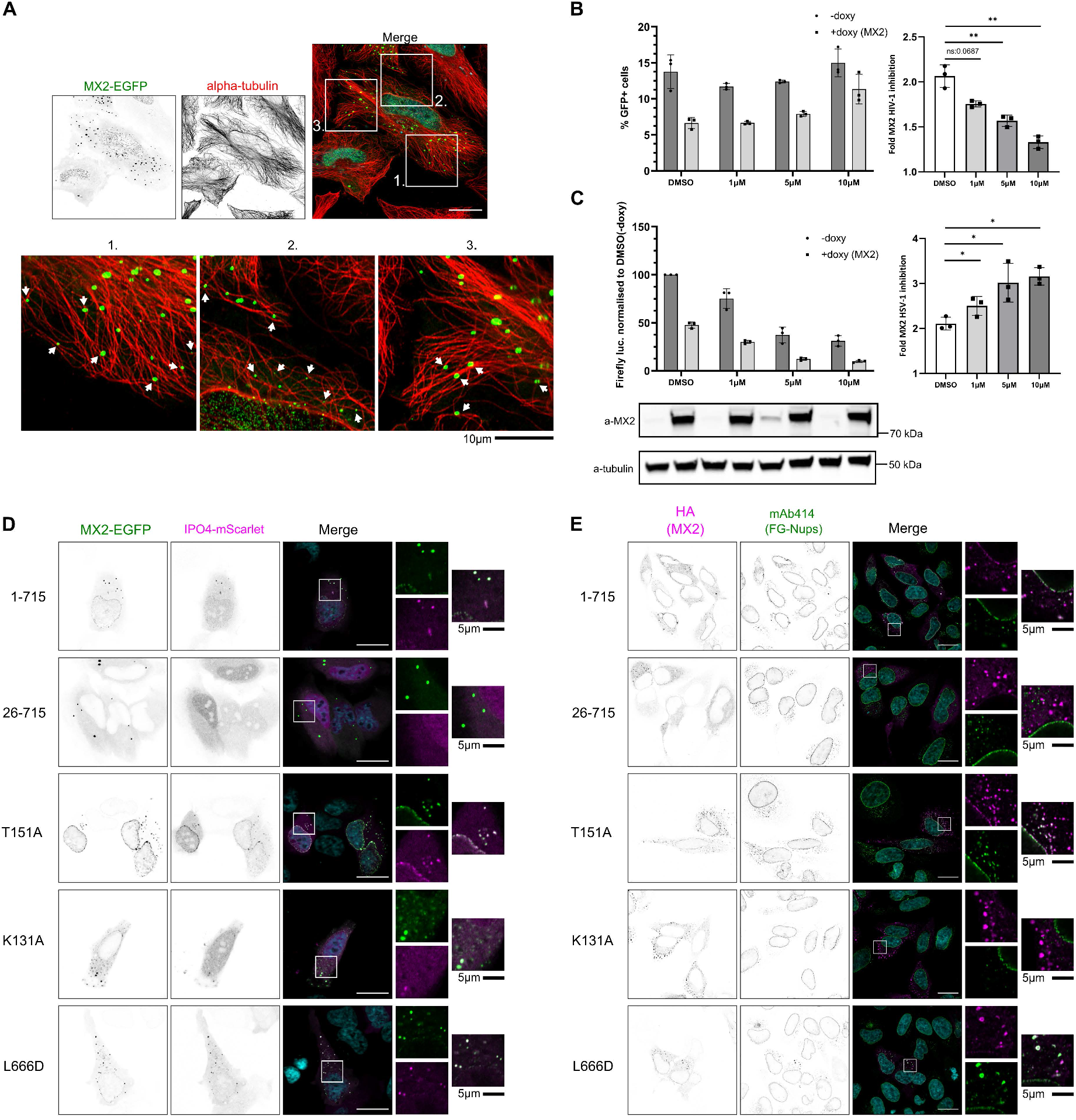
MX2 condensates interact with incoming HIV-1 and HSV-1 capsids at different steps of their journey towards the nucleus. **(A)** HeLa cells expressing MX2-EGFP. Intracellular localization of MX2-EGFP and endogenous alpha-tubulin were assessed 24 h post-transfection by confocal microscopy. MX2 condensates colocalizing with MTs are indicated with white arrows. **(B, C)** Antiviral experiments preformed in HeLa MX2 cells with **(B)** VSV-G.HIV-1-GFP and **(B)** HSV-1 firefly reporter virus in the presence of DMSO or 1, 5, or 10 µM nocodazole. Data analysis similar to **Figures S3B and S3C,** respectively. **(D)** HeLa cells expressing IPO4-mScarlet and EGFP-tagged MX2(1-715) or mutant MX2 variants. **(E)** HeLa cells expressing HA-tagged MX2(1-715) or mutant MX2 variants. **(D, E)** Intracellular localization of **(D)** EGFP- and mScarlet tagged proteins **(E)** MX2 and endogenous FG-Nups were assessed 24 h post-transfection by confocal microscopy. Nuclei were stained with (**A, D, E)** DAPI (cyan). Scale bar: 20 µm. Bottom or right “zoom” panels show an enlarged view of the highlighted area (without DAPI).

Importantly, all condensates with MX2 mutants that retain the first 25 residues, accumulated importins-beta (IPO4), which facilitate viral translocation from the centrosome to the NPCs and which dock to FG-rich cytoplasmic filaments. While the GTPase deficient MX2(K131A) generated an NPC-mimicking environment, it assembled distinct FG-Nups than the other MX2 mutants (Figure 7D and 7E).

In conclusion, MX2 condensates interact with HIV-1 capsids during their travel along the MT network and with HSV-1 capsids at a different step of their journey towards the nucleus. Additionally, the delocalization of importins-beta is independent of GTP-binding and - hydrolysis but is controlled by the first 25 (NTD) residues of MX2, while the specific assembly of FG-Nups is influenced by MX2 GTPase activity.

## Discussion

MX2 can restrict retroviruses and herpesviruses, two highly divergent virus families. Despite a growing body of literature, the underlying mechanisms of MX2’s antiviral activity remain largely elusive, although evidence for the involvement of additional host proteins is increasing^11^.

MX2’s proximity interactome reported here, was strongly enriched with two protein clusters: FG-domain related nucleocytoplasmic transport components and cytoplasmic granule-associated ribonucleoproteins. Remarkably, both protein clusters have a high propensity to organize and function as LLPS compartments.

FG-rich Nups from different nucleopore subcomplexes contribute to HIV-1 infection, MX2 subcellular localization, and MX2-mediated HIV-1 restriction ^37,61^. Nups such as NUP35, NUP62, NUP153, NUP88, NUP98, and NUP358(RANBP2) can bind HIV-1 capsids and participate in the MX2-mediated HIV-1 restriction ^37,47,49,61,69^. We identified TNPO1 (previously reported for HIV-1^37,61^), SAMD4A, and NUP35 as significant contributors to MX2 activity against both HIV-1 and HSV-1, while NUP62 influenced only the restriction of HIV-1.

SAMD4A, TNPO1, and NUP35 associated with the punctate cytoplasmic fraction of MX2, in poorly characterized cytoplasmic structures^20,21,51,61^, often referred to as puncta^8,17,25,26^. Supported by proteomics, co-localization, live cell imaging, and FRAP analysis, we firmly establish that MX2 forms dynamic cytoplasmic biomolecular condensates. MX2 condensates are multiprotein and heterogenous, comprising FG-Nups. Moreover, they can recruit RNA-binding and/or disordered domain-containing proteins and can associate with PBs and SGs. FG-Nups can undergo LLPS creating three dimensional networks with hydrogel-like properties that mimic the permeability features of the NPCs^42,43,70,71^. MX2 assembled FG-Nups at the outer face of the condensates, creating a gel-like subcompartment, that allowed dynamic interactions with importins-beta.

MX2 condensate formation depended on its dimerization and its 90 residue NTD. Different MX2 regions in the NTD contribute to the formation of FG-Nup-rich condensates, with the 1-25 amino acid residues and the second intrinsically disordered region facilitating the assembly of FG-Nups/importins-beta^37,38^ and phase separation properties, respectively. GTP-binding and -hydrolysis, however, are dispensable for condensate formation but affect their properties. The reduced presence of MX2 GTP-binding defective mutant (K131A) at the surface of the condensates can be a consequence of the dependency on GTP-binding for MX2 interaction with NUP358(RANBP2) which was described to also result in a reduced accumulation of MX2(K131A) at the nuclear membrane^59^. Previous reports where the first 90 amino acid residues of MX2 were transferred to MX1 or irrelevant dimerization competent proteins, conferred the ability to restrict HIV-1^4,26,61^ and HSV-1^9^ while some studies also reported cytoplasmic MX2 NTD-driven puncta formation^26,72,73^.

MX2 antiviral condensate formation is regulated by a fine-tuned balance between MX2’s structural characteristics and interacting proteins. Cellular factors such as TNPO1 and SAMD4A, which regulate condensate formation, impact MX2’s HIV-1 and HSV-1 restriction, while cellular factors with varying levels of involvement between the two activities (e.g. NUP62), only alter the composition of the condensates, possibly influencing the MX2 mechanism by facilitating viral antigen-MX2 binding. We confirmed that TNPO1-MX2 binding is controlled by the first 25 residues, in line with Dick et al.,^37^ but additionally requires at least the second part of the 26-90 NTD, which promotes phase separation. TNPO1 is a regulator of phase separation properties by acting as a chaperone for its interacting proteins. It has been proposed that the affinity of the NLS for TNPO1 may be modulated by post-translational modifications (PTMs) in or near the NLS^74–77^. Phosphorylation, can either enhance or reduce phase separation and recently emerged as a bidirectional modulator of MX2 anti-HIV-1 activity that controls MX2 binding to TNPO1 and nuclear membrane accumulation^38,78–80^.

Together, these data raise an intriguing question: Are there two independent pools of MX2, one at the nuclear rim and one in the condensates, or are these pools connected and interchangeable? Our time-lapse microscopy showed association of MX2 cytoplasmic condensates with the more intense nuclear membrane localized MX2 punctate fraction (see Videos S1, S1_ZoomB, S3, S11, S17). Additionally, MX2 localization at the outer face of the NPCs depends on its condensate forming properties as shown by the reduced or absent nuclear membrane accumulation of the fusion incompetent MX2(Δ26-90), MX2(1-715) in TNPO1 KD cells^37^, chimeric _(Δ26-90_ΜΧ2)_MX1, and chimeric _(1-90_MX2)_EGFP proteins. These observations suggest that the cytoplasmic and nuclear membrane fractions of MX2 share similar LLPS properties. Examining the varying nuclear membrane accumulation of different MX2 mutants can enhance our understanding of the properties of the NPC-bound fraction of MX2 in comparison to MX2 condensates.

MX2 binds to the HIV-1 core preventing the virus uncoating process, facilitated by the NTD and GTPase domain but not requiring GTP-binding and -hydrolysis^11,20,81,82^. More recently, it has been demonstrated that MX2-containing cell fractions can disassemble alphaherpesvirus capsids^13^. This process requires MX2 binding to HSV-1 capsids, GTPase activity and involves, to some extent, the first 25 residues of MX2^9,13^. Our antiviral and confocal imaging data demonstrate that only MX2 mutants that can form FG-Nups-rich condensates could restrict HIV-1 and HSV-1. The discovery of condensate formation as an essential feature of the MX2 antiviral mechanism, can explain why the MX2(26-715) short isoform, that maintains partial HSV-1 and HIV-1 capsid binding and disassembly activity in cell lysates but cannot form FG-Nups-rich condensates, fails to restrict either virus in a cellular context^8,9,13,81^. Moreover, the GTP-binding-defective MX2(K131A), which was shown to maintain binding to HIV-1 but not HSV-1 capsids, restricts only HIV-1 through its capability to generate cytoplasmic condensates, underlining the indispensable role of both capsid binding and phase separation^9,13,59^.

Incoming HIV-1 and HSV-1 capsids colocalized with MX2 condensates. High resolution imaging, however, revealed a striking difference: HIV-1 capsids resided inside MX2 condensates while HSV-1 capsids were positioned at the cytoplasmic boundary. These distinct MX2 condensate-viral capsid interaction patterns remarkably resemble how HIV-1 and HSV-1, in the absence of MX2, interact with the NPC upon nuclear targeting^62,63^ (Figure 6A-6E). Therefore, MX2-capsid binding is sufficient for the encapsulated viral genome of HIV-1 to enter the condensates, whereas the DNA genome of HSV-1 remains protected within the capsid outside of the condensates. This emphasizes the necessity for MX2 to disassemble HSV-1 capsids and release the viral genome to exert its anti-HSV-1 activity.

Based on (1) the MX2 condensate-viral capsids colocalization pattern, (2) the different virus-specific Nups requirements for MX2 antiviral activity, (3) the essential condensate-forming requirement of MX2’s anti-HIV-1 and -HSV-1 activity, and (4) the FG-Nups assembly at MX2 condensates, we propose that MX2 condensates mimic the NPC environment, therefore acting as nuclear pore decoys that facilitate MX2-capsid association.

Our nuclear pore-mimicking condensate model for MX2 can explain how Nups that localise at different sites of the NPC and can bind viral components, can affect MX2’s antiviral activity without altering its nuclear envelop accumulation^61^. Our data is not in line with the canonical view that MX2 primarily localizes at the cytoplasmic face of the NPC only interacting with NPC cytoplasmic filaments and outer-ring Nups, but supports a model of a previously overlooked LLPS-driven cellular compartment in which all these diversely localized Nups could associate with MX2 to facilitate its antiviral activity.

Recent studies showed that HIV-1 capsids mimic karyopherins to enter the FG phase of nuclear pores, strongly supporting our observations of HIV-1 capsids being able to penetrate MX2 condensates in a similar manner as the NPC^83,84^. Furthermore, HIV-1 capsid mutants partially (e.g. N74D CA, CPSF6 binding deficient) or completely insensitive (e.g. P90A CA, CypA binding deficient) to MX2 restriction, are still capable of interacting with MX2. These mutants have a lower nuclear entry efficiency with altered dependence on cofactors associated with nuclear entry, and are specifically insensitive to FG-rich NUP358(RANBP2) and NUP153 depletion allowing them to (partially) avoid the MX2 decoy condensates^5,6,20,47,49,61,85–88^. Reduced accumulation of P90A CA EGFP-IN PICs in MX2 condensates is in line with this (Figure 6F). MX2 escape mutations can modify the flexibility/conformation of CA either by preventing the binding to cofactors (e.g. CA CypA binding loop) or independent of cofactors (e.g. CA helix 6, 116A)^85,89,90^. Biophysical insights of HIV-1 CA nuclear entry demonstrated that the shape, orientation, and elasticity of the capsid are also important for nuclear entry^62,91–93^. Taken together, it is possible that the MX2 escape mutants can associate with MX2 condensates but are unable to (sufficiently) penetrate their surface due to altered or destabilized HIV-1 capsid properties.

HeLa cells expressing MX2(1-715) contained significantly fewer clickable HSV-1 genomes in the nucleus, but more in the cytoplasm compared to control cells. As the CLICK reagents have no access to genomes completely packaged within HSV-1 capsids, these data imply that MXB(1-715) had induced pre-mature capsid disassembly in the cytosol. This notion is also consistent with the localization of several clickable genomes on the surface of the MX2 biomolecular condensates, our observations of ‘’delayed’’ capsids in the cytoplasm, and with biochemical in vitro studies demonstrating MX2 induced disassembly of alphaherpesvirus capsids^13^. Such prematurely released HSV-1 genomes might expand and unroll as indicated by the different fluorescent signals in the cytoplasm when compared to nuclear clickable HSV1 genomes. At present, the fate of such cytosolic genomes remains unclear: degradation by cytosolic DNases, like Trex-1 or sensed by cGAS?

Disrupting the MT network reduced MX2 restriction of HIV-1 but not HSV-1, indicating MX2’s interference with HIV-1 during capsid MT-directed transport. This correlates with HIV-1 capsids being exposed during MT-traveling, allowing MX2 binding, while HSV-1 capsids rely on tegument proteins for MT motor binding, protecting them against MX2^13,47,48^. MX2’s inhibition of herpesviruses occurs post-entry and after tegument protein dissociation but before viral DNA uncoating and nuclear import, suggesting a cytoplasmic gatekeeper role of MX2 condensates prior to capsid docking to the NPC^8^.

In conclusion, by combining proteomics data, antiviral experiments, and extensive bio-imaging, we demonstrate that intracellular condensate formation is an essential process in MX2-mediated restriction of HIV-1 and HSV-1. Additionally, we describe the composition and properties of MX2 cytoplasmic biomolecular condensates, which was a long-lasting missing piece of MX2 biology. Our findings support a decoy model in which NPC-like cytoplasmic MX2-containing condensates interact with incoming HIV-1 and HSV-1 capsids diverting them away from the NPC for MX2 to trap or disassemble them, respectively, thus preventing the nuclear entry of their viral genome.

## Supporting information

Link to supplementary videos

## Acknowledgements

We are grateful to Dr. Herman Favoreel, Dr. Stacey Efstathiou, Dr. Gary H. Cohen, Dr. Roselyn J. Eisenberg, and Dr. Linos Vandekerckhove for the generous provision of tools including precious antibodies and virus stocks. We would like to thank Dr. Jonathan Maelfait (Ghent University and VIB, Ghent, Belgium) for the helpful discussions. We acknowledge Katie Boucher from the VIB Proteomics Facility (VIB-UGent) for operating the MS instrument, Dr. Linos Vandekerckhove’s team member Zoe Stylianidou for arranging the anti-(HIV-1) p24 clone [*AG3.0]* antibody delivery, and Dr. Chris Boutell (Center for Virus Research, University of Glasgow, U.K.) for teaching on how to prepare clickable HSV-1. G.D.M. was supported by an FWO strategic basic research PhD student fellowship and F.H. by the Hannover Biomedical Research School (HBRS) and the Center for Infection Biology (ZIB). This work was also supported by FWO projects VIREOS (G0H7518N) and VIREOS 2.0 (G0H7322N) to X.S, by the FWO project grant (G042918N) to S.E., by the German Research Council (Deutsche Forschungsgemeinschaft, EXC2155 RESIST 390874280; So 403/6-1; grant ID 443889136; http://www.dfg.de/) to B.S. and by the FWO project grant (G091522N) to Z.D.

## Author contributions

Conceptualization, X.S. and S.E.; Methodology, X.S., S.E., B.S., G.D.M. and F.H.; Investigation, G.D.M., L.D, R.C., F.H., A.B., D.D.S., E.F., A.S.D, A.M., H.G, S.L., and Z.D.; Writing – Original Draft, X.S., S.E., and G.D.M.; Writing – Review & Editing, X.S., S.E., G.D.M., L.D., F.H. and B.S.; Software, G.D.M., L.D. and E.P.; Visualization, G.D.M. and L.D.; Funding Acquisition, X.S., S.E., B.S. and Z.D.; Resources, X.S., S.E., B.S. and Z.D.; Supervision, X.S., S.E. and B.S.

## Inclusion and diversity statement

**-**

## STAR Methods

### Key Resources Table

| REAGENT or RESOURCE | SOURCE | IDENTIFIER |
| --- | --- | --- |
| **Antibodies** |  |  |
| Mouse monoclonal anti-α-tubulin<br>(dilution for immunoblotting 1:1,000) | Sigma-Aldrich | Cat# T5168; RRID: AB_477579 |
| Rabbit polyclonal anti-MX1(MxA)<br>(dilution for immunoblotting 1:5,000) | Novus Biologicals | Cat# H00004599-D01P; RRID: AB_2147716 |
| Rabbit polyclonal anti-MX2<br>(dilution for immunoblotting 1:1,000; dilution for confocal microscopy 1:200) | Sigma-Aldrich | Cat# HPA030235; RRID: AB_10601970 |
| Mouse monoclonal anti-V5-tag<br>(dilution for immunoblotting 1:5,000; dilution for confocal microscopy 1:500) | Thermo Fisher Scientific | Cat# R960-25; RRID: AB_2556564 |
| Rabbit polyclonal anti-FLAG-tag<br>(dilution for immunoblotting 1:1,000) | Sigma-Aldrich | Cat# F7425; RRID: AB_439687 |
| Rabbit polyclonal anti-EGFP<br>(dilution for immunoblotting 1:1,000) | Cell Signaling Technology | Cat# 2555; RRID: AB_10692764 |
| Rabbit polyclonal anti-NC-1 antiserum<br>(dilution for immunoblotting 1:2,000) | Cohen, G. H. et al.<sup>95</sup>; generous gift by Dr. Gary H. Cohen and Dr. Roselyn J. Eisenberg (University of Pennsylvania, Philadelphia, USA) | N/A |
| Mouse monoclonal anti-FLAG-BioM2 antibody | Sigma-Aldrich | Cat# F9291; RRID: AB_439698 |
| Rabbit polyclonal anti-SAMD4A<br>(dilution for immunoblotting 1:200) | Proteintech | Cat# 17387-1-AP; RRID: AB_2301339 |
| Mouse monoclonal anti-TNPO1 Transportin 1 [45]<br>(dilution for immunoblotting 1:1,000) | Abcam | Cat# ab10303, RRID: AB_2206878 |
| Rabbit polyclonal anti-NUP35<br>(dilution for immunoblotting 1:1,000; dilution for confocal microscopy 1:500) | Novus Biologicals | Cat# NB100-93322, RRID: AB_1237255 |
| Mouse monoclonal anti-Nuclear Pore Complex Proteins (FG-Nups) [Mab414]<br>(dilution for immunoblotting 1:2,000 (for NUP62); dilution for confocal microscopy 1:500) | Abcam | Cat# ab24609, RRID: AB_448181 |
| Rabbit monoclonal anti-HA-tag<br>(dilution for confocal microscopy 1:1,500) | Cell Signaling Technology | Cat# 3724 (also 3724S), RRID: AB_1549585 |
| Mouse monoclonal anti-α-tubulin<br>(dilution for confocal microscopy 1:150) | Thermo Fisher Scientific | Cat# MA1-19162; RRID: AB_1070259 |
| Mouse monoclonal anti-HIV-1 p24 [39/5.4A]<br>(dilution for confocal microscopy 1:500) | Abcam | Cat# ab9071; RRID: AB_306981 |
| Mouse monoclonal anti-HIV-1 p24 [AG3.0]<br>(dilution for confocal microscopy 1:250) | Obtained through the NIH HIV Reagent Program, Division of AIDS, NIAID, NIH contributed by Dr. Marie-Claire Gauduin; provided to us by Dr. Linos Vandekerckhove (Ghent University, Ghent, Belgium) | Cat# ARP-4121 |
| Rabbit polyclonal serum anti-HSV-1 capsids<br>(dilution for confocal microscopy 1:500) | Döhner, K. et al.<sup>96</sup>; Serrero, M.C. et al.<sup>13</sup> | N/A |
| Goat anti-Rabbit IgG IRDye 800CW<br>(dilution for immunoblotting 1:5,000) | LI-COR Biosciences | Cat# 926-32211; RRID: AB_621843 |
| Goat anti-Mouse IgG IRDye 800CW<br>(dilution for immunoblotting 1:5,000) | LI-COR Biosciences | Cat# 926-32210; RRID: AB_621842 |
| Goat anti-Rabbit IgG IRDye 680RD<br>(dilution for immunoblotting 1:5,000) | LI-COR Biosciences | Cat# 925-68071; RRID: AB_2721181 |
| Goat anti-Mouse IgG IRDye 680RD<br>(dilution for immunoblotting 1:5,000) | LI-COR Biosciences | Cat# 926-68070; RRID: AB_10956588 |
| Donkey anti-mouse DyLight™ 650<br>(dilution for confocal microscopy 1:1,000) | Thermo Fisher Scientific | Cat# SA5-10169; RRID: AB_2556749 |
| Donkey anti-rabbit Alexa Fluor™ 568<br>(dilution for confocal microscopy 1:500) | Thermo Fisher Scientific | Cat# A10042; RRID: AB_2534017 |
| Donkey anti-rabbit Alexa Fluor™ 488<br>(dilution for confocal microscopy 1:500) | Thermo Fisher Scientific | Cat# A-21206; RRID: AB_2535792 |
| Donkey anti-mouse Alexa Fluor™ 568<br>(dilution for confocal microscopy 1:500) | Thermo Fisher Scientific | Cat# A10037; RRID: AB_2534013 |
| Donkey anti-mouse DyLight™ 488<br>(dilution for confocal microscopy 1:1,000) | Thermo Fisher Scientific | Cat# SA5-10166; RRID: AB_2556746 |
| **Bacterial and virus strains** |  |  |
| HSV-1 strain F | Kindly provided by Dr. Herman Favoreel (Ghent University, Ghent, Belgium) | N/A |
| HSV-1 C12, recombinant HSV-1 strain SC16 containing EGFP expression cassette in the US5 locus | Glauser, D.L. et al.<sup>97</sup>; kindly provided by Dr. Stacey Efstathiou (University of Cambridge, Cambridge, UK) | N/A |
| HSV-1 strain 17<sup>+</sup> Lox-BAC-pHCMV-FireFly Luciferase reporter virus | Khadivjam, B. et al.<sup>98</sup>; | N/A |
| HSV-1 strain 17<sup>+</sup> Lox-wild-type BAC (K33, clone K33) | Sandbaumhüter, M. et al.<sup>99</sup> | N/A |
| HSV-1 strain 17<sup>+</sup> Lox-wild-type BAC (K33, clone K33), with EdC/A labelled genome. | This paper | N/A |
| HIV-1 GFP reporter lentivirus | This paper | N/A |
| HIV IN-EGFP CA WT, single-round viral particles containing fluorescently labelled integrase (IN) | Albanese, A. et al.<sup>100</sup>; Borrenberghs, D. et al.<sup>101</sup> | N/A |
| HIV IN-EGFP CA P90A, single-round viral particles containing fluorescently labelled integrase (IN) | This paper | N/A |
| **Biological samples** |  |  |
| **Chemicals, peptides, and recombinant proteins** |  |  |
| Streptavidin Alexa Fluor™ 680 conjugate | Thermo Fisher Scientific | Cat# S21378 |
| Recombinant human interferon-alpha 2 alpha (rhIFN-a2a) | ImmunoTools | Cat# 11343504 |
| Doxycycline | Sigma-Aldrich | Cat# D9891 |
| Nocodazole | Sigma-Aldrich | Cat# M1404 |
| Biotin | Sigma-Aldrich | Cat# B4639 |
| Sodium arsenite (NaAsO<sub>2</sub>) | Sigma-Aldrich | Cat# S7400 |
| Polybrene infection reagent | Sigma-Aldrich | Cat# TR-1003-G |
| Cycloheximide | Sigma-Aldrich | Cat# C4859 |
| Benzonase | Millipore | Cat# 70746 |
| Trypsin | Promega | Cat# V5111 |
| FuGENE 6 transfection reagent | Promega | Cat# E2691 |
| FuGENE® HD Transfection Reagent | Promega | Cat# E2311 |
| Polyethylenimine (PEI) | Polysciences, Inc. | Cat# 23966-1 |
| DharmaFECT 1 transfection reagent | Horizon Discovery | Cat# T-2001-02 |
| Lipofectamine™ RNAiMAX transfection reagent | Thermo Fisher Scientific | Cat# 13778075 |
| DAPI (4',6-Diamidino-2-Phenylindole, Dihydrochloride) | Thermo Fisher Scientific | Cat# D1306 |
| Nuclear stain NucSpot 650/665 | Biotium | Cat# 41034 |
| Critical commercial assays |  |  |
| NanoBiT® PPI Starter Systems | Promega | Cat# N2014 |
| Nano-Glo® Live Cell Assay System | Promega | Cat# N2011 |
| Luciferase Assay Substrate of the Dual-Luciferase® Reporter Assay System | Promega | Cat# E1910 |
| Deposited data |  |  |
| Proteomics data, MX2 BioID (-IFN, +IFN, HSV-1) | This paper | PRIDE: PXD042544 |
| Experimental models: Cell lines |  |  |
| HEK293T cells | Originate from Lin, Y.-C. et al.<sup>102</sup> | N/A |
| HeLaP4 cells, which express the CD4 receptor | Mondor, I. et al.<sup>103</sup>; Obtained through the National Institutes of Health (NIH) AIDS reagent program | N/A |
| HeLa cells | American Type Culture Collection (ATCC) | CCL-2 |
| U87-MG cells | American Type Culture Collection (ATCC) | HTB-14 |
| U87 CD4+CXCR4+ Cells | Obtained through the NIH HIV Reagent Program, Division of AIDS, NIAID, NIH contributed by Drs. HongKui Deng and Dan Littman | Cat# ARP-4036 |
| Vero cells | American Type Culture Collection (ATCC) | CCL-81 |
| hTERT RPE-1 cells | American Type Culture Collection (ATCC) | CRL-4000 |
| BHK-21 [C-13] cells | American Type Culture Collection (ATCC) | CCL-10 |
| MT-4 cells | Obtained through the NIH HIV Reagent Program, Division of AIDS, NIAID, NIH contributed by Dr. Douglas Richman. | Cat# ARP-120 |
| Experimental models: Organisms/strains |  |  |
| Oligonucleotides |  |  |
| RT qPCR primers, see Supplementary Table 4 | This paper | N/A |
| ON-TARGETplus SMARTpool siRNA information, see Supplementary Table 4 | Horizon Discovery | N/A |
| Ultramer oligo, see Supplementary Table 4 | ID&T | N/A |
| Recombinant DNA |  |  |
| Gene of Interest (GOI) used for GGW cloning sequence information, see Supplementary Table 4 | This paper | N/A |
| Packaging vector gag-pol-rev (pCMV-dR8.74) | Addgene | Cat# 22036 |
| Vesicular stomatitis virus G (VSV-G) envelope expression vector (pMD2.G) | Addgene | Cat# 12259 |
| Expression vector of enhanced green fluorescent protein (EGFP) (pRRLsin.CMV-GFP) | Derived from pRRLsin.cPPT.PGK-GFP.WPRE (Addgene; Cat# 12252)<sup>104</sup> | N/A |
| HIV-1 clone pNL4-3.Luc.R-E<sup>-</sup> | Obtained through the NIH HIV Reagent Program, Division of AIDS, NIAID, NIH contributed by Dr. Nathaniel Landau. | Cat# ARP-3418 |
| HIV-1 clone pNL4-3.Luc.R-E<sup>-</sup> (P90A CA) | This paper | N/A |
| pVpr-IN-EGFP | Albanese, A. et al.<sup>100</sup>; Borrenberghs, D. et al.<sup>101</sup> | N/A |
| HIV-1 strain NL4-3 Infectious Molecular Clone (pNL4-3) | Obtained through the NIH HIV Reagent Program, Division of AIDS, NIAID, NIH: contributed by Dr. M. Martin | Cat# ARP-114 |
| pLV[Exp]-Hygro-EF1a-tTS_rtTA | VectorBuilder | N/A |
| pSpCas9(BB)-2A-Puro (PX459) | Addgene | Cat# 48139 |
| Software and algorithms |  |  |
| MaxQuant software | Cox, J., and Mann, M.<sup>105</sup>; Tyanova, S. et al.<sup>106</sup> | N/A |
| Perseus | Tyanova, S. et al.<sup>107</sup> | N/A |
| FlowJo | FlowJo, LLC | N/A |
| GraphPad Prism | GraphPad Software | N/A |
| g: Profiler | Reimand, J. et al.<sup>108</sup> | N/A |
| Cytoscape | Shannon, P. et al.<sup>109</sup> | N/A |
| SAFE | Baryshnikova, A.<sup>110</sup> | N/A |
| R package ggplot | H. Wickham (2016). ggplot2: Elegant Graphics for Data Analysis. Springer-Verlag New York. | N/A |
| R package enrichR | Jawaid W (2022). enrichR: Provides an R Interface to 'Enrichr'. | N/A |
| Arivis vision 4D | Zeiss | N/A |
| ZEN blue 3.5 | Zeiss | N/A |
| Other |  |  |

### Resource availability

#### Lead contact

Further information and requests for resources and reagents should be directed to and will be fulfilled by the lead contact, Xavier Saelens.

#### Materials availability

The authors declare that the plasmids and cell lines used in this study are available upon reasonable request.

#### Data and code availability

All datasets generated in this study are disclosed as source data. Complete protein lists, plasmid sequences, primers and siRNA information are provided in supplementary tables. The mass spectrometry proteomics data have been deposited to the ProteomeXchange Consortium via the PRIDE^111^ partner repository with the dataset identifier PXD042544.

### Method details

#### Molecular Cloning

All the plasmids used in this study were generated by an in-house BsaI-based Golden Gate assembly cloning system and in-house vector backbones^36^. All coding sequences (including mutant variants) were introduced as entry vectors to support assembly of all the expression constructs in this study. To acquire coding DNA sequence(s) [CDS(s)], complementary DNA (cDNA) derived from U87-MG was amplified by PCR. Alternatively, CDS(s) were obtained as DNA fragments from Twist Biosciences. The resulted sequences of the genes of interest (GOI) were compared with the NCBI available sequences and used if there was a 100% amino acid similarity (GOI sequence information can be found in Supplementary Table.4).

Specifically, for the generation of MX2[NTD]-MX1 and -EGFP chimeric constructs the individual MX2[NTD(1-90 amino acids) and truncated variants] sequences, together with MX1[44-662 amino acids] C-terminally HA tagged sequence and EGFP sequence were introduced as individual entry vectors to the BsaI-based Golden Gate assembly cloning system. The resulting expression constructs produced MX2[NTD] chimeric proteins where between the MX2[NTD] and MX1[44-662 amino acids]-HA or EGFP there is a short (SGGSLQ amino acids) sequence. Our MX2[NTD] chimeric constructs were designed according to ^72^.

Generation of lentiviral vectors was achieved by a Gateway cloning step following the initial Golden Gate assembly reaction to combine two expression cassettes in a lentiviral backbone. The second cassette expresses a selection marker conferring antibiotic resistance for positive selection.

Required sequence mutations, for the removal of BsaI sites in CDSs, for the generation of MX2 mutants and for the generation of pNL4-3.Luc.R^-^E^-^ P90A were performed by fusion PCR.

#### Cells

All cell lines used in this study were cultured in Dulbecco’s modified Eagle medium (DMEM) GlutaMAX supplement (#10566016, Thermo Fisher Scientific), supplemented with 10% fetal bovine serum (FBS, #10270106, Thermo Fisher Scientific), 100 U ml^−1^ penicillin and 100 µg ml^−1^ streptomycin, unless indicated otherwise. Cells were transduced at MOI of 1 for stable introduction of transgenes. For the generation of stable doxycycline-inducible U87-MG (rtTA/tTS _MX2_MX2-T2A-TurboID) and U87-MG (rtTA/tTS _MX2_MX2-MUTT2A-TurboID) cell lines, U87-MG cells were first transduced with Tet-On transactivator (rtTA/tTS) lentivirus and selected with 200 µg ml^−1^ hygromycin (#ant-hg-1, Thermo Fisher Scientific) for 14 days. U87-MG (rtTA/tTS) were subsequently transduced with untagged MX2 expression vectors and selected with 5 µg ml^−1^ blasticidin (#ant-bl-1, Thermo Fisher Scientific) for 14 days. These cells are referred to as U87-MG(MX2). The final U87-MG cell lines used for the BioID experiments were obtained by transducing the U87-MG (rtTA/tTS _MX2) cells with either MX2-T2A-TurboID or MX2-MUTT2A-TurboID lentivirus and selecting with 2 µg ml^−1^ puromycin (#ant-pr-1, Thermo Fisher Scientific) for 7 days. For the generation of stable doxycycline inducible HeLa-MX2 cell lines, HeLa cells were first transduced with rtTA/tTS lentivirus, selected and maintained with 200 µg/ml hygromycin for 14 days, similar to U87-MG. HeLa (rtTA/tTS) were then transduced with untagged MX2(1-715), referred to as HeLa(MX2), V5- or EGFP-tagged MX2(1-715) or MX2 mutants and selected with 2 µg ml^−1^ puromycin.

For the generation of HeLa cells with endogenously tagged (dual V5 and HA tag) MX2 alleles a C-terminal dual V5-HA tag in MX2 was introduced into low passage HeLa cells by nucleofection (Amaxa NucleofectorTM II system, Kit R, Program I-13) of the pSpCas9(BB)-2A-Puro (PX459) vector (sgRNA sequence: ctgaagggcggcgatgcctg) and a single strand oligonucleotide (ID&T ultramer oligo: see Supplementary Table 4). Transfected cells were selected with 2 µg ml^−1^ puromycin for 2 days, and then allowed to recover before immunostaining experiments on the mixed cell population with or without the double V5-HA tag on the C-terminus of MX2.

#### Lentiviral vector production

Lentiviral vector stocks were obtained by transfection of 6.5 x 10^6^ HEK293T cells with packaging vector gag-pol-rev (pCMV-dR8.74), a vesicular stomatitis virus G (VSV-G) envelope expression vector (pMD2.G) and a lentiviral transfer vector containing the transgene in a ratio of 2:1:3 by calcium phosphate transfection. The culture medium was refreshed 1-4 h before transfection and 24 h after transfection. Lentiviral particles-containing supernatants were harvested 48 and 72 h post-transfection, filtered with a 0.45 µm cutoff filter (Millex® filter, #SLHVM33RS, Millipore), concentrated by ultracentrifugation at 22.000 rpm for 2.5 h at 4°C (Optima™ XPN-80 – Beckman Coulter), resuspended in 100 µL of DMEM and stored in small aliquots at −80 °C. For VSV-G.HIV-1-GFP stocks intended for antiviral assays, pRRLsin.CMV-GFP plasmid was derived from (pRRLsin.cPPT.PGK-GFP.WPRE, #12252, Addgene) driving the expression of enhanced green fluorescent protein (EGFP) from a CMV promoter^104^. VSV-G.HIV-1-GFP was titrated by infecting a defined number of HeLa cells and determine the percentage of infected GFP-positive cells 2 days post-transduction by flow cytometry (Attune™ NxT Acoustic Focusing Cytometer, Invitrogen). Single-round viral particles containing fluorescently labelled integrase (IN), referred to as HIV IN-EGFP (wild-type CA or P90A CA), were generated by Vpr-mediated trans-incorporation as described previously^100,101^. Briefly, producer cells (HEK293T) were co-transfected with three plasmids: pVSV-G for the envelope, the HIV-1 clone pNL4-3.Luc.R^-^E^-^ or pNL4-3.Luc.R^-^E^-^ P90A and pVpr-IN-EGFP. The viral molecular clone pNL4-3.Luc.R^-^E^-^ was obtained through the NIH AIDS Research and Reagent Reference program^112^. Upon maturation, the HIV protease cleavage leads to the production of fluorescent IN-EGFP that is incorporated in the viral particle. Viruses were harvested 48 hours after transfection, filtered and concentrated as described above. For pNL4-3.HIV-1 virus intended for analysis of HIV-1 capsids intracellular localization in_IFN-stimulated U87 CD4+CXCR4+ cells: The viral molecular clone pNL4-3 was used to produce wild type HIV-1. The NL4-3.HIV-1 virus was produced as described before^113^.Briefly, HEK293T cells were transfected with pNL4-3 plasmid and the virus was harvested in supernatant after 72 hours. The produced virus was used for the infection of MT-4 cells grown in RPMI 1640 medium supplemented with 12% FCS and 50 μg/ml gentamicin (#15750060, Thermo Fisher Scientific) for scaling up the viral production. The supernatant was collected by centrifugation and quantified for viruses by measuring HIV-1 capsid (p24) protein by ELISA (INNOTEST p24-ELISA, Innogenetics).

#### HSV-1 Viruses

Herpes simplex virus-1 (HSV-1) stocks of HSV-1 strain F, HSV-1 strain C12 and HSV-1 strain 17+ Lox-BAC-pHCMV-FireFly Luciferase reporter virus were grown and titrated on Vero cells as described by Grosche et al^114^. HSV-1 strain 17+ Lox-wild-type BAC (K33, clone K33) stocks for intracellular viral capsid microscopy experiments were gradient-purified to remove residual cell debris and cytokines as described by Grosche et al^114^ using Ficoll® 400 (#F4375, Sigma-Aldrich) gradient solution. EdC/A pre-labelled HSV-1 virions were prepared and purified as described before^114^ while pulsed with EdC/A as established previously^115^. Briefly, RPE cells were seeded in 175 cm^2^ flasks, cultured to 80 - 90% confluency, washed with PBS, and infected with 0.005 plaque-forming units (PFU) per cell of an HSV-1 strain 17+ Lox-wild-type BAC (K33, clone K33) virus stock derived from BHK cells. The virus was diluted in 5 mL CO2-independent medium (#18045-070, Life technologies Gibco) containing 0.1% bovine serum albumin (BSA, PAA Laboratories) and cells inoculated on a rocking platform for 1 hr at RT. Then, 21 mL of culture medium containing 0.2% FCS were added, and cells were incubated at 37°C and 5% CO2. After 14 h, 40 h, and 64 h cells were pulsed with 0.5 µM 5-Ethynyl-2’-deoxycytidine (EdA, #CLK-N003-10, Jena Bioscience) and 0.5 µM 7-Deaza-7-ethynyl-2’-deoxyadenosine (EdA, #CLK-099, Jena Bioscience). When nearly all cells had rounded up (ca. 75h), the supernatant and detached cells were harvested. Cell debris was pelleted by centrifugation for 10 min at 4,000 rpm and 4°C (JA-10 rotor, Avanti J-25 centrifuge, Beckman Coulter). To concentrate only the extracellular virions, the supernatant was centrifuged for 25 min at 20,000 rpm and 4°C (Beckman rotor SW32, Beckman Coulter). The pellets were gently resuspended in 600 to 1000 µL of MNT buffer (30 mM 2-(N-morpholino)ethanesulfonic acid, 100 mM NaCl, 20 mM Tris-HCl, pH = 7.4) and allowed to swell. After 24 h, the suspension was further homogenized by repeated pipetting, and the medium pellet virus preparation was aliquoted, snap frozen, and stored at -80 °C.

#### MX2 Proximity Biotinylation (BioID) for LC-MS/MS

MX2 BioID experiments were performed in three conditions: (i) doxycycline-treated (-IFN), (ii) doxycycline-treated combined with IFNa2a stimulation (+IFN), and (iii) doxycycline-treated and infected with HSV-1 (HSV-1). All three BioID experiments were performed simultaneously using same passage U87-MG (rtTA/tTS_MX2_MX2-T2A-TurboID) and U87-MG (rtTA/tTS _MX2_MX2-MUTT2A-TurboID) cell lines. For every condition, four control samples (T2A) and four MX2 specific samples (MUTT2A) were prepared. For every sample a 145 cm^2^ culture dish (Nunc) was seeded with 5 × 10^6^ cells. Twenty-four hours after seeding, 15 ng ml^−1^ of doxycycline was added to each sample. For the +IFN condition samples the medium was also supplemented with 1,000 IU ml^−1^ rhIFN-a2a. Twenty-four hours after doxycycline stimulation, biotin was added to a final concentration of 50 μM to every culture dish, with the HSV-1 condition samples first challenged with HSV-1 strain F at MOI of 20. The virus was first inoculated on ice at 4°C for 2 h and then samples were washed with PBS and incubated with biotin containing medium at 37°C and 5% CO_2_. Biotinylation was performed for 2 h for all the samples. Then the growth medium was aspirated and cells were washed two times with ice-cold PBS before were lysed in 750 μL RIPA lysis buffer (50 mM Tris-HCl pH 7.5, 150 mM NaCl, 1% NP-40, 1 mM EDTA, 1 mM EGTA, 0.1% SDS, 1X EDTA-free protease inhibitor cocktail [#04693132001, Roche], and 0.5% sodium deoxycholate) and collected using a cell scraper. Lysates were incubated with agitation for 1 h at 4 °C with 250 IU of benzonase followed by 15 min centrifugation at 15,000 x g at 4°C to remove the insoluble fraction of the lysate. Supernatants were transferred to a new tube and total protein concentration was determined by the Bradford protein assay to normalize input material to the maximum shared total protein amount across the samples between all three conditions. For every sample, 30 μL of streptavidin Sepharose high-performance bead suspension (#GE17-5113-01, Cytiva) was used for the enrichment of biotinylated proteins. Beads were pelleted by centrifugation at 500 x g for 1 min and washed three times in RIPA lysis buffer prior to addition to the lysate. Affinity purification was performed on a rotator at 4°C for 3 h. Beads were recovered by centrifugation of the samples at 500 x g for 1 min at 4°C and aspirating the supernatant. Beads were washed three times in RIPA lysis buffer, twice in 50 mM ammonium bicarbonate pH 8.0, and once in 20 mM Tris-HCl pH 8.0, 2 mM CaCl2 prior to resuspension in 20 μL of 20 mM Tris-HCl pH 8.0. Trypsin digest was allowed overnight at 37°C by the addition of 1 μg of trypsin (#V5111, Promega) to each sample. Peptide mixtures were depleted of beads by centrifugation at 500 x g for 1 min, after which the supernatant was transferred to a new Eppendorf tube. To ensure complete digestion, peptide mixtures were incubated with an additional 500 ng of trypsin for 3 h before acidification with formic acid to a final concentration of 2%, followed by centrifugation at 15,000 x g at room temperature (RT). Peptides containing supernatants were transferred in mass spectrometry vials (#9301-0978, Agilent).

#### LC-MS/MS

Purified peptides were re-dissolved in 21.5 µL solvent A of which 2.5 µL was injected for LC-MS/MS analysis on an Ultimate 3000 RSLCnano system (Thermo Fisher Scientific) in line connected to a Q Exactive HF Biopharma mass spectrometer (Thermo Fisher Scientific) equipped with a pneu-Nimbus dual ion source (Phoenix S&T). Trapping was performed at 10 μL/min for 4 in loading solvent A on a 20 mm trapping column (made in-house, 100 μm inner diameter (ID), 5 μm beads, C18 Reprosil-HD, Dr. Maisch). The peptides were separated on a 250 mm nanoEase column (M/Z HSS T3 column, 100 Å, 1.8 μm, 75 μm I.D., Waters) kept at a constant temperature of 40°C. Peptides were eluted by a non-linear increase from 1 to 55% MS solvent B (0.1% FA in water/ACN (2:8, v/v)) over 75 min, at a flow rate of 300 nL/min, followed by a 15 min wash reaching 97% MS solvent B and re-equilibration with 99% MS solvent A (0.1% FA in water). The mass spectrometer was operated in data-dependent mode, automatically switching between MS and MS/MS acquisition for the 12 most abundant ion peaks per MS spectrum. Full-scan MS spectra (375-1,500 m/z) were acquired at a resolution of 60,000 in the Orbitrap analyzer after accumulation to a target value of 3,000,000. The 12 most intense ions above a threshold value of 13,000 (minimum AGC of 1,000) were isolated for fragmentation at a normalized collision energy of 30%. The C-trap was filled at a target value of 100,000 for maximum 80 ms and the MS/MS spectra (200-2,000 m/z) were acquired at a resolution of 15,000 in the Orbitrap analyzer with a fixed first mass of 145 m/z. Only peptides with charge states ranging from +2 to +6 were included for fragmentation and the dynamic exclusion was set to 12 sec.

#### Mass Spectrometry Data Processing and Analysis

Data analysis of the Obtained Xcalibur raw files (.raw) was performed using MaxQuant (version 1.6.11.0) using the Andromeda search engine with default search settings. PSM, protein, and site false discovery rates were set at 0.01. Spectra were searched against all human proteins in the Uniprot/Swiss-Prot database (database release version of January 2019 containing 20,413 protein sequences) (taxonomy ID 9606) and human herpesvirus 1 (strain 17) viral proteins taxonomy (HHV-1) (Human herpes simplex virus 1) (proteome ID UP000009294), including a FASTA file for the N-terminally V5 tagged TurboID sequence.

Enzyme specificity was set as C-terminal to arginine and lysine (trypsin), also allowing cleavage at arginine/lysine-proline bonds with a maximum of two missed cleavages. Methionine oxidation and N-terminal acetylation were set as variable modifications. Matching between runs was enabled with an alignment time window of 20 min and a matching time window of 0,7 min. Only proteins with at least one peptide were retained to compile a list of identified proteins. Proteins were quantified by the MaxLFQ algorithm integrated in the MaxQuant software. A minimum of two ratio counts from at least one unique peptide was required for quantification and the Fast LFQ option was disabled. The minimum peptide length was set at seven amino acids, and a maximum peptide mass of 4,600 Da was used.

Further data analysis was performed with the Perseus software (version 2.0.3.1). For each MX2 BioID condition, LFQ intensities of four control samples (T2A) and four MX2 specific samples (MUTT2A) were imported as initial matrix with the exemption of +IFN control samples (three samples). Hits identified in the reverse database, only identified by modification site and contaminants were removed and protein LFQ intensities were log2 transformed. Replicate samples were grouped, proteins with less than four valid values in at least one group were removed and missing values were imputed from a normal distribution with a width of 0.3 and a down shift of 1.8 to compile a list of quantified proteins. 2318 proteins were quantified. On the quantified proteins, for each condition a t-test was performed for a pairwise comparison between the control (T2A) and MX2 specific (MUTT2A) samples to reveal specific MX2 interaction partners. Number of randomizations was set to 1,000, FDR to 0.01 and S0 to 1. The results of these t-tests are shown in the volcano plots in Fig. 1 visualized using RStudio also showing proteins identified with an FDR 0.05. For each differential analysis, the log2 (T2A/MUTT2A) fold change value is indicated on the X-axis, while the statistical significance (−log10 p-value) is indicated on the Y-axis. Proteins outside the curved lines, set by an FDR value of 0.01 or 0.05 and an S0 value of 1, represent specific MX2 interaction partners.

#### MX2 SAFE network

The MX2 SAFE network was generated by the FDR 0.01 enriched common MX2 interactors between the three MX2 BioID conditions (-IFN, +IFN, HSV-1). Using Cytoscape (version 3.6.1), the complete BioGRID Protein-Protein Interactions network (*H. sapiens*) was uploaded and the 230 commonly enriched proteins were used to filter the nodes and edges to build the MX2 network as edge-weighted spring-embedded layout. SAFE is an automated network annotation tool that requires annotations for network nodes. Using g:Profiler we created an attribute file list of all known GO (Gene Ontology) cellular component terms assigned to each node of the MX2 network that we coupled to SAFE reaching 98% network coverage. SAFE analysis was performed assuming edges of the network are undirected, map-weighted distance, with 15 percentile threshold (manual assessment suggested that a value in this range should be suitable). SAFE generates an enrichment score for each node according to evidence of association to a specific GO cellular component term and assigns a color according to that score generating separate clusters (“domains” or ‘’landscapes’’ in SAFE terminology) with a known GO term. To build a composite map, a minimum of 5 nodes was set as minimum landscape size. The final MX2 SAFE network was generated by visualizing the 4 main (umbrella) landscapes resulted from the analysis and export the resulting SAFE network as PNG image. Each node represents a protein in the network, colored according to the SAFE enrichment score matching evidence of association to a specific Gene Ontology (GO) cellular component term.

#### Protein network visualization and Dotplot generation

Following SAFE analysis in Cytoscape, nodes and edges clustering to individual SAFE domains of the MX2 SAFE network were selected and visualized as separate protein networks maintaining the same layout. The color of the nodes for each protein network was selected according to the SAFE assigned GO-specific color. For the nuclear pore protein network the nodes layout was manually organized at specific protein sub-clusters and subcomplexes according the NCP organization described by^94^.

For the proteins of each network, dotplot analysis was generated using RStudio with the size and color of each dot representing the protein enrichment (log2(FC)) and statistical significance (−log10 p-value), respectively. Proteins are ranked according to their average enrichment (log2(FC)) in all three BioID conditions.

#### SDS-PAGE and immunoblotting

Cells were lysed in 2X Laëmmli buffer (125 mM Tris-HCl pH 6.8, 4% SDS, 20% glycerol, 0.004% Bromophenol blue supplemented with 20 mM dithiothreitol [DTT]), in TAP lysis buffer^45^ (50 mM HEPES-KOH [pH 8.0], 100 mM KCl, 2 mM EDTA, 0.1% NP-40, 10% glycerol) or in RIPA lysis buffer (50 mM Tris-HCl pH 7.5, 150 mM NaCl, 1% NP-40, 1 mM EDTA, 1 mM EGTA, 0.1% SDS) both supplemented with 1X EDTA-free protease inhibitor cocktail (#04693132001, Roche). For the two latter lysis conditions lysates were complemented with XT sample buffer (#1610791, Bio-Rad) and XT reducing agent (#1610792, Bio-Rad) to a final volume of 30 μL.

Protein samples were boiled for 10 min at 95 °C prior to SDS-PAGE. Samples were loaded on 4–20% polyacrylamide gradient gels (#M42015, GenScript), according to the guidelines of the manufacturer. Proteins were transferred to PVDF membrane (#IPFL00010, Merck) for 3 h at 60 V or for 30 minutes at 100 V in blotting buffer (48 mM Tris, 39 mM glycine, 0.0375 % SDS and 20 % methanol). Membranes were blocked for 1 h at room temperature (RT) with blocking buffer (#927-50000, LI-COR) and incubated with primary antibodies overnight at 4°C in blocking buffer. The next day, membranes were washed four times for 10 min with TBS-Tween20 0.1% (v/v) buffer (TBS-T) and further incubated at RT for 1 h with the appropriate secondary antibody in blocking buffer. Membranes were washed four times for 10 min with TBS-T. Immunoreactive bands were visualized on a LI-COR-Odyssey infrared scanner (Li-COR).

#### Co-immunoprecipitation

Coimmunoprecipitations were performed to confirm the interaction between MX2-TurboID fusion protein with FLAG-tagged MX2. Eight million HEK293T cells were seeded in 10-cm tissue culture dishes. The following day, cells were transfected using polyethylenimine (PEI) with 5 μg of the indicated DNA constructs using a 1:5 ratio with PEI. Twenty-four hours after transfection, cells were lysed with TAP lysis buffer supplemented with 1 mM dithiothreitol [DTT], 0.5 mM phenylmethylsulfonyl fluoride (PMSF), 0.25 mM sodium orthovanadate, 50 mM B-glycerophosphate, 10 mM NaF, and 1X EDTA-free protease inhibitor cocktail. Lysates were incubated with 1 μg anti-FLAG-BioM2 antibody or without antibody for 3 h at 4°C under constant rotation. Afterwards protein complexes were immunoprecipitated by the addition of 10 µL of magnetic Dynabeads™ MyOne™ Streptavidin T1 (#65601, Thermo Fisher Scientific) for 10 min at RT under constant rotation. Beads were captured in magnet and washed 2 times with lysis buffer followed by protein elution directly in 1X loading buffer. Proteins that bound to the beads were detected by immunoblotting.

#### DNA plasmid transfection

Plasmid transfection for the validation of untagged MX2 and MX2-T2A or MUTT2A-TurboID expression vectors: For evaluation of protein expression (Figure S1A), 1 million HEK293T cells were seeded into 6-well plates and transfected using polyethylenimine (PEI) with 1 μg of the indicated DNA constructs at a 1:5 ratio with PEI. For HSV-1 C12 strain transient expression antiviral experiments (Figure S1C), 10,000 HeLa cells were seeded into 96-well black/clear bottom plates and transfected using PEI with 70 ng of the indicated DNA constructs at a 1:5 ratio with PEI. For VSV-G.HIV-1-GFP transient expression antiviral experiments (Figure S1D), 35,000 HeLa cells were seeded into 48-well plates and transfected using PEI with 300ng of the indicated DNA constructs.

Plasmid transfection for evaluation of fluorescently tagged MX2 expression vectors (Figure S4A): For both VSV-G.HIV-1-GFP 2-LTR cycles measurement and HSV-1 firefly strain transient expression antiviral experiments, 35,000 HeLa cells were seeded into 48-well plates and transfected using PEI with 300ng of the indicated DNA constructs.

Plasmid transfection for transient expression microscopy experiments, 15,000 HeLa-derived cells were seeded into 8-well imaging µ-Slide (#80826, Ibidi GmbH) and transfected using FUGENE HD with 100 ng of DNA at 1:3 ratio with FUGENE HD.

#### siRNA-mediated knockdown

For HeLa and doxycycline-inducible HeLa(MX2), EGFP- or V5-tagged HeLa(1-715) and MX2 mutants: For the RNA interference (RNAi) based imaging (Figure 5I) and functional experiments using HSV-1 firefly and VSV-G.HIV-1-GFP reporter viruses, 15,000 HeLa based cells were seeded in 48-well plates and transfected, with 15 nM (single gene target) or 30 nM (two gene targets or single gene target with Non-Targeting or MX2 siRNA, 15 nM each) of SMARTpool siRNAs (Dharmacon) (SMARTpool siRNAs gene specific catalog numbers can be found in Supplementary Table.4) according to manufacturer’s instructions using DharmaFECT 1 transfection reagent in antibiotic free medium. Six hours post-transfection the medium was refreshed with complete culture medium complemented or not with doxycycline. For experiments with HeLa cells stimulated with IFN, the culture medium was refreshed 24 h post-transfection with medium complemented or not with IFNa2a. For the RNA interference (RNAi) based functional experiments using HSV-1 C12 strain GFP reporter virus, 5,000 HeLa based cells were seeded in 96-well black/clear bottom plates and transfected as described above. For HeLa and HeLa based cells, cell lysates to document protein knockdown were collected 48 hours post-transfection for all experiments regardless type of cell stimulation (doxycycline or IFNa2a).

For U87-MG and doxycycline-inducible U87-MG(MX2): For the RNA interference (RNAi) based functional experiments using HSV-1 C12 strain GFP reporter virus and VSV-G.HIV-1-GFP reporter viruses, 8,500 U87-MG based cells were seeded in 96-well plates and transfected twice, 24 h apart with 10 nM (single gene target) or 20 nM (single gene target with Non-Targeting or MX2 siRNA, 10 nM each) of SMARTpool siRNAs (Dharmacon) (SMARTpool siRNAs gene specific catalog numbers can be found in Supplementary Table.4) according to manufacturer’s instructions using Lipofectamine™ RNAiMAX transfection reagent in antibiotic free medium. Seven hours after the second transfection the medium was refreshed with complete culture medium complemented or not with doxycycline or IFNa2a. For U87-MG(MX2), cell lysates to document protein knockdown were collected 48 hours post-transfection/doxycycline-stimulation (Figure 2D and E) and for U87-MG, cell lysates to document protein knockdown were collected 24 hours post-transfection/IFN-stimulation (Figure 2H).

#### RT-qPCR for siRNA-mediated knockdown efficiency validation

cDNA synthesis was performed using the PrimeScript RT kit (#RR037A, Takara Bio) using 250 ng RNA input from siRNA transfected HeLa(MX2) cells, collected 24 h after transfection. TNPO1, TNPO2, SAMD4A, SAMD4B, NUP58, NUP62, NUP35, NUP54, TRIM26, TRIM56, FUBP3, and FAM120C specific primer pairs were designed using the PrimerQuest design tool (IDT, primers list can be found in Supplementary Table.4). Target transcripts were amplified with the SensiFAST SYBR No-ROX kit (#BIO-98005, Meridian Bioscience) according to the manufacturer’s instructions and signal was detected using a LightCycler 480 System (Roche). Samples were measured in technical duplicates in a 384-well plate. The following cycling conditions were used: 1 cycle at 95°C for 5 min, 40 cycles at 95°C for 10 s, 60°C for 10 sec and 72°C for 10 sec, followed by melting curve analysis to validate unique amplicons. Quantitation cycles (Cq) were normalized to housekeeping genes (SDHA, UBC, YWHAZ) using geometric averaging. Normalized Cq values were subsequently normalized relative to the Non-Targeting siRNA sample. All RT-qPCR analyses were done in qbase+ (CellCarta), the geNorm algorithm was used to ensure stability of the housekeeping genes across all samples^116^. Visualization of the data was performed in GraphPad Prism.

#### qPCR to detect 2-LTR circles for VSV-G.HIV-1-infected cells

DNA was extracted using QIAamp DNA Micro kit (#56304, Qiagen). DNA yields were quantified with a NanoPhotometer (Implen). Quantitative PCRs were performed in technical duplicates for each sample with an input of 30 ng DNA. Target sequences were amplified with the SensiFAST SYBR No-ROX kit (#BIO-98005, Meridian Bioscience) according to the manufacturer’s instructions. Primers for the quantification of 2-LTR circles are listed in Supplementary Table 4. Cycling was performed for 1 cycle at 95°C for 5 min, 40 cycles at 95°C for 10 s, 60°C for 10 s and 72°C for 10 s, followed by melting curve analysis to validate unique amplicons. Cq values were normalized to CDKN1A and ACTB housekeeping genes using geometric averaging, and DCq values were normalized to the non-targeting siRNA control. All RT-qPCR analyses were done in qBase+ (CellCarta).

#### In vitro infection with Herpes Simplex Virus Type 1 strain C12 GFP reporter virus (HSV-1 C12)

Twenty-four hours after siRNA/DNA transfection and stimulation [doxycycline and IFNa2a (1,000 IU mL^-1^)] of HeLa- and U87-MG-derived cells, seeded in 96-well black/clear bottom plates, cells were challenged with HSV-1 C12 at MOI of 0.05.

Infection was maintained in 2% FBS FluoroBrite DMEM medium (#A18967-01, Thermo Fisher Scientific), supplemented with/out 10 ng mL^-1^ doxycycline when doxycycline-inducible HeLa(MX2) or U87-MG(MX2) were used. GFP fluorescence was monitored at 24 h intervals using the Infinite® 200 PRO Reader and Tecan iControl Software (Tecan Life Sciences). Each individual measurement (4 biological replicates) was normalized to the mean value of the uninfected wells (4 biological replicates). The relative area under the curve (AUC) from 0 to 48h (U87-MG-derived cells) or 72 h (HeLa-derived cells) post-infection was calculated for each growth curve. For proteins expression analysis, 48 h post-infection cells from a single well were washed with ice-cold PBS and collected in 2X Laëmmli buffer. Protein expression was analysed by immunoblot using anti-MX2, anti-tubulin, and anti-NC-1(VP5) antibodies.

#### In vitro infection with Herpes Simplex Virus-1 (HSV-1) strain 17^+^ Lox-BAC-pHCMV-FireFly <u>Luciferase reporter virus (HSV-1 firefly)</u>

Forty-eight hours after siRNA transfection/doxycycline stimulation of HeLa(MX2) cells or 24 h after DNA transfection of HeLa cells, seeded in 48-well plates, cells were challenged with HSV-1 firefly reporter virus at MOI of 1. Infection was maintained in 2% FBS DMEM medium supplemented with/out doxycycline. Six and a half hours post-infection cells were lysed in 50 µL of 1X Passive Lysis Buffer and luminescence was determined using the Luciferase Assay Substrate of the Dual-Luciferase® Reporter Assay System, measured on a plate reader (GloMax® Navigator Microplate Luminometer, Promega). For proteins expression analysis, the same lysates were used. Protein expression was analysed by immunoblot using anti-MX2, and anti-tubulin antibodies.

#### In vitro infection with HIV-1 based lentiviral vector expressing EGFP (VSV-G.HIV-1-GFP)

Forty-eight hours after siRNA transfection/doxycycline stimulation of HeLa-derived cells or 48h after siRNA transfection (24 h after IFNa2a stimulation with 1,000 IU mL^-1^) of HeLa cells or 24 h after DNA transfection of HeLa cells, seeded in 48-well plates cells were challenged with VSV-G.HIV-1-GFP reporter virus.

Forty-eight hours after siRNA transfection/doxycycline stimulation of U87-MG(MX2) cells or 24 h after siRNA transfection/IFNa2a stimulation with 1,000 IU mL^-1^ of U87-MG cells seeded in well 96-well plates, cells were challenged with VSV-G.HIV-1-GFP reporter virus.

Infection was maintained in 10% FBS DMEM supplemented with 8 µg ml^-1^ polybrene infection reagent (#TR-1003-G, Sigma-Aldrich). Twenty-four hours post-infection cells were washed with PBS and the medium was refreshed with complete culture medium. The percentage of infected GFP-positive cells was determined 48 h post-transduction by flow cytometry (Attune™ NxT Acoustic Focusing Cytometer, Invitrogen), following cell detachment and fixation with 4% paraformaldehyde (PFA). For HeLa and U87MG IFN experiments proteins expression analysis, 24 h post-IFN stimulation (same time-point as the virus inoculation) cells from a single well were washed with ice-cold PBS and collected in RIPA lysis buffer and 2X Laëmmli buffer, respectively. Protein expression was analysed by immunoblot using anti-MX2, and anti-tubulin antibodies.

#### Analysis of HSV-1 capsids intracellular localization

HeLa (rtTA/tTS-Control/NoMX2) cells and doxycycline-inducible HeLa cells expressing MX2(1-715) or mutant variants V5-tagged were treated with different doxycycline amounts to achieve comparable expression levels. Twenty-four hours after doxycycline stimulation were inoculated with gradient purified HSV-1 strain 17+ Lox -wild-type BAC virus stock (MOI:50) in RPMI 1640 medium, GlutaMAX™ supplement (#61870010, Thermo Fisher Scientific) containing 0.2% (w/v) Bovine Serum Albumin (BSA) and 20 mM HEPES for 2 h on ice to allow virus binding. After 2 h, cells were washed twice with RPMI 1640 medium, and incubated for 2.5 h at 37°C and 5% CO_2_ in 2% FBS DMEM supplemented with 100 µg ml^-1^ cycloheximide (#C4859, Sigma-Aldrich) to prevent viral proteins synthesis. Subsequently, HSV-1 infected cells were fixed using 3% paraformaldehyde (PFA) in PBS for 10 min at RT and then permeabilized with 0.2% Triton X-100 in PBS for 10 min at RT. Detection of intracellular incoming capsids was achieved by staining the infected cells with pre-absorbed anti-capsids SY4563 antibody. Briefly, SY4563 antibody was diluted 1:100 in blocking buffer and pre-absorbed with fixed and permeabilized HeLa cells in a 6-well plate with constant rocking overnight (O/N) at 4°C. The next day, the pre-absorbed antibody was collected in an Eppendorf tube and centrifuged for 5 min at 15,000 x g at RT. Antibody containing supernatant was collected and diluted in a final volume of 1:500 in blocking buffer.

#### Analysis of HIV-1 capsids intracellular localization

For VSV-G.HIV-1 GFP virus capsid intracellular localization in HeLa-derived cells: HeLa cells expressing MX2(1-715) C-terminally tagged with EGFP, HeLa (rtTA/tTS-Control/NoMX2) cells and doxycycline-inducible HeLa cells expressing MX2(1-715) or mutant variants EGFP-tagged were treated with different doxycycline amounts to achieve comparable expression levels. Twenty-four hours after DNA transfection/doxycycline stimulation cells were inoculated with filtered (0.45 µm, 33 mm Millex® filter, #SLHVM33RS, Millipore) VSV-G.HIV-1-GFP lentiviral particles (MOI:25) for 1 h on ice in culture medium in the presence of 100 µg ml^-1^ cycloheximide and 8 µg ml^-1^ polybrene infection reagent and then placed at at 37°C and 5% CO_2_. After 6 h, cells were fixed using 3% paraformaldehyde (PFA) in PBS for 10 min at RT and then permeabilized with 0.2% Triton X-100 in PBS for 10 min at RT. Detection of intracellular incoming capsids was achieved by staining the infected cells with mouse monoclonal anti-(HIV-1) p24 clone [39/5.4A] or clone [AG3.0] antibodies.

For pNL4-3.HIV-1 virus capsid intracellular localization in IFN-stimulated U87 CD4+CXCR4+ cells: One day prior to infection, 50,000 U87 CD4+CXCR4+ cells were seeded in 8-well imaging µ-Slide. The next day cells were left untreated or treated with 1,000 IU ml−1 of rhIFN-a2a. Twenty-four hours after IFN stimulation cells were infected with 8 × 10^5^ pg (0,8 ug) of p24 replication-competent NL4-3.HIV-1. After 6 hours, the cells were washed with PBS and fixed in 4% paraformaldehyde (PFA) in PBS for 20 min at RT and then permeabilized with 0.2% Triton X-100 in PBS for 10 min at RT. Detection of intracellular incoming capsids was achieved by staining the infected cells with mouse monoclonal anti-(HIV-1) p24 clone [AG3.0] and anti-MX2 specific antibody.

#### Analysis of HIV IN-EGFP particles intracellular localization

For the analysis of HIV IN-EGFP (wild-type capsid) intracellular localisation in HeLaP4 cells: One day prior to infection, 30,000 HeLaP4 cells were seeded into 8-well imaging µ-Slide (#80826, Ibidi GmbH) that was previously coated with 0.1 mg ml^-1^ Poly-D-lysine hydrobromide (#P6407, Sigma-Aldrich). The following day, cells were transfected with MX2-mScarlet vector using FuGENE 6 transfection reagent following the manufacturers protocol. The cells were maintained in 10% FBS DMEM at 37°C and 5% CO_2_. After 24 hours, when the MX2-mScarlet protein was expressed, the cells were infected with IN-EGFP VSV-G pseudotyped HIV-1 virus (HIV IN-EGFP) with an amount corresponding to 4 µg of p24 antigen. After 6 hours, the cells were washed with PBS and fixed in 4% paraformaldehyde (PFA) in PBS for 15 min at RT. After fixation, the cells were treated with 1 µg ml^-1^ nuclear stain NucSpot 650/665 for 30 min at RT. The cells were washed again and stored in PBS at 4°C until imaging.

For the analysis of HIV IN-EGFP (wild-type capsid and P90A capsid) intracellular localisation in doxycycline-inducible V5-tagged MX2(1-715) HeLa cells: One day prior to infection, 30,000 doxycycline-inducible V5-tagged MX2(1-715) HeLa cells were seeded into 8-well imaging µ-Slide (#80826, Ibidi GmbH) that was previously coated with 0.1 mg ml^-1^ Poly-D-lysine hydrobromide (#P6407, Sigma-Aldrich) in culture medium in the presence of 200 ng ml^−1^ doxycycline. After 24 hours, when the MX2-V5 protein was expressed, the cells were infected with either IN-EGFP VSV-G pseudotyped HIV-1 virus (HIV IN-EGFP) or HIV IN-EGFP P90A with an amount corresponding to 4 µg of p24 antigen for each virus. After 6 hours, the cells were washed with PBS and fixed in 4% paraformaldehyde (PFA) in PBS for 15 min at RT and then permeabilized with 0.2% Triton X-100 in PBS for 10 min at RT. Detection of MX2-V5 was achieved by staining the infected cells with mouse anti-V5-tag antibody. After staining, the cells were treated with 1 µg ml^-1^ nuclear stain NucSpot 650/665 for 30 min at RT. The cells were washed again and stored in PBS at 4°C until imaging.

For the EGFP-IN PICs visualization/quantification: At some images of infected cells the viral inoculum generated EGFP clusters at the outside of the cells which were segmented, and excluded from the quantitation [see <u>Quantification in Arivis vision 4D version 4.0.0 (Carl Zeiss)</u> section].

#### CLICK experiments for the analysis of intracellular EdC/A labelled HSV-1 genomes

HeLa (rtTA/tTS-Control/NoMX2) cells and doxycycline-inducible HeLa cells expressing V5-tagged MX2(1-715) or mutant variants were cultured on 12mm coverslips for 16h until treated with different doxycycline amounts to achieve comparable expression levels. Twenty-four hours after doxycycline stimulation, cells were synchronously infected as described previously^68^. Briefly, cells were inoculated with MOI of 25 pfu/cell corresponding to 1.25 x 10^7^ pfu/mL for 1.5h on ice on a rocking platform to facilitate virus binding. Afterwards, cells were washed twice with ice-cold PBS and growth medium containing cycloheximide was added. Virus internalization was initiated by shifting to 37°C and 5%CO_2_ for 2h. Cells were washed twice with CSK buffer (10mM Hepes, pH 7.0, 100mM NaCl, 300mM Sucrose, 3mM MgCl2, 5mM EGTA (pH 8.0)), then fixed and permeabilized with 2% (w/v) paraformaldehyde, and 0.5% (v/v) Triton X-100 in CSK buffer for 10 min and subsequently washed twice with CSK buffer as established by Alandijany et al^115^. The residual paraformaldehyde was quenched with 50 mM NH_4_Cl for 10 min. To block unspecific protein interactions, the samples were incubated with 2% (v/v) serum from a healthy HSV-1 seronegative volunteer in PBS overnight at 4°C. The samples were washed once with PBS and then the EdC/A present in the virus’ genomes was coupled to Alexa-Fluor-488-Picolylazide (#CLK-1276-1, Jena Bioscience) in a cupper-catalysed CLICK reaction using the Click-iT® Plus Alexa Fluor® Picolyl Azide Toolkit (#C10643, Thermo Fisher). Afterwards they were probed with primary and secondary antibodies diluted in blocking solution. DNA was stained by 0.05 µg/mL 4′,6-diamidino-2-phenylindole (DAPI) in PBS with 0.05% (v/v) DMSO, 0.0005% (v/v) NP-40, 0.025% (w/v) bovine serum albumin, 0.05 mM Tris-HCl (pH 7.4), 0.73 mM NaCl, 0.01 mM CaCl_2_, and 0.11 mM MgCl_2_. Coverslips were embedded using Immunoselect Antifading Mounting Medium (#SCR-038447, Dianova).

For the HSV-1 genome visualization/quantification: A minor proportion of cytosolic viral genomes was also observed in every condition, which were often in close proximity to HSV-1 capsids. These viral particles likely represented defective particles from the inoculum^117^. Under all conditions, even w/o the addition of virions, there were some background signals from the CLICK reaction in the nucleoli and the cytoplasm which were segmented, and excluded from the quantitation [see Figure 6G and S7F and <u>Quantification in Arivis vision 4D</u> <u>version 4.0.0 (Carl Zeiss</u>) section].

#### Nocodazole treatment

For antiviral assays, 15,000 HeLa MX2 cells were grown with/out doxycycline in 48-well plates. Twenty-four hours after stimulation medium was refreshed with DMSO or 1, 5 or 10 μM of nocodazole supplemented culture medium. Three hours after nocodazole treatment cells were infected in the presence similar DMSO or 1, 5 or 10 μM of nocodazole conditions. For HSV-1 firefly reported virus challenge, the infection was maintained in DMSO or 1, 5 or 10 μM nocodazole supplemented 2% FBS DMEM medium for 6,5 h followed by cell lysis. Luminescence was determined as described before. For VSV-G.HIV-1-GFP challenge, the infection was maintained in DMSO or 1, 5 or 10 μM nocodazole supplemented 10% FBS DMEM medium with 8 µg mL^-1^ polybrene infection reagent. Seven hours after transduction, cells were washed with PBS and the medium was refreshed with 10% FBS DMEM medium. The percentage of infected GFP-positive cells was determined 48 h post-transduction by flow cytometry as described before.

For MX2-EGFP/tubulin confocal microscopy, 15,000 HeLa cells were seed into 8-well imaging µ-Slide (#80826, Ibidi GmbH) and transfected with MX2(1-715)-EGFP expression vector. Twenty-four hours after transfection, medium was refreshed with DMSO or 1, 5 or 10 μM of nocodazole supplemented culture medium. Three hours after nocodazole treatment cells were fixed using 2% paraformaldehyde (PFA) in PBS for 10 minutes at RT and permeabilized with 0.2% Triton X-100 in PBS for 10 min at RT. Detection of MTs was achieved by staining the treated cells with mouse monoclonal anti-α-tubulin antibody.

#### NanoBIT complementation assay

For NanoBIT experiments 12,000 HeLa cells were seeded in 96-well black plates and each well was transfected with 100 ng of total DNA using 1:3 ratio of FUGENE HD (#E2311, Promega). Different amounts of plasmid DNA were used to achieve comparable expression of LgBiT/FLAG-tagged and SmBiT/V5-tagged proteins. LgBit and SmBiT complementation generated luminescent signal was measured in live cells using Nano-Glo® Live Cell Assay System (#N2011, Promega) according to the manufacturer’s instructions. In brief, 24 h after transfection cells were washed with PBS and the medium was replaced with 100 μL of FBS-free DMEM medium. Luminescence was determined by adding 25 μL of Nano-Glo® Live Cell Reagent to each well and after a 10 min incubation period, measurements were taken for 15 min, 1 measurement per min, on a plate reader (EnSight Multimode Plate Reader or EnVision XCite 2105 Multimode Plate Reader, PerkinElmer). The final graph and heat map resulted by calculating the average of all 15 measurements for each biological replicate. For proteins expression analysis, after measurements, cells from a single well were washed with ice-cold PBS and collected in 2X Laëmmli buffer.

#### Confocal Microscopy

15,000 HeLa based were seeded into 8-well imaging µ-Slide (#80826, Ibidi GmbH). For transient expression experiments each well was transfected with 100 ng of DNA using 1:3 ratio of FUGENE HD (#E2311, Promega). Twenty-four hours after transfection or doxycycline stimulation, cells were fixed using, prewarmed to 37°C, 2% paraformaldehyde (PFA) in PBS for 10 min at RT. Fixed cells were washed three times for 5 min in PBS and then permeabilized with 0.2% Triton X-100 in PBS for 10 min at RT. Cells were washed with PBS followed by 1 h incubation in blocking buffer [PBS complemented with 0.5% BSA (IgG-Free, Protease-Free, #001-000-162, Jackson ImmunoResearch), 0.02% Triton X-100 and 1:100 diluted donkey serum] at RT. Cells were incubated overnight at 4°C with primary antibodies diluted in blocking buffer. The next day, cells were washed three times for 5 min in PBS and then incubated for 2 h with secondary antibodies diluted in blocking buffer. After the secondary antibody was washed three times for 5 min, nuclei were stained with DAPI (4’,6-Diamidino-2-Phenylindole, Dihydrochloride) (#D1306, Thermo Fisher Scientific) (dilution 1:1,000 in PBS) for 15 minutes at RT (unless indicating otherwise) and, after washing, cells were maintained in PBS or in polyvinyl alcohol mounting medium with DABCO (#10981, Supelco). Images of the CLICK experiment for the analysis of intracellular EdC/A labelled HSV-1 genomes were acquired on an LSM980 Airyscan (Carl Zeiss, Jena, Germany) with plan-apochromat 40x/1.4 oil immersion objectives, and 405-, 488-, 561-, and 639-nm lasers using the operating software ZEN Blue. Representative Z-Stacks were collected using the Airyscan 2 detector in the SuperResolution (SR) mode. All the other imaging data in this study, were collected on an LSM880 Airyscan (Carl Zeiss, Jena, Germany) with either a Plan-Apochromat 63x/1.4 Oil, DIC M27 or a Plan-Apochromat 40x/1.3 Oil DIC UV-IR M27. The operating software is ZEN Black 2.3 SP1. The lasers used to excite the different fluorophores were a 405, 561, and 633 nm diode laser, and the 488 nm line of an Argon laser. The corresponding filter sets used were BP 420-480, LP570, LP645, and BP 495-550 in conjunction with the Airyscan detector in the SuperResolution (SR) mode of the FastAiryScan or in full Airyscan. A pixel reassignment and 2D wiener deconvolution step were carried out post-acquisition in ZEN Black. Image processing was performed using the ZEN Blue 3.5 software. For quantification purposes: Z-stacks covering the whole cell volume were recorded with the optimal z-interval according to the Nyquist criterium being 0.16 μm. A pixel reassignment and 3D wiener deconvolution step were carried out post-acquisition in ZEN Black. Imaging of HIV IN-EGFP particles in MX2-mScarlet transiently expressing cells and pNL4-3.HIV-1 virus capsid intracellular localization in IFN-stimulated U87 CD4+CXCR4+ cells were done at the KU Leuven cell and tissue imaging core using LSM880 Airyscan (Carl Zeiss, Jena, Germany) in SuperResolution (SR) mode of the FastAiryScan mode with a Plan-Apochromat 63x/1.4 Oil, DIC M27. A field of view was selected with full cell volume and IN-EGFP/NL4-3.HIV-1 viral particles. Z-stacks were taken to visualize the bulk of cytoplasm and as many viral particles as possible. For imaging of HIV IN-EGFP particles in MX2-mScarlet transiently expressing cells IN-EGFP, MX2-mScarlet and NucSpot 650/665 were excited using laser wavelengths of 488 nm, 561 and 633 nm, respectively. For imaging of pNL4-3.HIV-1 virus capsid intracellular localization in IFN-stimulated U87 CD4+CXCR4+ cells MX2, NL4-3.HIV-1 capsids [AG3.0] and NucSpot 650/665 were excited using laser wavelengths of 488 nm, 561 and 633 nm, respectively. The corresponding emission filter sets used were BP 495-550, 495-620, and LP 645.

#### Live cell imaging

Cells were seeded and transfected into 8-well imaging µ-Slide (#80826, Ibidi GmbH) and maintained during the experiments in culture medium at at 37°C and 5% CO_2_. Twenty-four hours after transfection live cell imaging was carried out on the LSM880 Airyscan (Carl Zeiss, Jena, Germany) with a Plan-Apochromat 63x/1.4 Oil, DIC M27. For this purpose, we used the Airyscan detector in the SR mode of the FastAiryScan. The set-up is equipped with an incubation chamber allowing CO_2_ and temperature control which was set at 37°C. Cells were imaged as single z-slice. The focus over time was maintained making use of the definite focus 2. The cells were monitored over a period of 37 min and 45 sec (150 frames) with a time interval of 15 sec. Accelerated videos were generated in ZEN blue 3.5 using a frame rate of 8 frames/sec.

#### FRAP

Cells were seeded and transfected into 8-well imaging µ-Slide and maintained during the experiments in culture medium at at 37°C and 5% CO_2_. Twenty-four hours after transfection, FRAP analysis was carried out on the LSM880 Airyscan (Carl Zeiss, Jena, Germany) in the regular confocal mode with a Plan-Apochromat 63x/1.4 Oil, DIC M27. The set-up is equipped with an incubation chamber allowing CO_2_ and temperature control which was set at 37°C. For each condition 7-10 condensates were measured. In each measurement 3 regions of interest (ROI) were analysed: one that was bleached (R1), one non-bleached in the background (R3), and one non-bleached in the neighbourhood of the bleached ROI (R2) to correct for the bleaching effect during imaging. The bleaching step was carried out using the 488 nm line of the Argon laser (for EGFP) with 2 iterations with 100% laser power, while the 561 nm laser was used for imaging mScarlet-tagged proteins. In the imaging sequence 10 images were taken prior to bleaching and 55 after bleaching resulting in a sequence of 100-120 sec in total (indicated in figures). From these data individual frames/images from representative FRAP experiments for each condition were selected to visualise the pre-bleached and post-bleached condensates. The recovery curves were obtained by subtracting the background signal (R3) from the data measured in the bleached region of interest (ROI) (R1). The resulting data were normalised relevant to the bleach point set as 0 and then normalized making use of the non-bleached neighbouring ROI (R2). These normalized data were fitted in a curve with nonlinear regression using one phase association model made in GraphPad Prism. From these fitted data we could extract the half-time (t1/2) and the plateau recovery for each EGFP-tagged protein component of the condensates.

#### Line profiles in ZEN blue 3.5

To get information on the intensity distribution of (1) MX2-V5 with NUPL1-EGFP and (2) MX2 (1-715) and mutants EGFP-tagged with IPO4-mScarlet, a line was drawn (in the profile menu) through the structures of interest. The resulting graph shows the intensity distribution of the selected channels versus the distance of the line.

#### Quantification in Arivis vision 4D version 4.0.0 (Carl Zeiss)

Analysis is performed based on high resolution z-stacks recorded with a system optimized z-interval using the SR mode of the FastAiryScan of the LSM880 (Carl Zeiss, Jena, Germany). Quantification of the MX2 mutants V5-tagged condensates: For this analysis 54 randomly selected cells of each MX2 mutant were measured. MX2 condensates were identified using the blob finder operation and an object feature filter which excludes the objects smaller than 0.1 µm^3^ and with a roundness factor smaller then 0.7. The number of MX2 condensates per cell were determined for all the described mutants. Quantification of the MX2 condensates in different siRNA treatments: For this analysis 96 randomly selected doxycycline-induced HeLa MX2(1-715)-EGFP cells of each siRNA variant were measured. MX2 condensates were identified using the blob finder operation. To discriminate between the total and the larger condensates, an object feature filter was applied excluding all condensates smaller than 0.1 µm^3^ for the total population and smaller than 2 µm^3^ for the larger condensates. The number of total MX2 condensates (including large) and % of large condensates were determined per cell for all the described siRNA variants. Quantification of the number of HSV-1 capsids in the total cell volume and in the nucleus for the different MX2 mutants: For this analysis 44 randomly selected infected cells of each MX2 mutant were measured. HSV capsids were identified using the blob finder operation. The nuclei were identified using an intensity threshold segmenter operation on the DAPI signal. Making use of the compartment operation we identified the capsids within the nuclear signal. The total number of capsids and the % of nuclear capsids were determined per cell for each of the MX2 mutants. Quantification of the number of IN-EGFP PICs WT CA and IN-EGFP PICs P90A CA in the total cell volume, in the cytoplasm, colocalizing or coinciding completely with MX2(1-715) condensates per cell: For this analysis 42 randomly selected doxycycline-induced HeLa MX2(1-715)-V5 cells infected with each virus were measured. The nuclei were identified using cell diameter threshold segmenter Cellpose-based model: CP. IN-EGFP PICs were identified by using machine learning operation based on voxel training to exclude background signal and EGFP clusters at the outside of the cells followed by filtering PICs signal smaller than 0,01 µm^3^. MX2 condensates were identified using the blob finder operation using probability threshold (15%) and filtered by diameter. Cytoplasmic PICs were calculated by subtraction of the total number of intracellular PICs by the PICs inside the nuclei. Cytoplasmic PICs intersecting and inside MX2(1-715) condensates were identified and used for the calculation of the % of cytoplasmic PICs colocalizing with MX2(1-715) condensates. Cytoplasmic PICs inside MX2(1-715) condensates were identified and used for the calculation of the % of cytoplasmic PICs coinciding completely within MX2(1-715) condensates.

Images of the CLICK experiment for the analysis of intracellular EdC/A labelled HSV-1 genomes were acquired on an LSM980 Airyscan (Carl Zeiss, Jena, Germany) as single z-slices 2D images. The nuclei were identified using cell diameter threshold segmenter Cellpose-based model: Cyto. HSV capsids were identified by using machine learning operation based on voxel training to exclude background signals from the CLICK reaction in the nucleoli and the cytoplasm followed by filtering HSV-1 CLICK based on VoxelCount(Volume) smaller 10. Making use of the compartment operation we identified the HSV-1 genomes within the nuclear signal. The total number of capsids and the % of nuclear HSV-1 genomes were determined per cell for each of the MX2 mutants.

### Quantification and statistical analysis

Data are represented as mean ± s.d. of experimental replicates, with the exact statistical tests, significant p values and n values indicated in the figure legends. BioID experiments were performed with ≥3 biological replicates per condition. Differences between experimental groups were evaluated, either by paired or unpaired two-tailed t-tests using GraphPad Prism. Experimental data not subjected to statistical analysis (e.g. Western blots, NanoBit assay, confocal imaging, live cell imaging, virus capsids intracellular microscopy) have been reproduced in at least three independent experiments.

## Supplementary tables

**Supplementary Table 1: Proximity labeling proteomics identifies the MX2 proximal interactome.**

Sheet ’MX2 int’ provides an overview of the generated proteomics data. Proteins colored in light green highlight significant proteins shared between all conditions at 1% FDR. Proteins colored in yellow highlight significant proteins shared between all conditions at 5% FDR. Sheets ’DEA -IFN’, ’DEA +IFN’, and ’DEA HSV-1’ provide output of the differential enrichment analysis (FDR 1% and 5% with s0-value of 1) of MUTT2A vs T2A for -IFN, +IFN and HSV-1 infection conditions, respectively. General proteomics statistics for each protein, and log2(LFQ) values for each replicate are provided.

**Supplementary Table 2: Network analysis for common MX2 interactors.**

Sheet ’Venn diagramm’ provides the observed overlap between all interactors at 1% FDR for all experimental setups. Color codes represent overlaps with specific experiments (see legend).

Sheet ’SAFE analysis’ provides the output generated by SAFE analysis for the common interactors using BioGRID to generate PPI networks. Networks are annotated using GO: CC. Different networks discussed in the main text are highlighted.

**Supplementary Table 3: Overlapping RNP-associated proteins with Youn et al.**

RNP-associated proteins identified in Youn et al and this study are highlighted by color codes.

**Supplementary Table 4: Information for (ribo)nucleic acids used in this study**

Sheet ’Plasmid GOI info’ provides sequence information for the Gene of Interests (GOIs) coding DNA sequences (CDSs) that were introduced as individual modules, part of the Golden Gate assembly cloning system. For each GOI, gene symbol, official full name, gene ID, NCBI gene/protein accession number and UniProtKB/Swiss-Prot ID are indicated.

Sheet ’ON-TARGETplus SMARTpool siRNAs’ provides information for siRNAs used in this study. For each ON-TARGETplus SMARTpool siRNA target, the gene symbol, gene ID, gene accession, GI number, Pool catalog number, duplex catalog number and sequence are provided.

Sheet ’RT pPCR primers’ provides information for RT qPCR primers used in this study. For each RT qPCR primer pair, the target gene’s symbol, and sequence are provided.

Sheet ’ID&T ultramer oligo’ provides information for the sequence of the Utramer used for the generation of HeLa cells with endogenously tagged (dual V5 and HA tag) MX2 alleles.

