## Supplementary figures for "MX2 restricts HIV-1 and herpes simplex virus type 1 by forming cytoplasmic biomolecular condensates that mimic nuclear pore complexes"

#### Supplementary Figures legends

**Supplementary figure 1: MX2 ''T2A spilt-link design'' BioID screening set up.**

**(A)** MX2-T2A and MUTT2A-TurboID constructs. Top: Schematic presentation of the CDS of the untagged MX2 and the MX2-T2A or MUTT2A-TurboID that were cloned in the expression constructs. Bottom: HEK293T cells expressing either the untagged MX2 or one of the MX2-T2A or MUTT2A-TurboID were lysed and proteins were analysed by immunoblot using anti-MX2 and anti-V5 antibodies.

**(B)** Interaction of MX2-MUTT2A-TurboID with FLAG-MX2. Transfected HEK293T cells expressing N-terminally FLAG-tagged MX2 together with MX2-MUTT2A-TurboID were lysed and proteins were immunoprecipitated (IP) in the presence or absence of anti-FLAG antibody. The IP and input fractions were analysed by immunoblot using anti-MX2, anti-V5, and anti-tubulin (input fraction only) antibodies.

**(C)** HeLa cells either mock-transfected or transfected with expression vectors encoding MX1, MX2, MX2-T2A, or MUTT2A-TurboID were mock-infected or infected with HSV-1 strain C12 GFP reporter virus (MOI: 0.05). Viral growth was determined by real-time quantification of the GFP signal every 24 h until 72 h.p.i. Top: Data points indicate the green GFP fluorescent intensity normalized by subtracting the mean intensity of the mock-infected samples (mean ± s.d., 4 biological replicates). Middle: Relative area under the curve (AUC) from 0 h to 72 h.p.i. was calculated for each condition. AUC are relative to mock-transfected samples which are each set to 1 (mean ± s.d., n = 3 independent experiments, two-tailed unpaired *t*-test to mock-transfected condition; *p* values indicated with $*p < 0.05$, $**p < 0.01$, $***p < 0.001$, and $****p < 0.0001$). a.u., arbitrary units. Bottom: Transfected or mock-transfected, infected HeLa cells were lysed 48 h.p.i and cellular and HSV-1 proteins in the lysates were analysed by immunoblot using anti-MX2, anti-V5, anti-tubulin and anti-NC-1(VP5) antibodies.

**(D)** Top: HeLa cells either mock-transfected or transfected with expression vectors encoding MX1, MX2, MX2-T2A-TurboID, or MX2-MUTT2A-TurboID (300 ng plasmid) or expressing MX2 together with MX2-T2A-TurboID or MX2-MUTT2A-TurboID (150 ng plasmid each). Twenty-four hours after transfection, cells were challenged with HIV-1-GFP (MOI:0.2). Top: Transduction efficiency was assessed 48 h post-challenge by flow-cytometry. Middle: the percentage of $GFP^+$ cells is shown relative to mock-transfected condition set as 100 (mean ± s.d., n = 3 independent experiments, two-tailed unpaired *t*-test to mock-transfected condition; *p* values indicated with $*p < 0.05$, $**p < 0.01$, $***p < 0.001$). Bottom: Transfected or mock-transfected HeLa cells were lysed 24h post-transfection and proteins were analysed by immunoblot using anti-MX2, anti-V5 and anti-tubulin antibodies.

**(E)** Left: HeLa cells expressing MX2-T2A-TurboID or MX2-MUTT2A-TurboID. Intracellular localization of the V5-tagged TurboID protein and MX2 was assessed 24 h post-transfection by confocal microscopy using anti-V5-tag and anti-MX2 specific antibodies. Nuclei were stained with DAPI. Scale bar: 20 µm. Right: Intensity profile graphs of V5 (TurboID), MX2, and DAPI.

**(F)** Left: HeLa cells expressing MX2-MUTT2A-TurboID together with C-terminally HA-tagged MX2. Intracellular localization of MX2-MUT2A-TurboID and MX2-HA was assessed 24 h post-transfection by confocal microscopy using mAb414 and anti-MX2 specific antibodies. Nuclei were stained with DAPI. Scale bar: 20 µm. Right: Intensity profile graphs of mAb414, MX2, and DAPI.

**(G)** Left: HeLa cells expressing MX2-MUTT2A-TurboID. Intracellular localization of MX2-MUT2A-TurboID(V5) and MX2-HA was assessed 24 h post-transfection by confocal microscopy using anti-V5-tag and anti-HA-tag antibodies. Nuclei were stained with DAPI. Scale bar: 20 µm. Right: Intensity profile graphs of V5 (TurboID), HA (MX2-HA), and DAPI.

**(H)** Doxycycline-inducible U87-MG MX2-T2A-TurboID and U87-MG MX2-MUTT2A-TurboID cells were mock-stimulated or stimulated with 5, 15, 50, 250 or 500 ng mL$^{-1}$ doxycycline. Twenty-four hours after stimulation, biotin (50 µM) was added to the cells. Cells were lysed 2 h after biotin addition and proteins were analysed by immunoblot using anti-MX2, anti-V5 and anti-tubulin antibodies. The biotinylation pattern in each condition was assessed on the same immunoblot using Alexa Fluor™ 680 conjugated streptavidin. The doxycycline concentrations used for the MS experiments are indicated with black arrows.

**(I)** Schematic representation of the MX2 ''T2A split-link design'' proximity biotinylation (BioID) screening. MX2 BioID was performed in three different conditions (-IFN, +IFN, and HSV-1). U87-MG MX2-T2A-TurboID and U87-MG MX2-MUTT2A-TurboID cell lines were stimulated with 15 ng mL$^{-1}$ doxycycline for 24 h, in the absence or presence of 1000 IU mL$^{-1}$ human IFNα2α, followed by the addition of 50 µM biotin for 2 h to biotinylate proteins in the proximity of free or MX2 linked TurboID, respectively. For the HSV-1 infected samples, biotin was added simultaneously with HSV-1 strain F infection (MOI:20). Biotinylation was allowed for 2 h for all the samples. After cell lysis, biotinylated proteins were enriched on streptavidin-conjugated beads and digested by trypsin on the beads. The resulting peptides were identified and quantified by LC-MS/MS analysis. Figure created with BioRender software.

**(J)** Profile plot of MX2 (cyan), V5-TurboID (orange) and three housekeeping proteins, VIM, GAPDH and EEF1A1 (different shades of purple), showing $\log_2$(LFQ Intensity) values for each individual MX2-T2A-TurboID and MX2-MUTT2A-TurboID MS sample across all three MX2 BioID conditions.

For all fluorescence imaging, DAPI is shown in cyan, single channels are shown in inverted grey and the merged images with colours indicated in each panel.

**Supplementary figure 2: MX2 associates with mitochondrial and coated vesicles proteins, besides nuclear transport proteins and granule-associated ribonucleoproteins.**

**(A)** Bar charts showing the top 10 Gene Ontology (GO) terms for biological processes (top) and cellular components (bottom), ranked by -$\log_{10}$(adjusted p-value) of the commonly enriched MX2 interactors.

**(B)** Top: Nodes and edges of the SAFE enriched ''mitochondrial containing-protein complex'' domain protein network. Bottom: Dotplot enrichment analysis of mitochondrial protein network. Endogenously biotinylated mitochondrial proteins that are known BioID contaminants are indicated with a red exclamation mark.

**(C)** Top: Nodes and edges of the SAFE enriched ''coated vesicle'' domain protein network. Bottom: Dotplot enrichment analysis of coated vesicle domain proteins ranked according to their average enrichment ($\log_2$(FC)) in all three conditions.

**(D)** Nodes and edges of the SAFE enriched ''granule ribonucleoprotein'' domain protein network.

**Supplementary figure 3: Knockdown of SAMD4A, TNPO1, and NUP35 affects MX2-mediated antiviral activity against HIV-1 and HSV-1.**

**(A)** RT-qPCR data showing relative to Non-Targeting expression mRNA levels of each targeted gene in HeLa(MX2) cells 24 h after transfection with the indicated SMARTpool siRNAs.

**(B-C)** Doxycycline-inducible HeLa(MX2) cells were transfected with siRNA and left untreated or treated with 100 ng $mL^{-1}$ doxycycline. **(B)** Left: 48 h after transfection, cells were infected with HSV-1 firefly reporter virus (MOI: 1). Luminescence is shown relative to the Non-Targeting (-doxy) condition. Immunoblots showing the presence of MX2 and tubulin in cell lysates. Right: Fold MX2-mediated HSV-1 inhibition calculated by dividing luminescence of untreated by doxycycline-induced samples in each condition. **(C)** Left: 48 h after transfection, cells were inoculated with VSV-G.HIV-1-GFP (MOI: 0.4). Right: Fold MX2-mediated HIV-1 inhibition calculated by dividing % of $GFP^+$ cells in untreated by doxycycline-induced samples in each condition. **(B, C)** (mean ± s.d., n = 4 independent experiments, two-tailed unpaired *t*-test to Non-Targeting siRNA; *p* values indicated with *$p < 0.05$, **$p < 0.01$, ***$p < 0.001$).

(D) For **(A and B)** cell lysates to document protein knockdown were collected 48 hours post-transfection. Immunoblot analysis was used to analyse protein expression using antibodies specific for the indicated proteins.

**(E)** HeLa cells were transfected with siRNAs targeting the indicated genes together with either Non-Targeting or MX2 siRNAs and left untreated or were treated with 1,000 IU $ml^{-1}$ of rhIFN-a2a. Twenty-four hours after transfection, cells were inoculated with VSV-G.HIV-1-GFP (MOI: 0.2). Ratio of % $GFP^+$ cells between untreated and the doxycycline-induced samples in each condition is shown (mean ± s.d., n = 5 independent experiments, two-tailed paired *t*-test to Non-Targeting siRNA; *p* values indicated with $*p < 0.05$, $**p < 0.01$, $***p < 0.001$). Immunoblots were used to reveal MX2 and tubulin expression in the cell lysates.

**(F)** Protein knockdown documented in cell lysates collected 48 hours post-transfection. Immunoblot analysis was used to analyse protein expression in these lysates using antibodies specific for the indicated proteins.

**(G)** Flow cytometry gating strategy following RNAi and HIV-1-GFP transduction for (top) HeLa and HeLa(MX2) cells and (bottom) U87-MG and U87-MG(MX2) cells. Cells were gated using the forward scatter (FSC) and side scatter (SSC). Single cells were selected using FSC-H and FSC-A. The $GFP^+$ population was gated using uninfected cells as negative control.

**Supplementary figure 4: Anti-HIV-1 and -HSV-1 activity of fluorescently tagged MX2 proteins.**

**(A)** HeLa cells either mock-transfected or transfected with expression vectors coding for MX2, or EGFP or mScarlet C-terminally tagged MX2. Top: Twenty-four hours after transfection, cells were challenged with VSV-G.HIV-1-GFP (MOI: 0.5). Twenty-four hours later DNA was extracted and qPCR_was performed to detect 2-LTR circles for VSV-G.HIV-1-infected cells. The normalised number of 2-LTR circles is shown relative to mock-transfected condition set as 1. (Middle) Twenty-four hours later cells were infected with HSV-1 firefly reporter virus (MOI: 1). Luminescence is shown relative to the mock-transfected condition set as 1. (mean ± s.d., n = 3 independent experiments, two-tailed unpaired *t*-test to mock-transfected condition; *p* values indicated with $*p < 0.05$, $**p < 0.01$, $***p < 0.001$). Bottom: Transfected or mock-transfected HeLa cells were lysed 24h post-transfection and proteins were analysed by immunoblot using anti-MX2 and anti-tubulin antibodies.

**Supplementary figure 5: MX2 condensates interact with PBs and SGs and comprise RNA-binding or/and disordered domain proteins.**

**(A)** Confocal microscopy of HeLa cells expressing EGFP-tagged LSM14A in the presence or absence of MX2-V5.

**(B)** HeLa cells expressing MX2-mScarlet together with EGFP-tagged LSM14A were imaged under time-lapse conditions.

**(C)** Confocal microscopy of HeLa cells expressing EGFP-tagged G3BP1 in the presence or absence of MX2-V5. Twenty-four hours post-transfection, cells were mock-treated or treated with 500 µM of sodium arsenite ($NaAsO_2$) for 45mins. Intracellular localization of the EGFP- and V5-tagged proteins was assessed by confocal microscopy.

**(D)** Confocal microscopy of HeLa cells expressing EGFP-tagged SAMD4B, FUBP3, or CNOT4 in the presence or absence of MX2-V5.

**(E)** Confocal microscopy of HeLa cells expressing NUPL1-V5, MX2-mScarlet and EGFP-tagged SAMD4B, FUBP3, or CNOT4.

**(F)** HeLa cells expressing MX2-mScarlet together with EGFP-tagged SAMD4B or FUBP3 were imaged under time-lapse conditions.

**(G)** HeLa cells expressing EGFP-tagged SAMD4B or FUBP3, together with MX2-mScarlet. FRAP was performed for the complete individual condensates 24 h post-transfection. Upper panel: Normalized FRAP curves obtained for EGFP signal over time (n=7). Plateau and half-time are indicated on the right of each curve. Bottom panel: Representative images showing the biomolecular condensates during FRAP (before, during, and 34 and 84 sec post-bleaching). Bleached, non-bleached, and background areas are highlighted with solid, dashed or dotted, white circles, respectively. Scale bar: 10 µm.

(**A, C, D, E)** Nuclei were stained with DAPI. Scale bar: 20 µm. Right "zoom" panels show an enlarged view of the highlighted white boxed area in individual fluorescence channels and channels overlay (without DAPI).

(**B, F)** Bottom "zoom" panels show an enlarged view of the highlighted white boxed area. Moving and fusing MX2 condensates are indicated with white arrows. Real time of imaging is indicated at the bottom left side of each image. Scale bar: 10 µm.

For all fluorescence imaging, DAPI is shown in cyan, single channels are shown in inverted grey and the merged images with colours indicated in each panel.

**Supplementary figure 6: MX2 N-terminal disordered domain and dimerization are required for condensate formation and anti-HIV-1 and -HSV-1 antiviral activity.**

**(A, B)** Doxycycline-inducible HeLa MX2(1-715) and MX2 mutants, C-terminally V5- tagged, were treated with different concentrations of doxycycline to obtain comparable expression levels.

**(A)** Forty-eight hours after stimulation, the cells were challenged with HIV-1-GFP (MOI:0.2). Transduction efficiency was assessed 48 h post-challenge by flow-cytometry. Percentage of $GFP^+$ cells of each HeLa MX2 cell line are shown relative to HeLa MX2(1-715) samples which are each set to 100 (mean ± s.d., n = 5 independent experiments). Flow cytometry gating strategy: cells were gated using the forward scatter (FSC) and side scatter (SSC). Single cells were selected using FSC-H and FSC-A. The $GFP^+$ population was gated using uninfected cells as negative control. Representative plots showing MX2(1-715) and MX2(M574D) are shown.

**(B)** Twenty-four hours after stimulation, cells were challenged with HSV-1 strain C12 (MOI:0.05). Viral growth was determined by real-time quantification of GFP signal every 24 h until 72 h.p.i. Upper panel:

Data points indicate the green GFP fluorescent intensity normalized to the mean intensity of the mock-infected samples (mean ± s.d., n = 4 biological replicates). a.u., arbitrary units. Lower panel: Relative area under the curve (AUC) from 0 h to 72 h.p.i. was calculated for each HeLa MX2 cell line. AUC are relative to HeLa MX2(1-715) samples which are each set to 100 (mean ± s.d., n = 5 independent experiments).

**(C)** HeLa cells expressing C-terminally EGFP-tagged MX2(1-715) and mutants. Intracellular localization of the EGFP-tagged proteins was assessed 24 h post-transfection by confocal imaging. Localization of EGFP-tagged MX2 proteins visualized in white. Scale bar: 20 µm. Bottom "zoom" panels show an enlarged view of the highlighted area.

**(D)** HeLa cells expressing MX2 N-terminal domain chimeric MX1, with C-terminal HA-tag, constructs. Intracellular localization of $_{NTDMX2}$MX1 chimeric proteins and endogenous FG-Nups was assessed 24 h post-transfection by confocal microscopy.

**(E)** HeLa cells expressing MX2 N-terminal domain chimeric EGFP, constructs. Intracellular localization of $_{NTDMX2}$EGFP chimeric proteins and endogenous FG-Nups visualized with mAb414, was assessed 24 h post-transfection by confocal microscopy.

**(D, E)** Nuclei were stained with DAPI (cyan). Scale bar: 20 µm. Bottom "zoom" panels show an enlarged view of the white boxed area (without DAPI). Single channels are shown in inverted grey and the merged images with colours indicated in each panel.

**(F)** Immunoblot corresponding to the MX2 mutants in the NanoBiT experiments (see **Figure 5H**) using anti-FLAG, anti-V5, and anti-tubulin antibodies.

For all fluorescence imaging, DAPI is shown in cyan, single channels are shown in inverted grey and the merged images with colours indicated in each panel, unless indicated otherwise.

**Supplementary figure 7: MX2 condensates interact with incoming HIV-1 and HSV-1 capsids.**

**(A, B, C)** HeLa (rtTA/tTS) cells and doxycycline-inducible HeLa MX2(1-715), MX2(26-715), and MX2(T151A) C-terminally EGFP-tagged cells were treated with different concentrations of doxycycline to obtain comparable expression levels.

**(A)** Twenty-four hours after induction, cells were lysed and protein expression analysed by immunoblot using anti-EGFP and anti-tubulin antibodies.

**(B)** Intracellular localization of the EGFP-tagged MX2 proteins assessed 24 h post-induction by confocal imaging.

**(C)** Intracellular localization of EGFP-tagged MX2 proteins and endogenous alpha-tubulin were assessed 24 h post-induction by confocal microscopy. Nuclei were stained with DAPI. Scale bar: 20 µm.

**(D)** HeLa cells mock-transfected or expressing MX2(1-715)-EGFP. Twenty-four hours post-transfection, cells were mock-infected or challenged with HIV-1 based lentiviral vector (MOI: 25). Six hours after transduction, intracellular localization of the MX2 proteins and HIV-1 capsids was assessed by confocal microscopy using anti-p24 HIV-1 clone [*39/5.4A]* antibody. Right "zoom" panel shows an enlarged view of the highlighted white boxed area in channels overlay (with DAPI). Incoming HIV-1 capsids colocalizing with MX2 condensates are indicated with white arrow heads. Nuclei were stained with DAPI. Scale bar: 20 µm.

**(E)** U87 CD4+CXCR4+ cells were left untreated or treated with 1,000 IU $ml^{-1}$ of rhIFN-a2a. Twenty-four hours after IFN stimulation cells were infected with $8 \times 10^5$ pg (0,8 ug) of p24 replication-competent NL4-3.HIV-1. Six hours after infection, intracellular localization of MX2 and HIV-1 capsids was assessed by confocal microscopy with z-series image acquisition using anti-p24 HIV-1 clone *[AG3.0]* and anti-MX2 specific antibodies. Single z-slice view of the complete cell is shown. z1-4: z-slice 1-4 of the highlighted area. Incoming HIV-1 capsids colocalizing with MX2 condensates are indicated with white arrows. Nuclei were stained with NucSpot 650/665 (white). Scale bar: 20 µm.

**(F)** CLICK experiments for the analysis of intracellular EdC/A labelled HSV-1 genomes in doxycycline-inducible V5-tagged MX2(1-715) or mutant MX2 variants HeLa cells with or without doxycycline induction. Top: Intracellular localization of the MX2 proteins, viral capsids and viral genomes were visualized by confocal microscopy. Nuclei were stained with DAPI. Scale bar: 20 µm.

For all fluorescence imaging, DAPI is shown in cyan, single channels are shown in inverted grey and the merged images with colours indicated in each panel.

**Supplementary figure 8: Depolymerization of the microtubule network affects anti-HIV-1 restriction by MX2.**

**(A)** HeLa cells mock-transfected or expressing MX2(1-715)-GFP, intracellular localization of MX2-GFP and endogenous alpha-tubulin was assessed 24 h post-transfection by confocal microscopy in the presence or absence of anti-alpha-tubulin antibody.

**(B)** HeLa cells expressing MX2(1-715)-EGFP. Twenty-four hours post-transfection, cells were either mock-treated with DMSO or treated with 1, 5, or 10 µM of nocodazole. Intracellular localization of MX2-EGFP and tubulin were assessed 3 h after treatment by immunostaining using anti-alpha-tubulin antibody.

**(C)** Doxycycline-inducible HeLa MX2 cells were left untreated or were treated with 100 ng $mL^{-1}$ doxycycline. After 48 h, cells were either mock-treated with DMSO or treated with 1, 5, or 10 µM of nocodazole. Three hours after nocodazole treatment the cells were challenged with HIV-1-GFP in the presence of similar DMSO or nocodazole treatment for 7 h before medium was refreshed with normal growth medium. The transduction efficiency was assessed 48 h post-challenge by flow-cytometry. Flow cytometry gating strategy following nocodazole treatment. Cells were gated using the forward scatter (FSC) and side scatter (SSC). Single cells were selected using FSC-H and FSC-A. $GFP^+$ population was gated using uninfected cells as negative control. Representative plots showing $GFP^+$ cells for DSMO mock-treated samples with and without doxycycline stimulation is shown.

Figure S1

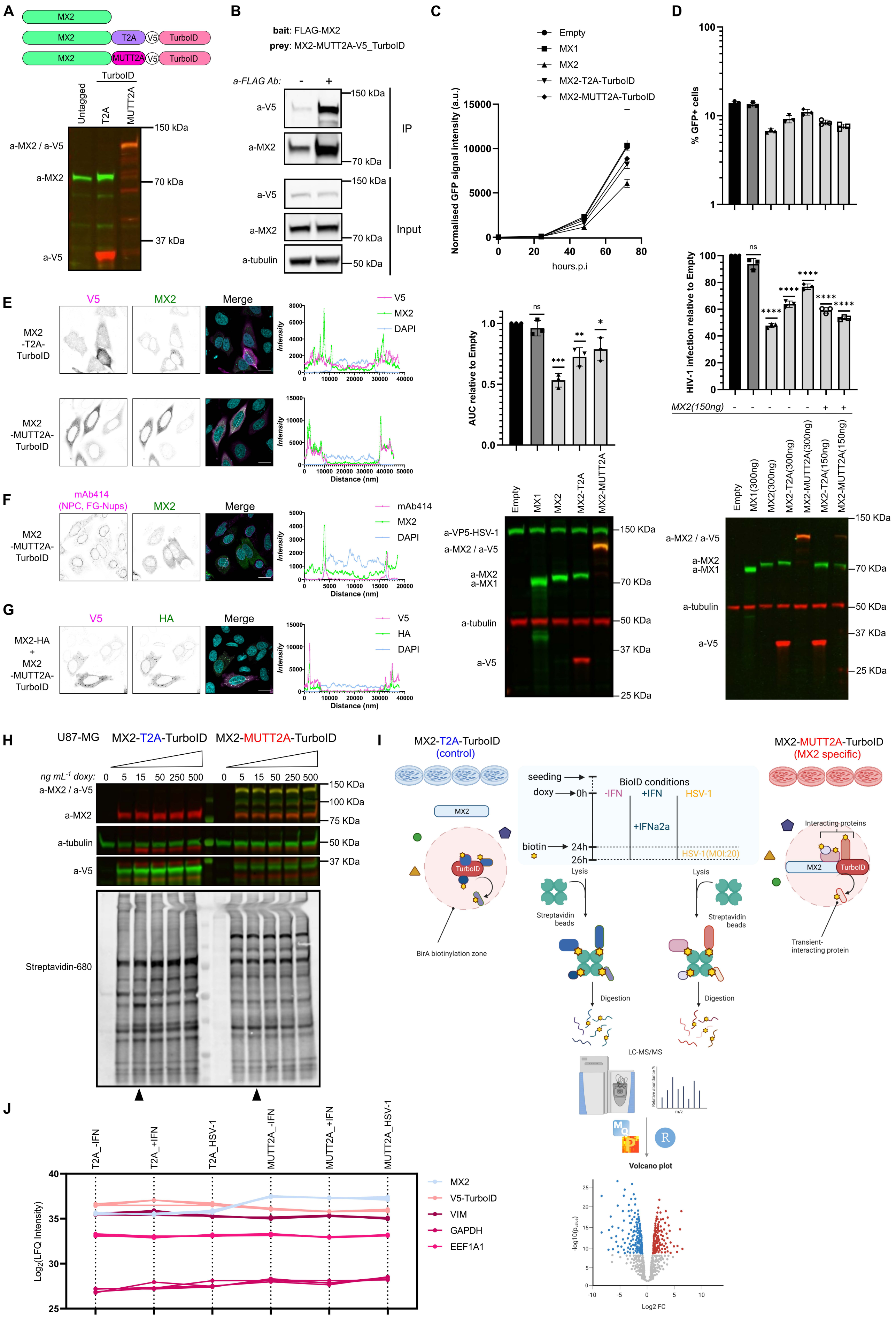

A
MX2
T2A
V5
TurboID
MUTT2A
Untagged
a-MX2 / a-V5
a-MX2
a-V5
150 kDa
70 kDa
37 kDa
B
bait: FLAG-MX2
prey: MX2-MUTT2A-V5_TurboID
a-FLAG Ab:
IP
Input
a-tubulin
50 kDa
C
Empty
MX1
MX2
MX2-T2A-TurboID
MX2-MUTT2A-TurboID
Normalised GFP signal intensity (a.u.)
hours.p.i
AUC relative to Empty
D
% GFP+ cells
HIV-1 infection relative to Empty
MX2(150ng)
MX1(300ng)
MX2(300ng)
MX2-T2A(300ng)
MX2-MUTT2A(300ng)
MX2-T2A(150ng)
MX2-MUTT2A(150ng)
a-VP5-HSV-1
a-MX2
a-MX1
E
V5
Merge
DAPI
MX2-T2A-TurboID
MX2-MUTT2A-TurboID
Intensity
Distance (nm)
F
mAb414
(NPC, FG-Nups)
G
HA
MX2-HA + MX2-MUTT2A-TurboID
H
U87-MG
MX2-T2A-TurboID
MX2-MUTT2A-TurboID
ng mL-1 doxy:
Streptavidin-680
100 KDa
75 KDa
I
(control)
(MX2 specific)
seeding
doxy
biotin
BioID conditions
-IFN
+IFN
HSV-1
+IFNa2a
HSV-1(MOI:20)
Lysis
Streptavidin beads
Digestion
LC-MS/MS
Volcano plot
BirA biotinylation zone
Interacting proteins
Transient-interacting protein
Log2 FC
J
T2A_-IFN
T2A_+IFN
T2A_HSV-1
MUTT2A_-IFN
MUTT2A_+IFN
MUTT2A_HSV-1
Log2(LFQ Intensity)
V5-TurboID
VIM
GAPDH
EEF1A1

Figure S2

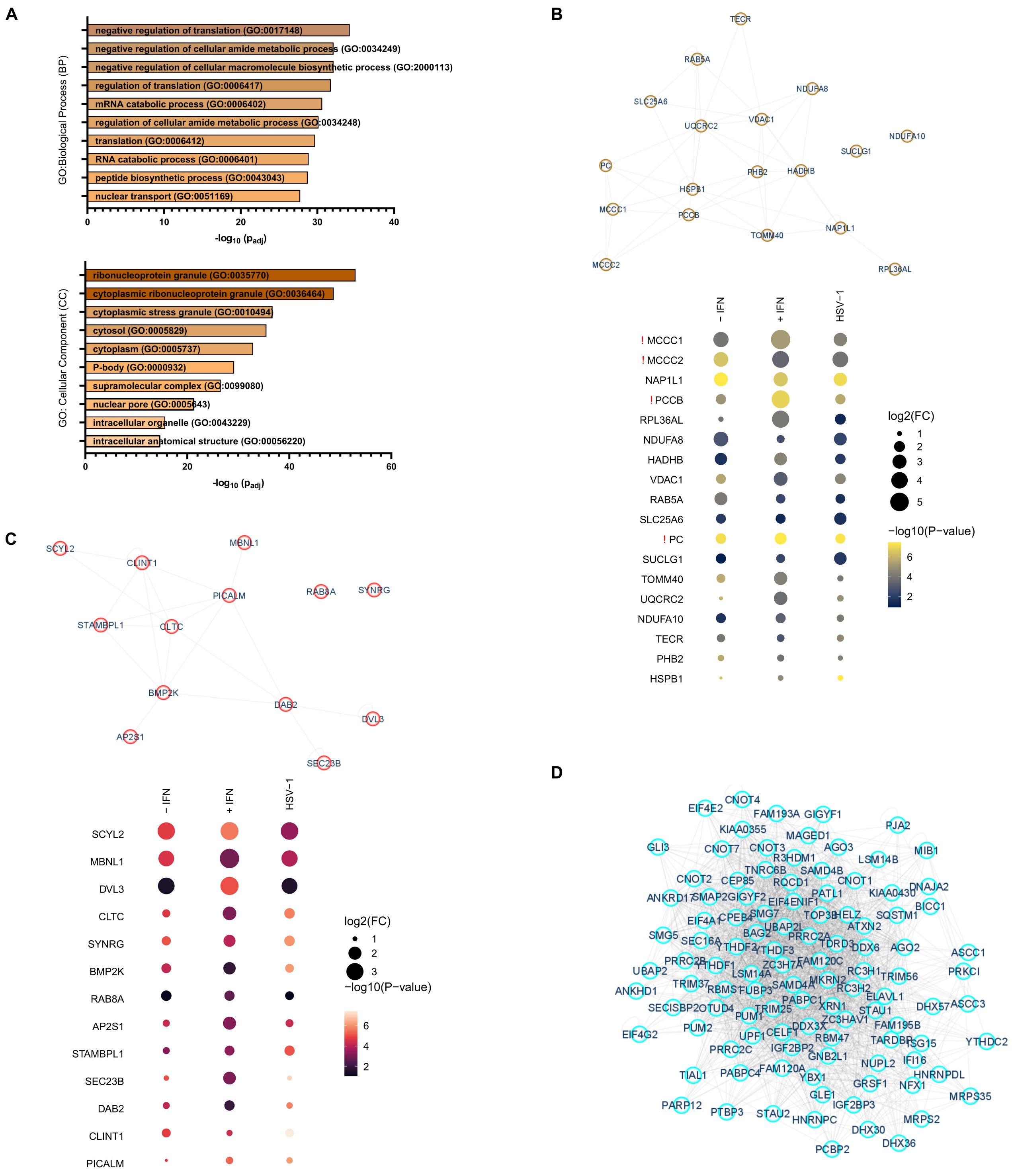
A
negative regulation of translation (GO:0017148)
negative regulation of cellular amide metabolic process (GO:0034249)
negative regulation of cellular macromolecule biosynthetic process (GO:2000113)
regulation of translation (GO:0006417)
mRNA catabolic process (GO:0006402)
regulation of cellular amide metabolic process (GO:0034248)
translation (GO:0006412)
RNA catabolic process (GO:0006401)
peptide biosynthetic process (GO:0043043)
nuclear transport (GO:0051169)
GO:Biological Process (BP)
0
10
20
30
40
-log10 (Padj)
ribonucleoprotein granule (GO:0035770)
cytoplasmic ribonucleoprotein granule (GO:0036464)
cytoplasmic stress granule (GO:0010494)
cytosol (GO:0005829)
cytoplasm (GO:0005737)
P-body (GO:0000932)
supramolecular complex (GO:0099080)
nuclear pore (GO:0005643)
intracellular organelle (GO:0043229)
intracellular anatomical structure (GO:00056220)
GO: Cellular Component (CC)
0
20
40
60
-log10 (Padj)
B
TECR
RAB5A
NDUFA8
SLC25A6
VDAC1
UQCRC2
NDUFA10
SUCLG1
PC
PHB2
HADHB
HSPB1
MCCC1
PCCB
TOMM40
NAP1L1
MCCC2
RPL36AL
− IFN
+ IFN
HSV−1
! MCCC1
! MCCC2
NAP1L1
! PCCB
RPL36AL
NDUFA8
HADHB
VDAC1
RAB5A
SLC25A6
! PC
SUCLG1
TOMM40
UQCRC2
NDUFA10
TECR
PHB2
HSPB1
log2(FC)
1
2
3
4
5
−log10(P−value)
6
4
2
C
SCYL2
MBNL1
CLINT1
PICALM
RAB8A
SYNRG
STAMBPL1
CLTC
BMP2K
DAB2
DVL3
AP2S1
SEC23B
− IFN
+ IFN
HSV−1
SCYL2
MBNL1
DVL3
CLTC
SYNRG
BMP2K
RAB8A
AP2S1
STAMBPL1
SEC23B
DAB2
CLINT1
PICALM
log2(FC)
1
2
3
−log10(P−value)
6
4
2
D
CNOT4
EIF4E2
FAM193A
GIGYF1
PJA2
KIAA0355
MAGED1
GLI3
CNOT7
CNOT3
AGO3
MIB1
R3HDM1
LSM14B
TNRC6B
SAMD4B
CNOT2
CEP85
RQCD1
CNOT1
DNAJA2
PATL1
KIAA0430
BICC1
EIF4A1
CPEB4
SMG7
TOP3B
HELZ
SQSTM1
ATXN2
SMG5
SEC16A
BAG2
UBAP2L
PRRC2A
YTHDF2
YTHDF3
TDRD3
DDX6
AGO2
ASCC1
PRRC2B
FAM120C
YTHDF1
ZC3H7A
UBAP2
LSM14A
RC3H1
TRIM56
PRKCI
TRIM37
SAMD4A
MKRN2
RBMS1
FUBP3
RC3H2
ANKHD1
PABPC1
ELAVL1
SECISBP2
OTUD4
TRIM25
XRN1
STAU1
DHX57
ASCC3
PUM1
ZC3HAV1
DDX3X
FAM195B
EIF4G2
PUM2
CELF1
UPF1
RBM47
TARDBP
ISG15
YTHDC2
PRRC2C
IGF2BP2
GNB2L1
NUPL2
IFI16
FAM120A
HNRNPDL
TIAL1
PABPC4
YBX1
GRSF1
NFX1
GLE1
MRPS35
PARP12
IGF2BP3
PTBP3
STAU2
HNRNPC
DHX30
MRPS2
PCBP2
DHX36

### Figure S3

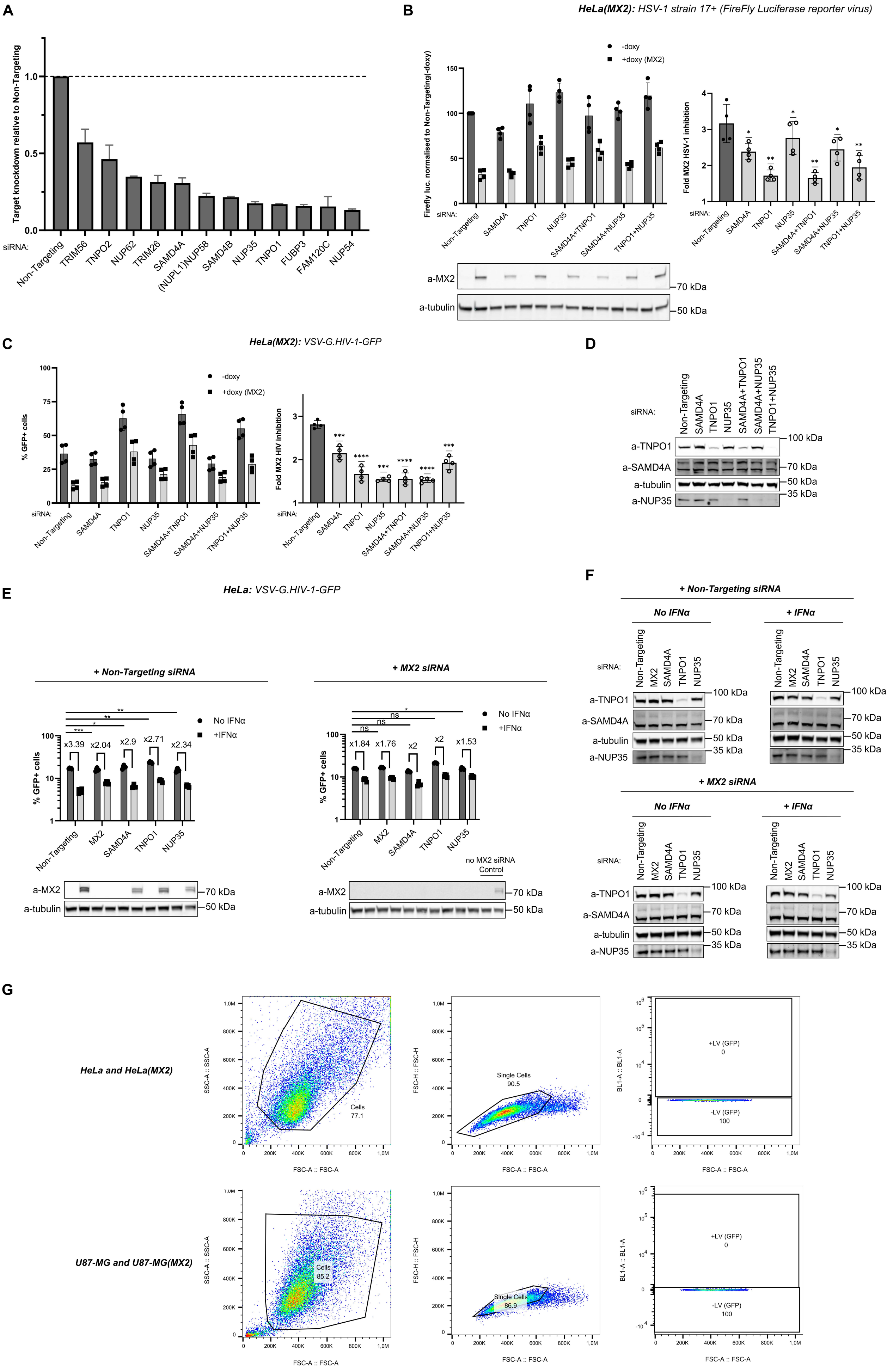
A
Target knockdown relative to Non-Targeting
siRNA:
Non-Targeting
TRIM56
TNPO2
NUP62
TRIM26
SAMD4A
(NUPL1)NUP58
SAMD4B
NUP35
TNPO1
FUBP3
FAM120C
NUP54
B
HeLa(MX2): HSV-1 strain 17+ (FireFly Luciferase reporter virus)
-doxy
+doxy (MX2)
Firefly luc. normalised to Non-Targeting(-doxy)
Fold MX2 HSV-1 inhibition
SAMD4A+TNPO1
SAMD4A+NUP35
TNPO1+NUP35
a-MX2
a-tubulin
70 kDa
50 kDa
C
HeLa(MX2): VSV-G.HIV-1-GFP
% GFP+ cells
Fold MX2 HIV inhibition
D
a-TNPO1
a-SAMD4A
a-tubulin
a-NUP35
100 kDa
35 kDa
E
HeLa: VSV-G.HIV-1-GFP
+ Non-Targeting siRNA
+ MX2 siRNA
No IFNα
+IFNα
x3.39
x2.04
x2.9
x2.71
x2.34
x1.84
x1.76
x2
x2
x1.53
ns
no MX2 siRNA Control
F
+ Non-Targeting siRNA
+ MX2 siRNA
No IFNα
+ IFNα
MX2
G
HeLa and HeLa(MX2)
U87-MG and U87-MG(MX2)
SSC-A :: SSC-A
FSC-A :: FSC-A
FSC-H :: FSC-H
BL1-A :: BL1-A
Cells
77.1
85.2
Single Cells
90.5
86.9
+LV (GFP)
0
-LV (GFP)
100

Figure S4

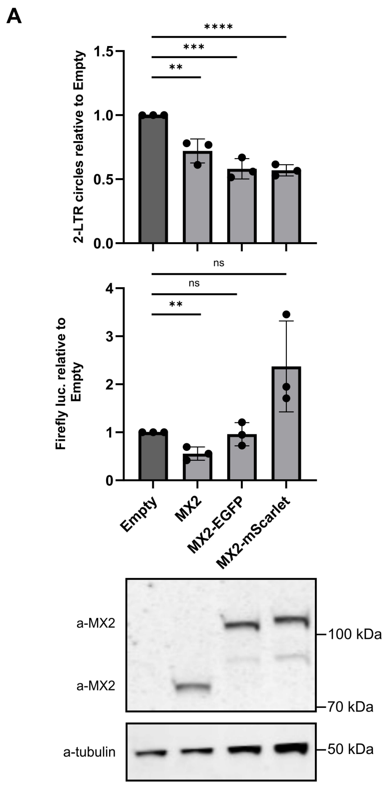
A
2-LTR circles relative to Empty
1.5
1.0
0.5
0.0
****
***
**
Firefly luc. relative to Empty
4
3
2
1
0
ns
ns
**
Empty
MX2
MX2-EGFP
MX2-mScarlet
a-MX2
a-MX2
a-tubulin
100 kDa
70 kDa
50 kDa

### Figure S5

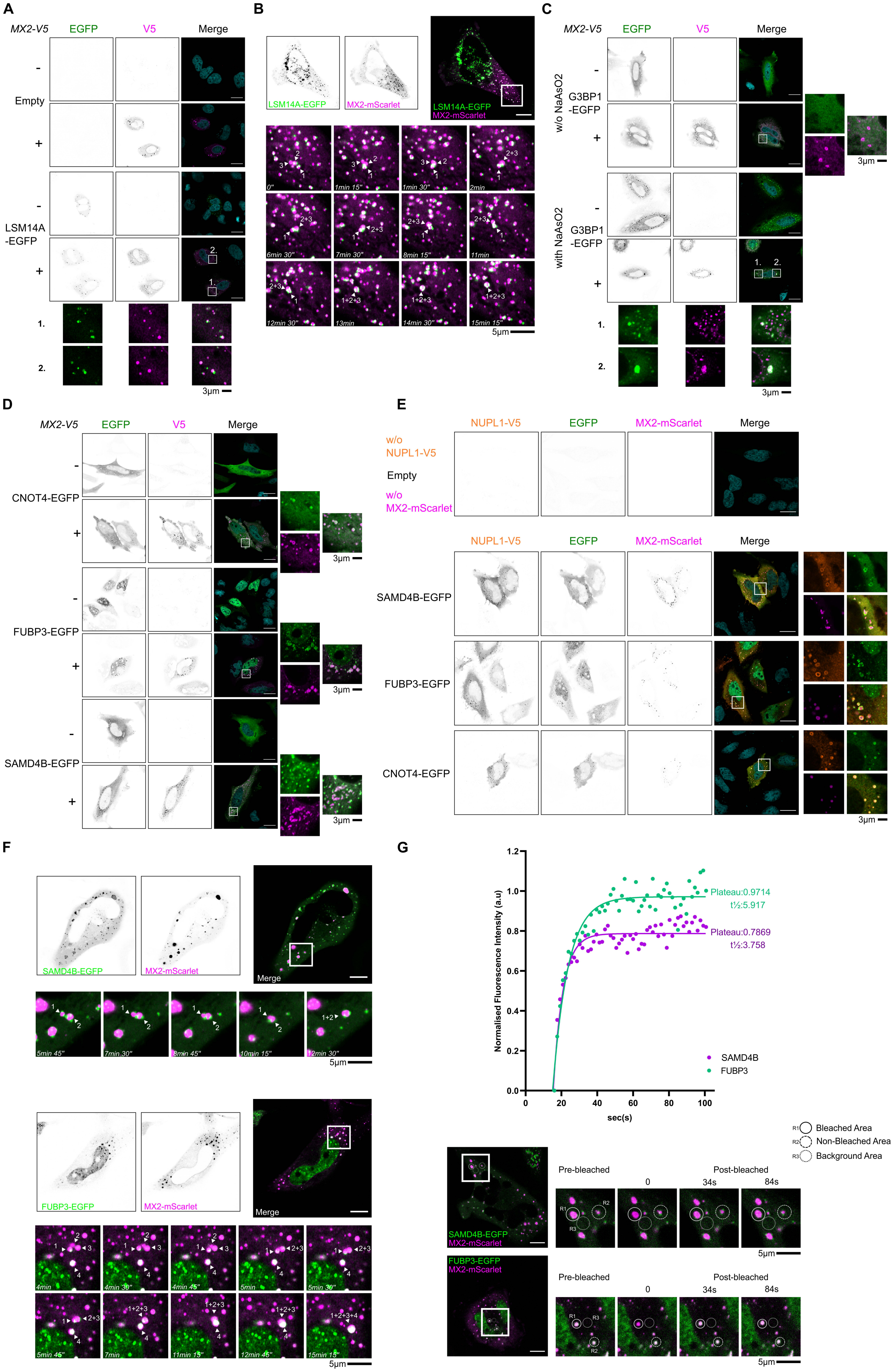

A
MX2-V5
EGFP
V5
Merge
Empty
LSM14A
-EGFP
3µm
B
LSM14A-EGFP
MX2-mScarlet
5µm
C
w/o NaAsO2
with NaAsO2
G3BP1
-EGFP
3µm
D
CNOT4-EGFP
FUBP3-EGFP
SAMD4B-EGFP
3µm
E
NUPL1-V5
MX2-mScarlet
w/o
NUPL1-V5
Empty
w/o
MX2-mScarlet
SAMD4B-EGFP
FUBP3-EGFP
CNOT4-EGFP
3µm
F
SAMD4B-EGFP
MX2-mScarlet
Merge
FUBP3-EGFP
5µm
G
Plateau:0.9714
t½:5.917
Plateau:0.7869
t½:3.758
Normalised Fluorescence Intensity (a.u)
sec(s)
SAMD4B
FUBP3
Bleached Area
Non-Bleached Area
Background Area
Pre-bleached
Post-bleached
0
34s
84s
5µm

### Figure S6

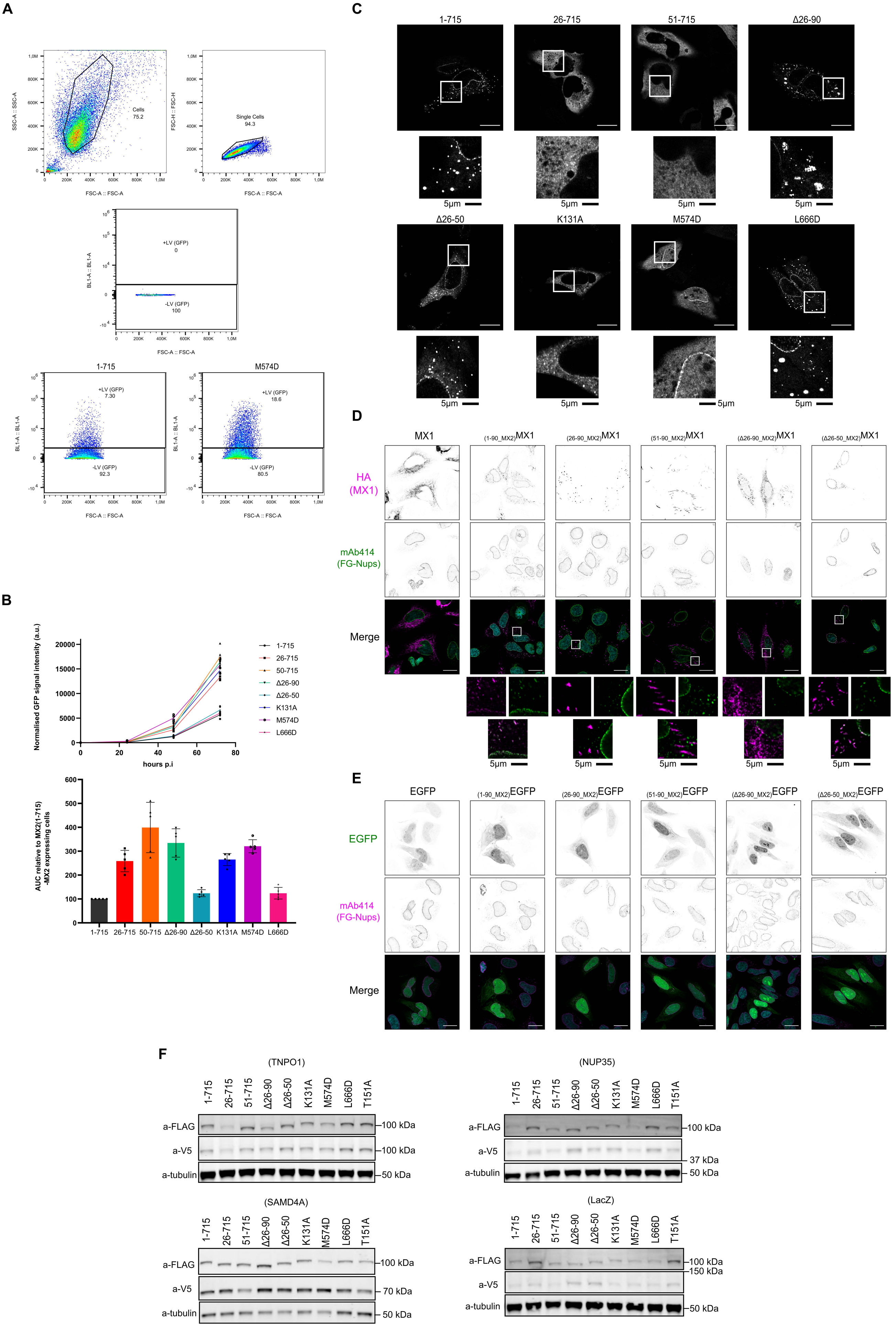

A
Cells
75.2
Single Cells
94.3
SSC-A :: SSC-A
FSC-H :: FSC-H
FSC-A :: FSC-A
BL1-A :: BL1-A
+LV (GFP)
0
-LV (GFP)
100
1-715
+LV (GFP)
7.30
-LV (GFP)
92.3
M574D
+LV (GFP)
18.6
-LV (GFP)
80.5
B
Normalised GFP signal intensity (a.u.)
hours p.i
1-715
26-715
50-715
Δ26-90
Δ26-50
K131A
M574D
L666D
AUC relative to MX2(1-715) -MX2 expressing cells
1-715
26-715
50-715
Δ26-90
Δ26-50
K131A
M574D
L666D
C
1-715
26-715
51-715
Δ26-90
Δ26-50
K131A
M574D
L666D
5μm
D
MX1
(1-90_MX2)MX1
(26-90_MX2)MX1
(51-90_MX2)MX1
(Δ26-90_MX2)MX1
(Δ26-50_MX2)MX1
HA (MX1)
mAb414 (FG-Nups)
Merge
5μm
E
EGFP
(1-90_MX2)EGFP
(26-90_MX2)EGFP
(51-90_MX2)EGFP
(Δ26-90_MX2)EGFP
(Δ26-50_MX2)EGFP
EGFP
mAb414 (FG-Nups)
Merge
F
(TNPO1)
(NUP35)
(SAMD4A)
(LacZ)
1-715
26-715
51-715
Δ26-90
Δ26-50
K131A
M574D
L666D
T151A
a-FLAG
a-V5
a-tubulin
100 kDa
150 kDa
70 kDa
37 kDa
50 kDa

### Figure S7

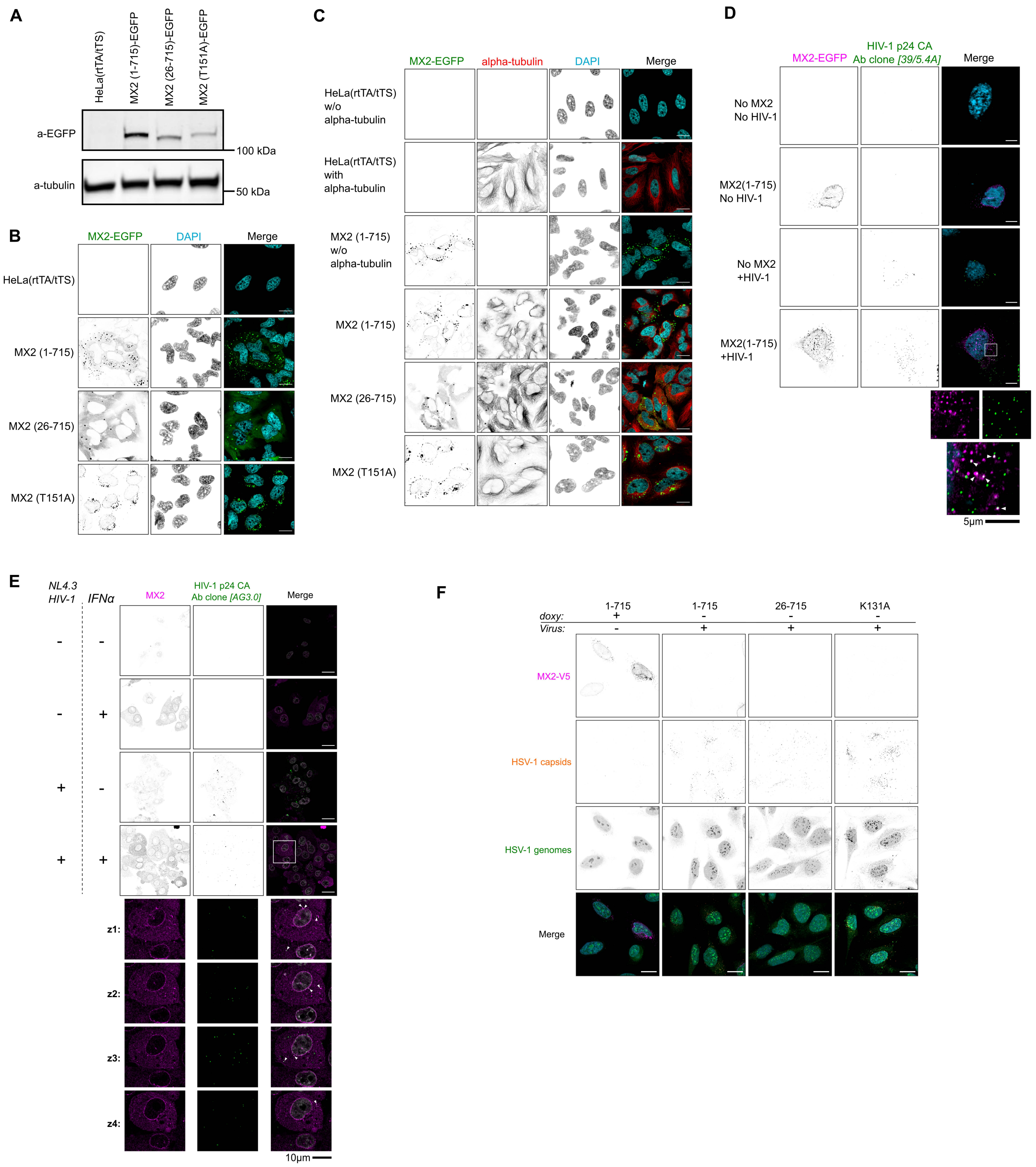

A
HeLa(rtTA/tTS)
MX2 (1-715)-EGFP
MX2 (26-715)-EGFP
MX2 (T151A)-EGFP
a-EGFP
a-tubulin
100 kDa
50 kDa
B
MX2-EGFP
DAPI
Merge
HeLa(rtTA/tTS)
MX2 (1-715)
MX2 (26-715)
MX2 (T151A)
C
MX2-EGFP
alpha-tubulin
DAPI
Merge
HeLa(rtTA/tTS) w/o alpha-tubulin
HeLa(rtTA/tTS) with alpha-tubulin
MX2 (1-715) w/o alpha-tubulin
MX2 (1-715)
MX2 (26-715)
MX2 (T151A)
D
MX2-EGFP
HIV-1 p24 CA Ab clone [39/5.4A]
Merge
No MX2 No HIV-1
MX2(1-715) No HIV-1
No MX2 +HIV-1
MX2(1-715) +HIV-1
5µm
E
NL4.3 HIV-1
IFNα
MX2
HIV-1 p24 CA Ab clone [AG3.0]
Merge
z1:
z2:
z3:
z4:
10µm
F
doxy:
Virus:
1-715
1-715
26-715
K131A
MX2-V5
HSV-1 capsids
HSV-1 genomes
Merge

### Figure S8

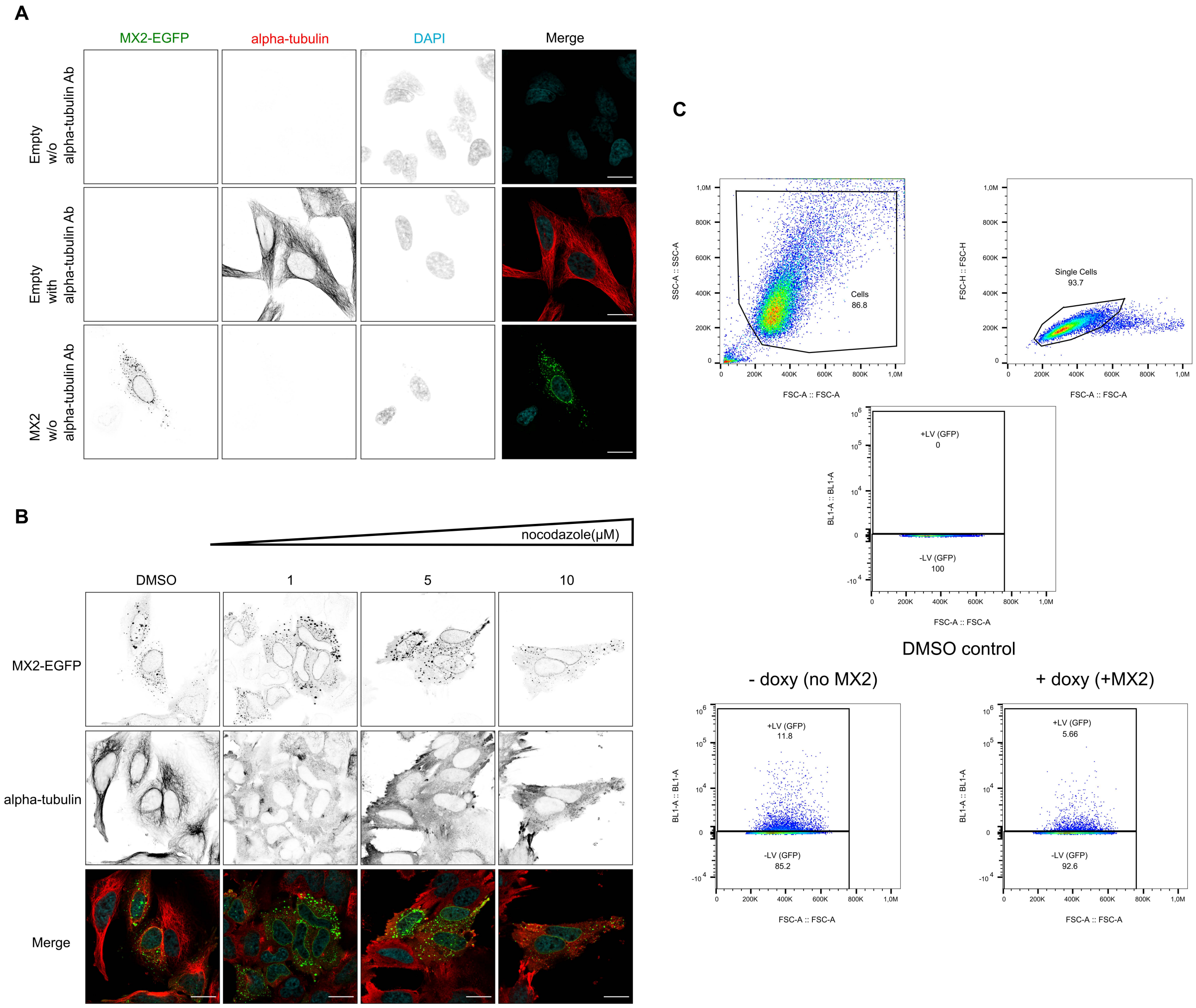

A
MX2-EGFP
alpha-tubulin
DAPI
Merge
Empty w/o alpha-tubulin Ab
Empty with alpha-tubulin Ab
MX2 w/o alpha-tubulin Ab
B
nocodazole(μM)
DMSO
1
5
10
MX2-EGFP
alpha-tubulin
Merge
C
Cells 86.8
Single Cells 93.7
+LV (GFP) 0
-LV (GFP) 100
DMSO control
- doxy (no MX2)
+ doxy (+MX2)
+LV (GFP) 11.8
-LV (GFP) 85.2
+LV (GFP) 5.66
-LV (GFP) 92.6
SSC-A :: SSC-A
FSC-H :: FSC-H
BL1-A :: BL1-A
FSC-A :: FSC-A
