## Supplementary material for "MX2 restricts HIV-1 and herpes simplex virus type 1 by forming cytoplasmic biomolecular condensates that mimic nuclear pore complexes": Link to supplementary videos

Videos for ‘’Moschonas et al. bioRxiv 2024. MX2 restricts HIV-1 and herpes simplex virus type 1 by forming cytoplasmic biomolecular condensates that mimic nuclear pore complexes’’ can be seen on (and downloaded from) [FigShare](https://figshare.com/articles/media/Movies_from_McKellar_et_al_bioRxiv_2024_Human_MX1_induces_the_cytoplasmic_sequestration_of_neo-synthesized_influenza_A_virus_vRNPs/25289647):

<https://figshare.com/articles/media/Supplementary_Videos_from_Moschonas_et_a_bioRxiv_2014_MX2_restricts_HIV-1_and_herpes_simplex_virus_type_1_by_forming_cytoplasmic_biomolecular_condensates_that_mimic_nuclear_pore_complexes/25513102>

**Supplementary videos**

**Sup.Video.1. LIVE_MX2-EGFP:** HeLa cells transiently expressing C-terminally EGFP-tagged MX2(1-715) [shown in green] imaged as single z-slice under time-lapse conditions. Single channels are shown in inverted grey and the merged images with colours. Timestamp is shown at top left. Scale bar: 10 µm.

**Sup.Video.1. LIVE_MX2-EGFP_ZoomA:** Zoom panel A of the highlighted white boxed area from Figure 3D. HeLa cells transiently expressing C-terminally EGFP-tagged MX2(1-715) [shown in green] imaged as single z-slice under time-lapse conditions. Zoomed single channels are shown in inverted grey and the merged images with colours. Timestamp is shown at top left. Scale bar: 5 µm.

**Sup.Video.1. LIVE_MX2-EGFP_ZoomB:** Zoom panel B of the highlighted white boxed area from Figure 3D. HeLa cells transiently expressing C-terminally EGFP-tagged MX2(1-715) [shown in green] imaged as single z-slice under time-lapse conditions. Zoomed single channels are shown in inverted grey and the merged images with colours. Timestamp is shown at top left. Scale bar: 5 µm.

**Sup.Video.2. LIVE_SAMD4A-EGFP + MX2-mScarlet:** HeLa cells transiently expressing C-terminally EGFP-tagged SAMD4A [shown in green] together with C-terminally mScarlet-tagged MX2(1-715) [shown in violet] imaged as single z-slice under time-lapse conditions. Single channels are shown in inverted grey and the merged images with colours. Timestamp is shown at top left. Scale bar: 10 µm.

**Sup.Video.2. LIVE_SAMD4A-EGFP + MX2-mScarlet_Zoom:** Zoom panel of the highlighted white boxed area from Figure 3E. HeLa cells transiently expressing C-terminally EGFP-tagged SAMD4A [shown in green] together with C-terminally mScarlet-tagged MX2(1-715) [shown in violet] imaged as single z-slice under time-lapse conditions. Zoomed single channels are shown in inverted grey and the merged images with colours. Timestamp is shown at top left. Scale bar: 5 µm.

**Sup.Video.3. LIVE_TNPO1-EGFP + MX2-mScarlet:** HeLa cells transiently expressing C-terminally EGFP-tagged TNPO1 [shown in green] together with C-terminally mScarlet-tagged MX2(1-715) [shown in violet] imaged as single z-slice under time-lapse conditions. Single channels are shown in inverted grey and the merged images with colours. Timestamp is shown at top left. Scale bar: 10 µm.

**Sup.Video.3. LIVE_TNPO1-EGFP + MX2-mScarlet_Zoom:** Zoom panel of the highlighted white boxed area from Figure 3E. HeLa cells transiently expressing C-terminally EGFP-tagged TNPO1 [shown in green] together with C-terminally mScarlet-tagged MX2(1-715) [shown in violet] imaged as single z-slice under time-lapse conditions. Zoomed single channels are shown in inverted grey and the merged images with colours. Timestamp is shown at top left. Scale bar: 5 µm.

**Sup.Video.4. LIVE_NUP35-EGFP + MX2-mScarlet:** HeLa cells transiently expressing C-terminally EGFP-tagged NUP35 [shown in green] together with C-terminally mScarlet-tagged MX2(1-715) [shown in violet] imaged as single z-slice under time-lapse conditions. Single channels are shown in inverted grey and the merged images with colours. Timestamp is shown at top left. Scale bar: 10 µm.

**Sup.Video.4. LIVE_NUP35-EGFP + MX2-mScarlet_Zoom:** Zoom panel of the highlighted white boxed area from Figure 3E. HeLa cells transiently expressing C-terminally EGFP-tagged NUP35 [shown in green] together with C-terminally mScarlet-tagged MX2(1-715) [shown in violet] imaged as single z-slice under time-lapse conditions. Zoomed single channels are shown in inverted grey and the merged images with colours. Timestamp is shown at top left. Scale bar: 5 µm.

**Sup.Video.5. LIVE_NUPL1-EGFP + MX2-mScarlet:** HeLa cells transiently expressing C-terminally EGFP-tagged NUPL1 [shown in green] together with C-terminally mScarlet-tagged MX2(1-715) [shown in violet] imaged as single z-slice under time-lapse conditions. Single channels are shown in inverted grey and the merged images with colours. Timestamp is shown at top left. Scale bar: 10 µm.

**Sup.Video.5. LIVE_NUPL1-EGFP + MX2-mScarlet_Zoom:** Zoom panel of the highlighted white boxed area from Figure 4E. HeLa cells transiently expressing C-terminally EGFP-tagged NUPL1 [shown in green] together with C-terminally mScarlet-tagged MX2(1-715) [shown in violet] imaged as single z-slice under time-lapse conditions. Zoomed single channels are shown in inverted grey and the merged images with colours. Timestamp is shown at top left. Scale bar: 5 µm.

**Sup.Video.6. LIVE_IPO4-EGFP + MX2-mScarlet:** HeLa cells transiently expressing C-terminally EGFP-tagged IPO4 [shown in green] together with C-terminally mScarlet-tagged MX2(1-715) [shown in violet] imaged as single z-slice under time-lapse conditions. Single channels are shown in inverted grey and the merged images with colours. Timestamp is shown at top left. Scale bar: 10 µm.

**Sup.Video.6. LIVE_IPO4-EGFP + MX2-mScarlet_Zoom:** Zoom panel of the highlighted white boxed area from Figure 4E. HeLa cells transiently expressing C-terminally EGFP-tagged IPO4 [shown in green] together with C-terminally mScarlet-tagged MX2(1-715) [shown in violet] imaged as single z-slice under time-lapse conditions. Zoomed single channels are shown in inverted grey and the merged images with colours. Timestamp is shown at top left. Scale bar: 5 µm.

**Sup.Video.7. LIVE_LSM14A-EGFP + MX2-mScarlet:** HeLa cells transiently expressing C-terminally EGFP-tagged LSM14A [shown in green] together with C-terminally mScarlet-tagged MX2(1-715) [shown in violet] imaged as single z-slice under time-lapse conditions. Single channels are shown in inverted grey and the merged images with colours. Timestamp is shown at top left. Scale bar: 10 µm.

**Sup.Video.7. LIVE_LSM14A-EGFP + MX2-mScarlet_Zoom:** Zoom panel of the highlighted white boxed area from Figure S5B. HeLa cells transiently expressing C-terminally EGFP-tagged LSM14A [shown in green] together with C-terminally mScarlet-tagged MX2(1-715) [shown in violet] imaged as single z-slice under time-lapse conditions. Zoomed single channels are shown in inverted grey and the merged images with colours. Timestamp is shown at top left. Scale bar: 5 µm.

**Sup.Video.8. LIVE_SAMD4B-EGFP + MX2-mScarlet:** HeLa cells transiently expressing C-terminally EGFP-tagged SAMD4B [shown in green] together with C-terminally mScarlet-tagged MX2(1-715) [shown in violet] imaged as single z-slice under time-lapse conditions. Single channels are shown in inverted grey and the merged images with colours. Timestamp is shown at top left. Scale bar: 10 µm.

**Sup.Video.8. LIVE_SAMD4B-EGFP + MX2-mScarlet_Zoom:** Zoom panel of the highlighted white boxed area from Figure S5F. HeLa cells transiently expressing C-terminally EGFP-tagged SAMD4B [shown in green] together with C-terminally mScarlet-tagged MX2(1-715) [shown in violet] imaged as single z-slice under time-lapse conditions. Zoomed single channels are shown in inverted grey and the merged images with colours. Timestamp is shown at top left. Scale bar: 5 µm.

**Sup.Video.9. LIVE_FUBP3-EGFP + MX2-mScarlet:** HeLa cells transiently expressing C-terminally EGFP-tagged FUBP3 [shown in green] together with C-terminally mScarlet-tagged MX2(1-715) [shown in violet] imaged as single z-slice under time-lapse conditions. Single channels are shown in inverted grey and the merged images with colours. Timestamp is shown at top left. Scale bar: 10 µm.

**Sup.Video.9. LIVE_FUBP3-EGFP + MX2-mScarlet_Zoom:** Zoom panel of the highlighted white boxed area from Figure S5F. HeLa cells transiently expressing C-terminally EGFP-tagged FUBP3 [shown in green] together with C-terminally mScarlet-tagged MX2(1-715) [shown in violet] imaged as single z-slice under time-lapse conditions. Zoomed single channels are shown in inverted grey and the merged images with colours. Timestamp is shown at top left. Scale bar: 5 µm.

**Sup.Video.10. LIVE_MX2(D26-90):** HeLa cells transiently expressing C-terminally EGFP-tagged MX2(Δ26-90) [shown in inverted grey] imaged as single z-slice under time-lapse conditions. Timestamp is shown at top left. Scale bar: 10 µm.

**Sup.Video.11. LIVE_MX2(1-715):** HeLa cells transiently expressing C-terminally EGFP-tagged MX2(1-715) [shown in inverted grey] imaged as single z-slice under time-lapse conditions. Timestamp is shown at top left. Scale bar: 10 µm.

**Sup.Video.12. LIVE_MX2(26-715):** HeLa cells transiently expressing C-terminally EGFP-tagged MX2(26-715) [shown in inverted grey] imaged as single z-slice under time-lapse conditions. Timestamp is shown at top left. Scale bar: 10 µm.

**Sup.Video.13. LIVE_MX2(51-715):** HeLa cells transiently expressing C-terminally EGFP-tagged MX2(51-715) [shown in inverted grey] imaged as single z-slice under time-lapse conditions. Timestamp is shown at top left. Scale bar: 10 µm.

**Sup.Video.14. LIVE_MX2(****D26-50):** HeLa cells transiently expressing C-terminally EGFP-tagged MX2(Δ26-50) [shown in inverted grey] imaged as single z-slice under time-lapse conditions. Timestamp is shown at top left. Scale bar: 10 µm.

**Sup.Video.15. LIVE_MX2(K131A):** HeLa cells transiently expressing C-terminally EGFP-tagged MX2(K131A) [shown in inverted grey] imaged as single z-slice under time-lapse conditions. Timestamp is shown at top left. Scale bar: 10 µm.

**Sup.Video.16. LIVE_MX2(M574D):** HeLa cells transiently expressing C-terminally EGFP-tagged MX2(M574D) [shown in inverted grey] imaged as single z-slice under time-lapse conditions. Timestamp is shown at top left. Scale bar: 10 µm.

**Sup.Video.17. LIVE_MX2(L666D):** HeLa cells transiently expressing C-terminally EGFP-tagged MX2(L666D) [shown in inverted grey] imaged as single z-slice under time-lapse conditions. Timestamp is shown at top left. Scale bar: 10 µm.

**Sup.Video.18. LIVE_MX2(T151A):** HeLa cells transiently expressing C-terminally EGFP-tagged MX2(T151A) [shown in inverted grey] imaged as single z-slice under time-lapse conditions. Timestamp is shown at top left. Scale bar: 10 µm.
